# Identifying and overcoming the sampling challenges in relative binding free energy calculations of a model protein:protein complex

**DOI:** 10.1101/2023.03.07.530278

**Authors:** Ivy Zhang, Dominic A. Rufa, Iván Pulido, Michael M. Henry, Laura E. Rosen, Kevin Hauser, Sukrit Singh, John D. Chodera

## Abstract

Relative alchemical binding free energy calculations are routinely used in drug discovery projects to optimize the affinity of small molecules for their drug targets. Alchemical methods can also be used to estimate the impact of amino acid mutations on protein:protein binding affinities, but these calculations can involve sampling challenges due to the complex networks of protein and water interactions frequently present in protein:protein interfaces. We investigate these challenges by extending a GPU-accelerated open-source relative free energy calculation package (Perses) to predict the impact of amino acid mutations on protein:protein binding. Using the well-characterized model system barnase:barstar, we describe analyses for identifying and characterizing sampling problems in protein:protein relative free energy calculations. We find that mutations with sampling problems often involve charge-changes, and inadequate sampling can be attributed to slow degrees of freedom that are mutation-specific. We also explore the accuracy and efficiency of current state-of-the-art approaches—alchemical replica exchange and alchemical replica exchange with solute tempering—for overcoming relevant sampling problems. By employing sufficiently long simulations, we achieve accurate predictions (RMSE 1.61, 95% CI: [1.12, 2.11] kcal/mol), with 86% of estimates within 1 kcal/mol of the experimentally-determined relative binding free energies and 100% of predictions correctly classifying the sign of the changes in binding free energies. Ultimately, we provide a model workflow for applying protein mutation free energy calculations to protein:protein complexes, and importantly, catalog the sampling challenges associated with these types of alchemical transformations. Our free open-source package (Perses) is based on OpenMM and available at https://github.com/choderalab/perses.

## INTRODUCTION

### Predicting the impact of amino acid mutations on protein:protein binding has important applications

Protein:protein interactions (PPIs) underlie fundamental biological processes, such as transcriptional regulation (e.g., p53:MDM2 [1]), signal transduction (e.g., GRB2-EGFR [2]), and membrane fusion (e.g., SARS-CoV-2 RBD:ACE2 [3]). As protein:protein interactions are driven by binding events, changes in protein:protein binding affinity often have functional impact. Even a single amino acid substitution can significantly alter binding and function, which can give rise to disease [4], impact the fitness of a pathogen [5], or alter the activity of monoclonal antibody drugs [6]. Thus, quantifying the impact of an amino acid mutation on protein:protein binding is highly useful for predicting the functional implication of a mutation, as it can provide mechanistic understanding of disease-associated genetic variants [7, 8] and facilitate the design of biologic drugs such as monoclonal antibodies [9, 10].

### Alchemical free energy calculations represent an accurate and generalizable approach for estimating mutational impact on PPIs

There are many experimental and computational approaches for quantifying the impact of a mutation on protein:protein binding. Experimental approaches, while highly accurate and generally considered “ground truth,” can be resource intensive, often requiring significant amounts of human labor time and costly reagents and instruments [11–14]. Computational methods, which can circumvent these resource challenges, serve as a complementary approach that can be combined with experimental methods to enable more efficient acquisition of high confidence data. Examples include computationally inexpensive methods (e.g., MM/PBSA and MM/GBSA [15, 16], machine learning (ML) models [17, 18], and Rosetta-based methods [19, 20]), as well as computationally expensive methods such as alchemical free energy calculations. Several studies have compared the tradeoffs between using computationally inexpensive methods and alchemical free energy calculations to predict the impact of a point mutation on binding [21–25]. While computationally inexpensive methods can be accurate for certain systems [21, 22], these methods often fail to account for key biophysical phenomena (i.e., conformational heterogeneity, explicit solvent interactions, multiple protonation states), and therefore perform worse on systems which require modeling of these properties [23–25]. Despite their increased computational cost, alchemical free energy calculations can account for these biophysical phenomena using rigorous statistical mechanics, so they tend to demonstrate better accuracy than the cheaper methods and are generalizable to any PPI with available structural data [24–27]. Ultimately, the optimal choice in approach will depend on the scientific goal and the computational resources available. However, given their accuracy and generalizability, as well as rapid advancements in graphics processing units (GPUs) that have made it feasible to carry out these calculations in reasonable timeframes [28–31], alchemical free energy calculations represent a highly promising approach for predicting mutational effect on protein:protein binding.

### Relative alchemical binding free energy calculations aim to predict the impact of a mutation on the free energy of binding (ΔΔG_binding_)

While there are numerous types of alchemical free energy calculations [32], relative alchemical binding free energy (RBFE) calculations estimate the relative binding free energies (ΔΔG_binding_) between two chemically similar complexes, e.g., protein:protein complexes that differ by an amino acid mutation (Figure 1A). In protein mutation RBFE calculations [21, 23–25], the wild-type (WT) residue is transformed into the mutant residue through molecular dynamics simulations of alchemical (non-physical) intermediate states bridging the WT and mutant states (Figure 2A). This alchemical transformation is performed in two phases, complex and apo, which correspond to the mutating protein in the presence and absence of a protein binding partner, respectively (Figure 1A). The change in free energy associated with each phase (ΔG_phase_) is estimated, and the difference in the ΔG_phase_s gives an estimate of the impact of the mutation on the binding free energy, ΔΔG_binding_ (Figure 1A).

**Figure 1.**
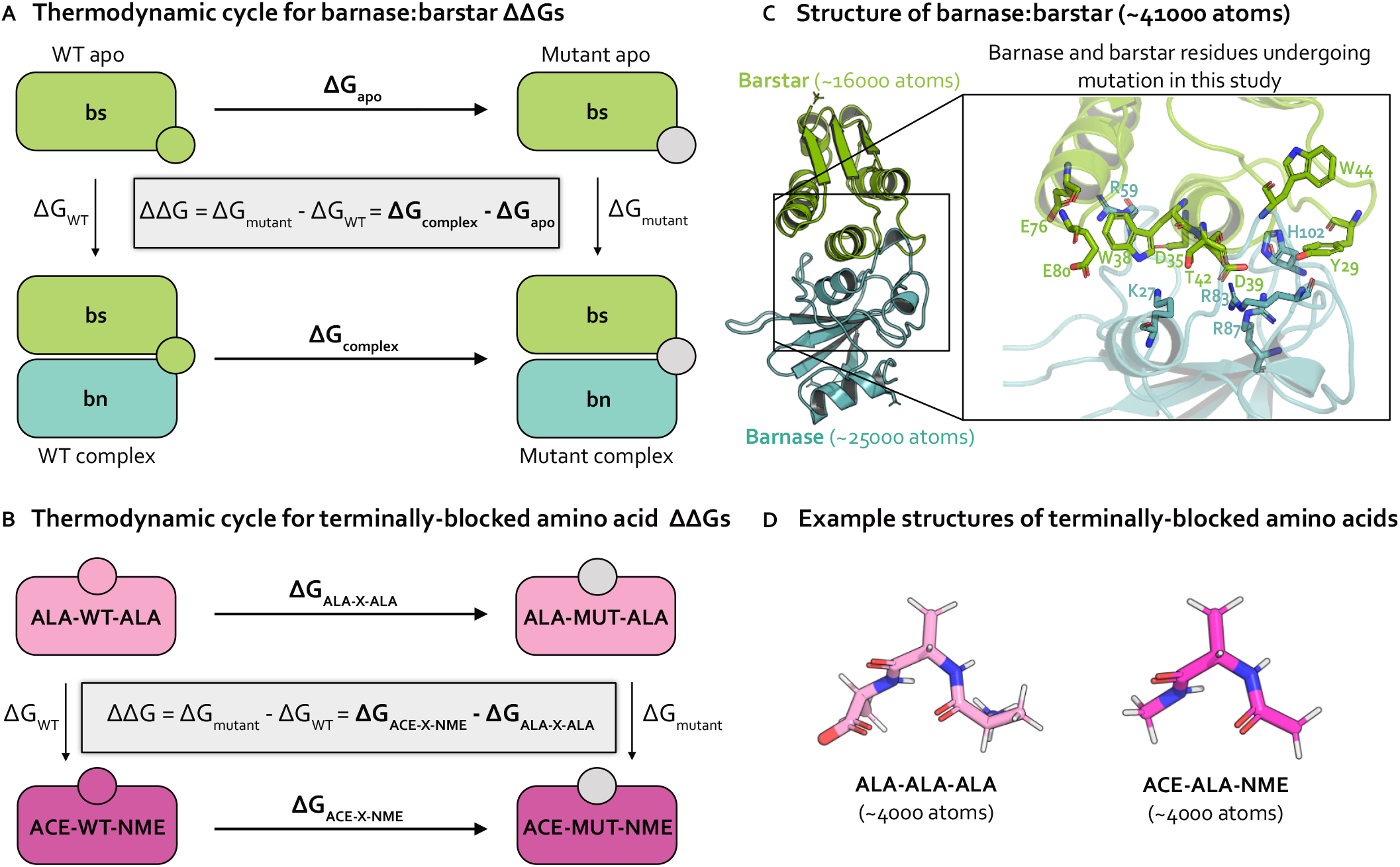
Relative free energy calculations predict the impact of single point mutations using thermodynamic cycles that each involve transformations in two environments. **(A)** Thermodynamic cycle representing how the relative binding free energy (ΔΔG_binding_) can be computed for a protein mutation in the barnase:barstar complex. By cycle closure, the ΔΔG equation shown inside the thermodynamic cycle can be recovered. In practice, it is easier to compute the horizontal legs (ΔG_apo_ and ΔG_complex_, shown in bold) [33], which involve transforming a WT residue (green circle) into a mutant residue (gray circle). The free energy differences for each phase (apo and complex) are subtracted to compute the ΔΔG_binding_. **(B)** Thermodynamic cycle representing how the relative free energy (ΔΔG) can be computed for a protein mutation between two phases of terminally-blocked amino acids. The horizontal legs (ΔG_ALA-X-ALA_ and ΔG_ACE-X-NME_, shown in bold) are simulated, which involve transforming a WT residue (magenta or pink circle) into a mutant residue (gray circle). The free energy differences for each phase (ACE-X-NME and ALA-X-ALA) are subtracted to compute the ΔΔG. **(C)** Structural model of barnase:barstar (PDB ID: 1BRS) with barstar shown in green and barnase shown in blue. Barstar and barnase contain ∼16000 and ∼25000 atoms, respectively (including hydrogens and solvent). Zoomed-in view of the barnase:barstar interface shows the 13 residues undergoing mutation in this study (all of which are interfacial) as sticks. Nitrogen atoms shown in blue and oxygen atoms are shown in red. **(D)** Example structural models of terminally-blocked amino acids: ALA-X-ALA and ACE-X-NME (where X is ALA) shown in pink and magenta, respectively. Each terminally-blocked amino acid contains ∼4000 atoms (including hydrogens and solvent). Nitrogen atoms are depicted in blue, oxygen atoms in red, and hydrogen atoms in white.

### Achieving sufficient sampling of protein and water conformations is particularly challenging for RBFE calculations applied to protein:protein interactions

During the last couple of decades, RBFE calculations have become increasingly widely used in drug discovery projects for predicting the effects of small molecule modifications on protein:small molecule binding [31, 34– 40]. In comparison to small molecule transformations, application of RBFE calculations to protein mutations has been relatively limited, though recent studies have demonstrated that these methods can accurately predict mutational impact on protein:small molecule binding [21–23, 41] and protein:protein binding [24, 25, 42–46] for a number of biologically-relevant complexes.

One reason for their lack of widespread use stems from the size of protein:protein complexes, which can frequently involve approximately double the number of atoms as are present in protein:small molecule complexes, making them more computationally expensive to simulate. However, the bigger hurdle has been the sampling challenges associated with alchemical transformations in protein:protein complexes [24, 47]. RBFE calculations require drawing decorrelated samples from the configurational probability distributions at each alchemical state [48], a nontrivial task for protein:protein complexes because the energy landscapes often contain many minima which can give rise to slow degrees of freedom. Slow degrees of freedom are more prevalent in protein:protein complexes because protein:protein interfaces are generally broader than protein:small molecule interfaces and typically involve complex protein and water interaction networks [4, 49]. Upon mutation, extensive reorganization of the mutating residue along with its closely-packed neighborhood of interfacial protein residues and waters may be required before one can draw decorrelated samples.

### Pinpointing slow degrees of freedom can help address the sampling problems in protein:protein RBFE calculations, but existing approaches are not automated

Determining the slow degrees of freedom causing sampling problems in alchemical free energy calculations is useful because it may enable improved sampling via methods that accelerate known slow degrees of freedom (e.g., metadynamics [50, 51], umbrella sampling [52], adaptive biasing force [53]). Moreover, identifying slow degrees of freedom helps enumerate the common challenges associated with alchemical transformations and the limitations of existing methods, which will facilitate the improvement of existing methods or the design of new ones. However, pinpointing the slow degrees of freedom in protein:protein interfaces can be challenging because it typically involves careful manual inspection of simulation trajectories [33, 54, 55], a process which requires biophysical intuition and can be tedious even for experienced practitioners. Moreover, the manual inspection approach is not scalable as examining the trajectories for tens or hundreds of mutations would be extremely impractical.

Here, we investigate sampling problems associated with protein mutation relative free energy calculations using (1) terminally-blocked amino acids, a small and simple test system relatively free of interfacial complexities and (2) barnase:barstar, a well-studied protein:protein complex. We augment an existing open-source relative free energy calculation package (Perses [56], https://github.com/choderalab/perses) to carry out these calculations and describe experiments and automated analyses that identify likely causes of sampling problems. We find that sampling challenges are more likely to occur for charge-changing mutations and can be attributed to mutation-dependent slow degrees of freedom. We also compare the accuracy and efficiency of state-of-the-art enhanced sampling approaches—alchemical replica exchange (AREX) [31, 57, 58] and alchemical replica exchange with solute tempering (AREST) [59, 60]—for overcoming the sampling challenges. We find that given sufficient simulation time, our predictions are accurate with respect to experiment (RMSE 1.61, 95% CI: [1.12, 2.11] kcal/mol), with 86% of predictions lying within 1 kcal/mol of experimental ΔΔ*G*_binding_s and 100% of predictions having the correct sign.

## THEORY

We perform alchemical free energy calculations using two state-of-the-art enhanced sampling approaches: (1) alchemical replica exchange (AREX) [31, 57, 58], the current recommended approach based on best practices [27] and (2) alchemical replica exchange with solute tempering (AREST) [59, 60], a sampling scheme which builds upon AREX by increasing the temperature of a region around the mutating residue and has been shown to improve sampling over AREX for some transformations [60–62]. Here, we give a brief overview of the salient aspects of each method, as well as the general approach we take to alchemical free energy calculations for protein mutations. The alchemical approach is implemented in an open-source package (Perses [56], available at https://github.com/choderalab/perses). Complete simulation details can be found in the **Detailed Methods**.

### Alchemical transformation

Alchemical free energy calculations aim to sample a set of alchemical states which are defined such that two endstates of interest are bridged by alchemical intermediate states with modified Hamiltonians. The Hamiltonians are modified such that the nonbonded interactions (and potentially valence terms) of the WT and mutant residues are gradually transformed between interacting and non-interacting. The first alchemical state, called the WT endstate, typically involves the WT residue fully interacting with its environment and the mutant residue completely non-interacting. The last alchemical state, known as the mutant endstate, involves the mutant residue fully interacting and the WT residue non-interacting. A series of intermediate alchemical states bridging the endstates is defined such that WT and mutant residues are partially interacting with their environments to varying extents. Taken together, these alchemical states form the alchemical transformation (Figure 2A). Note that amino acids with backbone cycles (such as proline) require a modified version of this approach to avoid the noninteracting residue influencing the conformational distribution of fully interacting residues [63], but the mutations in this study do not involve this type of amino acid.

To characterize an alchemical transformation, we need to specify both *which* interactions will be alchemically modified during the transformation and *how* we will modify them. We first identify the alchemical interactions by defining an atom mapping, which exploits the partial similarity in WT and mutant topologies by pairing up the WT and mutant atoms that will share coordinates. The atom mapping is then used to classify each atom into an “atom class”: “unique old” atoms are unmapped atoms that are only present in the WT residue, “unique new” atoms are unmapped atoms that are only present in the mutant residue, “core” atoms are mapped atoms that are shared between the WT and mutant residues (and include atoms in the residues immediately preceding and following the mutating residue), and “environment” atoms are mapped atoms that are shared between the topologies but lie outside of the core atoms. Interactions involving “unique old”, “unique new” and “core” atoms are considered alchemical interactions. Since increasing the number of alchemical interactions also increases the thermodynamic length (i.e., the distance between alchemical states) [64], an atom mapping should be defined to maximize the number of atoms mapped between the two residues, which minimizes thermodynamic length. An optimal mapping finds a balance between minimizing thermodynamic length and taking advantage of the built-in enhanced sidechain sampling that occurs when the unmapped atoms are non-interacting (i.e., the nonbonded interactions are scaled to zero, so the sidechains can more easily sample alternate rotameric states). To this end, we chose an atom mapping that maps all atoms between the WT and mutant residues up to and including the beta carbon (but not including beta hydrogens). Because we constrain bonds to hydrogen, we un-map any hydrogen atoms whose bond lengths would change between WT and mutant.

To specify how the energies should be modified during the alchemical transformation, we introduce an alchemical parameter *λ* ∈ [0, 1] into the potential energy function *U* (*x*), forming the alchemical potential energy function *U* (*x*; *λ*). The alchemical potential *U* (*x*; *λ*) is typically evaluated at a different *λ* value for each alchemical state. For the WT endstate, *U* (*x*; *λ* = 0) is identical to the unmodified WT potential with the addition of the standard valence terms (and not the nonbonded interactions) of the unique new atoms (sometimes called “dummy” atoms). For the mutant endstate, *U* (*x*; *λ* = 1) is identical to the unmodified mutant potential, but with unique old atoms as noninteracting dummy atoms that only retain their valence terms. For the alchemical intermediate states (*λ* ∈ (0, 1)), *U* (*x*; *λ*) is a modified potential where interactions involving the WT and mutant residues are scaled to varying extents. The set of *λ* values sampled in the alchemical transformation is termed the “alchemical protocol”. To define the alchemical protocol for this study, we selected evenly spaced *λ* values from a simple linear function (Supplementary Figure 1A). The number of *λ* values used for each calculation was chosen such that the neighboring alchemical states have good phase space overlap (for more details, see **Detailed Methods**), which permits robust estimation of free energy differences between the fully-interacting WT and mutant endstates [65, 66].

**Figure 2.**
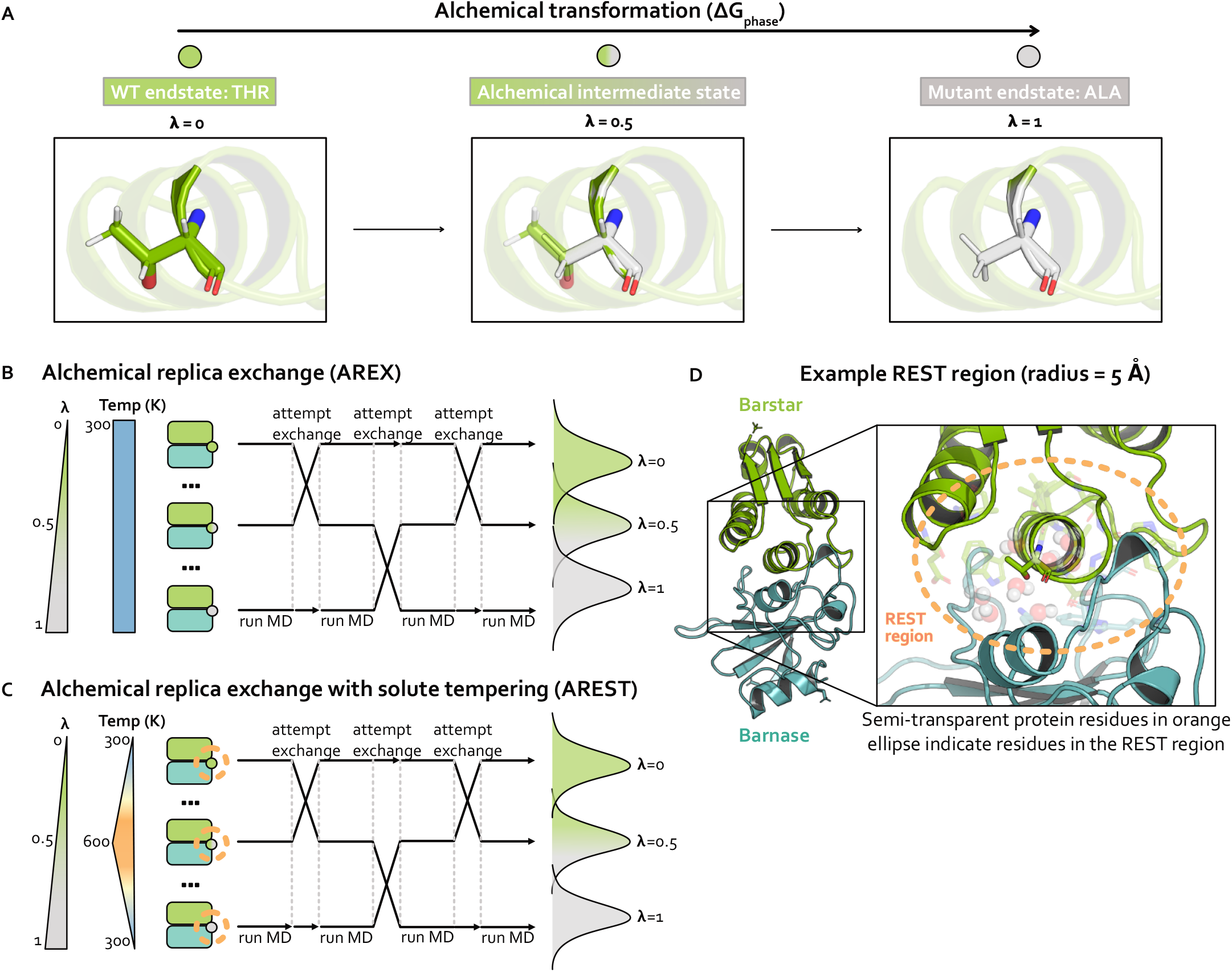
Strategies for sampling an alchemical transformation: Alchemical replica exchange (AREX) and alchemical replica exchange with solute tempering (AREST). AREST modifies AREX by introducing local heating around the alchemical region at intermediate alchemical states. **(A)** Schematic representing an alchemical transformation (with one alchemical intermediate state) for one simulation phase. The WT (*λ* = 0, green) endstate contains a fully-interacting threonine residue and the mutant (*λ* = 1, gray) endstate contains a fully-interacting alanine residue. The alchemical intermediate (*λ* = 0.5, green-gray gradient) state contains partially interacting threonine and alanine residues. Nitrogen atoms are shown in blue and oxygen atoms are shown in red. **(B)** Schematic representing alchemical replica exchange (AREX), sometimes called Hamiltonian replica exchange among alchemical states, which utilizes multiple replicas (in this schematic, three replicas) to explore alchemical states that bridge the WT (*λ* = 0, green circle) and mutant (*λ* = 1, gray circle) fully interacting states. The temperature remains constant at 300 K for all alchemical states. Representative configurational distributions for each alchemical state are shown (on the right) to be overlapping for neighboring states, which is a requirement for accurate ΔΔG estimates. **(C)** Schematic representing alchemical replica exchange with solute tempering (AREST), which elevates the effective temperature for a small region (i.e., the REST region) to further enhance sampling. The REST region is shown as an orange, dashed circle. The effective temperature of the REST region reaches a maximum at 600 K at *λ* = 0.5. Representative configurational distributions for each alchemical state are shown (on the right) with less overlap than in AREX (panel B), because increasing the effective temperature usually causes increased thermodynamic length. **(D)** Structural model of barnase:barstar with barstar shown in green and barnase shown in blue. Zoomed-in view highlights an example REST region (orange, dashed circle) for residue T42 in barstar. T42 and neighboring residues (within 5 Å of T42) are shown as sticks and neighboring waters (also within 5 Å of T42) are shown as spheres.

Using the alchemical parameter *λ*, we define the potential energy functions for electrostatic and steric interactions. We compute the alchemically modified electrostatics interaction energy according to the Particle Mesh Ewald (PME) method [67] with linearly interpolated charges. For the direct space electrostatics contribution:

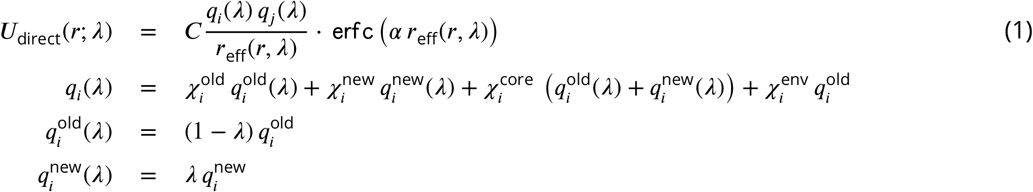

where *α* is an internal PME parameter (calculated based on the PME error tolerance and the cutoff distance, with dimension 1/length), *C* is the Coulomb constant (with dimension energy/length^2^), *q*_*i*_(*λ*) and *q*_*j*_ (*λ*) are the functions for computing the potentially alchemically modified charges of atoms *i* and *j* (with dimension of charge), and 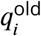 and 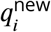 are the charges of atom *i* in the old topology and new topology, respectively.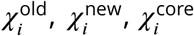, and 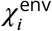 are indicator functions denoting whether atom *i* belongs in the unique old, unique new, core, and environment atom classes, respectively.

*r*_eff_(*r, λ*) denotes the *effective interaction distance* used for computing the interaction energy, and depends on both the actual inter-particle separation *r* and the alchemical parameter *λ*. To avoid singularities in the computation of electrostatics (and sterics) energies, we use a softcore approach that involves “lifting” certain inter-atomic distances into the “4th dimension”, inspired by work of Pomès [68]:

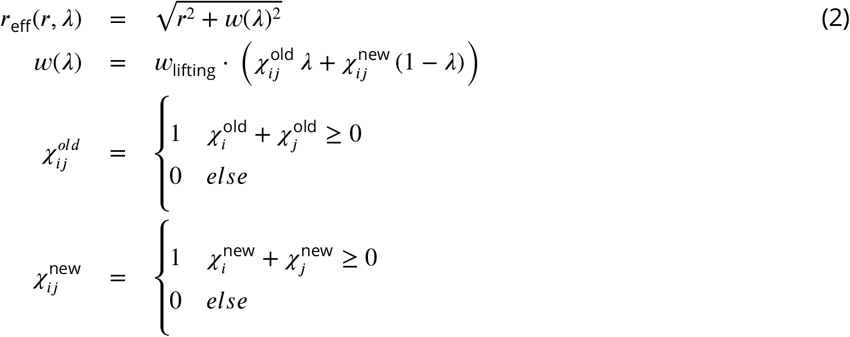

where *r* is the distance between atoms *i* and *j, w*(*λ*) is the function for computing the lifting distance, and *w*_lifting_ is the maximal lifting distance, which was selected to minimize the number of alchemical states needed to produce robust free energy estimates while maintaining good overlap among neighboring alchemical states. 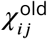 is an indicator function that assumes the value of unity only when at least one of the atoms (*i* and *j*) belongs in the unique old atom class (and zero otherwise), and 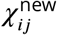 is an indicator function indicating whether at least one of the atoms (*i* and *j*) belongs in the unique new atom class. For more details on this approach, see **Detailed Methods**.

We compute the PME reciprocal space and self-energy contributions using the default energy functions in OpenMM [29], but with linearly interpolated charges, where interpolation was performed in the same manner as was done for the direct space.

We compute the alchemically modified sterics interaction energy as a standard Lennard-Jones 12-6 potential [69–71] with linearly interpolated *σ* and *ϵ* and “lifted” interaction distances to create a softcore potential:

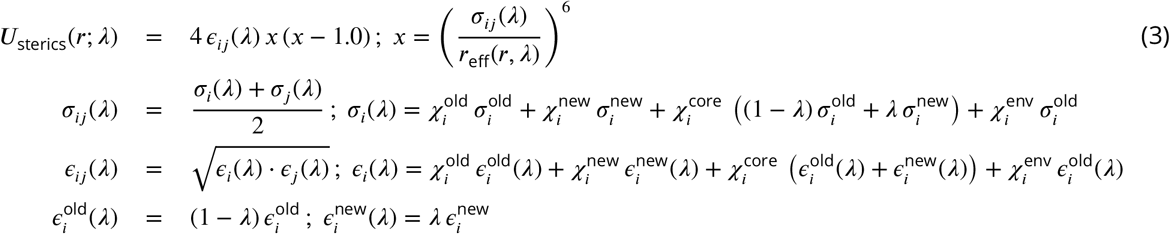

Here, *σ*_*ij*_ (*λ*) is the function for computing the potentially alchemically modified distance at which the interaction energy crosses zero for atoms *i* and *j*. 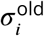 and 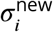 are the distances at which the energy equals zero for atom *i* in the old topology and new topology, respectively. *ϵ*_*ij*_ (*λ*) is the function for computing the potentially alchemically modified interaction strength for atoms *i* and *j*. 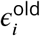 and 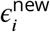 are the interaction strengths for atom *i* in the old topology and new topology, respectively.

For charge-changing mutations, we ensure the system remains electrostatically neutral by transforming a water molecule in the WT system into a sodium or chloride ion in the mutant system. The ΔΔGs for charge-changing mutations in terminally-blocked amino acids are internally consistent, indicating that in the absence of sampling problems, our counterion scheme enables robust estimation of free energies (Figure 3A). Further details on this implementation can be found in **Detailed Methods**.

We estimate free energy differences using the Multistate Bennett Acceptance Ratio (MBAR), which is an asymptotically unbiased estimator that, in the large sample limit, often has lower variance compared to other commonly used estimators [66].

### Alchemical replica exchange (AREX)

Alchemical free energy calculations must sample from a chain of alchemical intermediate states bridging the two endstates of interest (Figure 2A). Because the introduction or deletion of bulky residues can often frustrate sampling within alchemical states in which these residues are almost fully interacting, alchemical free energy calculations often use replica exchange simulations to help reduce correlation times and overcome sampling challenges. Replica exchange enhances sampling by allowing each replica to visit multiple alchemical states, including those states which may help more rapidly decorrelate slow degrees of freedom because of their modified Hamiltonians [31, 57, 58]. Here, we refer to this approach as alchemical replica exchange (AREX).

AREX can be thought of as a Markov Chain Monte Carlo (MCMC) algorithm that aims to generate equilibrium samples from a family of ***K*** probability densities corresponding to the ***K*** alchemical states:

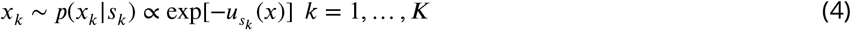

where *x*_*k*_ is a configuration drawn from state *k, s*_*k*_ is the *k*^th^ state, and 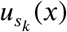 is the potential energy of sample *x* at state *s*_*k*_.

To generate equilibrium samples, AREX utilizes weakly-coupled replicas (copies of the system of interest), where the number of replicas is typically equal to the number of alchemical states ***K***. AREX employs a Gibbs sampling framework where in each iteration *n*, the positions 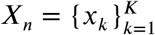 of all ***K*** replicas are first updated with molecular dynamics simulations, yielding *X*_*n*+1_, and then the permutation set of alchemical state indices 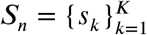 associated with the corresponding positions are updated based on the updated positions *X*_*n*+1_, yielding *S*_*n*+1_:

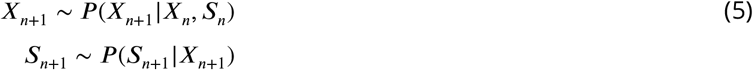

In sufficiently long simulations, the resulting samples (***X***_*n*_, ***S***_*n*_) are distributed with respect to the joint probability density ***P*** (***X***_*n*_, ***S***_*n*_) such that

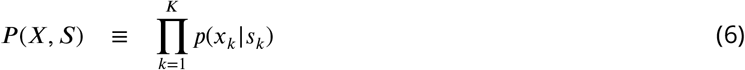

The algorithm updates the state indices by exchanging the alchemical state labels for pairs of replicas according to a Metropolis criterion that compares the energies of the two replicas considered for swapping. The replica swap acceptance rate will depend on how well the alchemical states overlap, i.e., the thermodynamic length between states (Figure 2B). Numerous methods can be used to update the states ***S***, including attempting exchanges only between replicas visiting neighboring thermodynamic states (where state overlap is highest). Here, we use a simple strategy that attempts to draw an independent permutation ***S***_*n*+1_ given configuration ***X***_*n*+1_ by attempting many swaps of pairs of alchemical state indices, which has been shown to enhance mixing and reduce correlation times [72].

AREX involves running many cycles of molecular dynamics followed by exchange attempts with the goal of ensuring all replicas ultimately perform a random walk through all alchemical states (Figure 2B). If the exchanges are accepted at a sufficiently high rate over the course of the simulation, we expect to observe improved sampling because configurational correlation times associated with the alchemical region are likely decreased for alchemical states with partially interacting residues.

### Alchemical replica exchange with solute tempering (AREST)

AREST is AREX with an added layer of sophistication that aims to enhance sampling to a greater extent than AREX. AREST involves running AREX with a REST (replica exchange solute tempering [59]) region, a user-defined set of atoms for which the effective temperature is increased in alchemical intermediate states (Figure 2C-D). Therefore, in AREST, the alchemical states do not solely differ by the extent to which the WT and mutant residues are interacting with their environment, they also differ by the effective temperature of the REST region. Although introducing differences in effective temperature will increase the thermodynamic length between alchemical states (Figure 2C), the goal is to decrease the correlation time of the slowest degrees of freedom sufficiently to compensate for the decrease in state overlap, yielding more decorrelated samples in the same amount of total simulation time.

To incorporate REST into AREX, we classify each bond, angle, torsion, and nonbonded interaction as “REST” (all atoms in the interaction are part of the REST region), “inter” (at least one atom is part of the REST region and at least one atom is not), or “non-REST” (none of the atoms are part of the REST region) based on an initial conformation. Each interaction energy is multiplied by a scale factor depending on the REST class. Therefore, total potential energy is defined as:

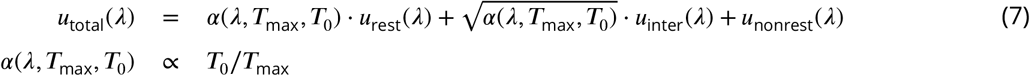

where ***T***_0_ is the temperature of the desired distribution and ***T***_max_ is the user-selected maximum effective temperature. The function we use to define the REST scale factor, *α*(*λ*, ***T***_max_, ***T***_0_), is shown in Supplementary Figure 1B. Note that when *λ* = 0 or 1, *α*(*λ*, ***T***_max_, ***T***_0_) = 1 to ensure the endstates are unscaled.

## METHODS AND SYSTEMS

For complete details on system setup and simulation parameters, see **Detailed Methods**.

### Barnase:barstar

Our investigation primarily focuses on the bacterial protein:protein complex barnase:barstar. The interaction of barnase, an extracellular ribonuclease, with its intracellular inhibitor, barstar, regulates RNA degradation in bacterial cells with a binding free energy of -19 kcal/mol [73]. Solvated barnase:barstar simulation models contain only ∼41,000 atoms (including hydrogens and solvent), making it a computationally tractable system for studying sampling challenges in high-affinity protein:protein interfaces (Figure 1C, see **Detailed Methods** for system preparation details). Barnase:barstar has been well-studied both computationally [25, 47, 74] and experimentally [73, 75–77]). The barnase:barstar mutations considered in this work come from Schreiber et al. [73], who used stopped-flow measurements to derive experimental relative binding free energies (ΔΔG_binding_s) for 14 single amino acid substitutions across 13 residue positions in either barnase or barstar. The ΔΔG_binding_s for this set of mutations span an unusually large dynamic range (7.8 kcal/mol) with a statistical error of 0.1 kcal/mol, and involves a diverse set of amino acids, making it particularly useful for assessing quantitative predictive models. All mutations occur within or are in close proximity to the barnase:barstar interface, which is a complex network of interactions dominated by electrostatic interactions and coordinated by buried waters (Figure 1C) [73]. Therefore, the mutations tend to disrupt numerous interfacial interactions, potentially requiring significant conformational and water reorganization to achieve equilibrium, which may give rise to sampling challenges.

### Terminally-blocked amino acids

As a control to the complexity of barnase:barstar, we also study terminally-blocked amino acids, which lack the complex interaction networks of barnase:barstar and have relatively few solute degrees of freedom. Specifically, we introduce mutations in small, solvated amino acids in two different environments: either terminally-blocked with ACE and NME caps at the N- and C-termini, respectively (ACE-X-NME), or terminally blocked with ALA residues with natural zwitterionic termini (ALA-X-ALA) (Figure 1D). The terminally-blocked mutation set consists of the same amino acid mutations as in the barnase:barstar mutation set, but contains only 10 total mutations (instead of 14 for barnase:barstar) as some of the barnase:barstar mutations involve the same amino acid transformation at different residue positions. By introducing the same mutations into the terminally-blocked amino acids, we separate the sampling challenges present in the barnase:barstar interface from the common challenges associated with alchemical free energy calculations.

To obtain relative free energies (ΔΔGs), we estimate the free energy differences (ΔGs) for two phases. For barnase:barstar, we are interested in the ΔΔG_binding_, so the two simulation phases are complex and apo (Figure 1A). For terminally-blocked amino acids, there is no notion of binding, so the two phases are: ACE-X-NME and ALA-X-ALA (Figure 1B).

## RESULTS

In the following sections, we investigate the sampling problems associated with applying relative free energy calculations to predict the impact of mutations in a model protein:protein complex. We establish an open-source workflow which consists of: (1) identifying mutations that are potentially plagued by sampling problems, (2) determining the slow degrees of freedom responsible for poor sampling, and (3) exploring state-of-the-art approaches for improving sampling. We first apply our workflow to a simple test system, terminally-blocked amino acids. We then focus the rest of the work on sampling challenges at the complex protein:protein interface of barnase:barstar, benchmarking to experimentally-determined binding free energies. While this study mainly focuses on analyzing the sampling problems in one protein:protein complex, past studies have performed similar types of analyses on other protein:ligand and protein:protein complexes [24, 54], so we expect our approach to be generalizable to other systems.

### 1 Mutations at protein:protein interfaces can be challenging for alchemical replica exchange relative free energy calculations, likely due to inadequate sampling in complex phase simulations

#### 1.1 The relative free energy differences (ΔΔGs) for terminally-blocked amino acid mutations are internally consistent, well converged, and relatively absent of sampling problems

We first establish that running alchemical replica exchange (AREX) with our alchemical approach (e.g., alchemical protocol, atom mapping, softcore and counterion approaches, etc.) is free of sampling and convergence issues when the mutation is not located in the context of a complex network of protein interactions. We estimated the ΔΔGs of 10 terminally blocked amino acid mutations between two environments: ACE-X-NME and ALA-X-ALA, where X is an amino acid. For each mutation, we ran simulations in both the forward (A→B) and reverse (B→A) directions, where A corresponds to the amino acid in the WT barnase:barstar crystal structure (PDB ID: 1BRS). A mutation was considered internally consistent if the ΔΔG for the forward mutation (A→B) was within statistical error of the −ΔΔG for the reverse mutation (B→A). We found that with 5 ns/replica AREX simulations, all of the mutations are internally consistent and the forward ΔΔGs match the negative of the reverse ΔΔGs with high accuracy (root mean square error (RMSE): 0.21, 95% confidence interval (CI): [0.12, 0.28] kcal/mol, Figure 3A).

We next confirm that our calculations lack replica mixing bottlenecks and convergence issues. We first checked that there are no replica mixing bottlenecks for any of the mutations, indicating that the alchemical states are spaced such that they have reasonable overlap (Supplementary Figure 2). We next determined the extent to which the free energy difference of mutating WT→Mutant in one phase (ΔG) is converged because a converged ΔG indicates that the simulation has likely sampled all relevant degrees of freedom sufficiently. We assessed convergence by monitoring the changes in ΔG as a function of simulation time, which we call a “ΔG time series”. A ΔG time series was considered converged if, within the last five nanoseconds, it appeared flat with a close-to-zero slope (0 ± 0.1 kcal/mol/ns) (Supplementary Figure 3A). A ΔG time series was considered not converged if the magnitude of the slope of the last 5 ns was not within statistical uncertainty of 0 kcal/mol/ns. We found that for all 10 mutations in both phases and in both the forward and reverse directions, the slope of the ΔG time series is within statistical uncertainty of 0 kcal/mol/ns (Figure 3C-D, Supplementary Figure 3B-C), suggesting that the calculations are converged and relatively free of sampling problems.

Finally, we verify that 5 ns/replica AREX simulations thoroughly sample the slowest degrees of freedom for terminally blocked amino acid mutations [78]. We monitored the *ϕ* and *ψ* angles for the ACE-X-NME phase of two representative mutations with significant sampling problems in barnase:barstar, A2T (ALA to THR at residue 2) and R2A (ARG to ALA at residue 2) (see Section 2). If the *ϕ* and *ψ* degrees of freedom are thoroughly sampled, the time series should rapidly decorrelate. We quantify the extent to which each time series is hindered by slow correlation times by estimating its statistical inefficiency, *g* = 2*τ* + 1, which is proportional to the autocorrelation of the time series *τ* [79, 80]. Since the sampling interval for the time series is 0.1 ns, if the statistical inefficiency (*g*) is close to 0.1 ns, the samples are completely decorrelated, and the larger the value of *g*, the more correlated the samples are [80]. We observed that for both representative mutations, ACE-X-NME phase simulations thoroughly sample both angles with *g* close to 0.1 ns (Supplementary Figure 4), providing further support that the terminally-blocked amino acid calculations are converged and have minimal sampling problems.

These results demonstrate that in the absence of significant sampling problems, running AREX with our alchemical approach can provide reliable estimates of relative free energy differences (ΔΔGs), indicating that we can use this approach to explore the challenges associated with applying RBFE calculations to interfacial residues in the barnase:barstar protein:protein complex.

#### 1.2 Several barnase:barstar mutation predictions show poor accuracy due to slow convergence of the complex phase free energy difference (ΔG_complex_), suggesting the presence of sampling problems

We assess the performance of AREX on predicting barnase:barstar relative binding free energies (ΔΔG_binding_s). We ran 10 ns/replica AREX simulations for the 14 mutations in the barnase:barstar mutation set in both the forward (i.e., mutations start from crystal structure residue) and reverse directions, resulting in a total of 28 ΔΔG_binding_ predictions. We first compared the predicted versus experimental ΔΔG_binding_s and considered a mutation to be significantly discrepant if the 95% CIs of its predicted and experimental ΔΔG_binding_s were not within 1 kcal/mol of each other. We observed relatively poor agreement (RMSE: 2.49, 95% CI: [1.32, 3.74] kcal/-mol) with 7% (2/28) of the predictions having the wrong sign and 21% (6/28) of the predictions considered significantly discrepant (Figure 6B, Supplementary Figure 6A). Moreover, when we compared the forward and negative of the reverse ΔΔG_binding_s for each mutation, we found that 21% (3/14) of mutations have poor internal consistency (i.e., the forward and negative reverse ΔΔG_binding_s are not within statistical error of each other) (Figure 6A or Figure 3B). We refer to the following mutations, all with poor accuracy with respect to experiment, as significantly discrepant mutations: A42T, R87A, D35A, H102A, A29Y, Q83R (Supplementary Figure 6A). A subset of these mutations (A42T, R87A, Q83R) also has poor internal consistency (Figure 6A, Supplementary Table 1).

We next demonstrate that all mutations with significant discrepancy have sufficiently overlapping alchemical states and some have slow ΔG_complex_ convergence. To assess state overlap, we checked for sufficient replica mixing in both phases of simulation and found that the replicas mix well, indicating that the ΔΔG_binding_ discrepancies are not a result of poor overlap of alchemical states (Supplementary Figure 5). We next determined whether the free energy difference of mutating WT→Mutant in each phase (ΔG) has converged by checking whether the slope of the last 5 ns of the ΔG time series (i.e., ΔG as a function of simulation time) is within statistical uncertainty of zero (0 ± 0.1 kcal/mol/ns). We found that 67% (4/6) of the significantly discrepant mutations (A42T, R87A, H102A, and Q83R) have ΔG_complex_s with slow convergence, suggesting that the corresponding simulations may contain significant sampling problems (Figure 3E-G). For the remaining 33% (2/6) of significantly discrepant mutations (D35A and A29Y), the ΔG_complex_s do converge within 10 ns (Figure 3G), which indicates that they may have minimal sampling problems, though it is possible that the slowest degrees of freedom in these simulations have correlation times longer than 10 ns and therefore have not yet been sampled.

**Figure 3.**
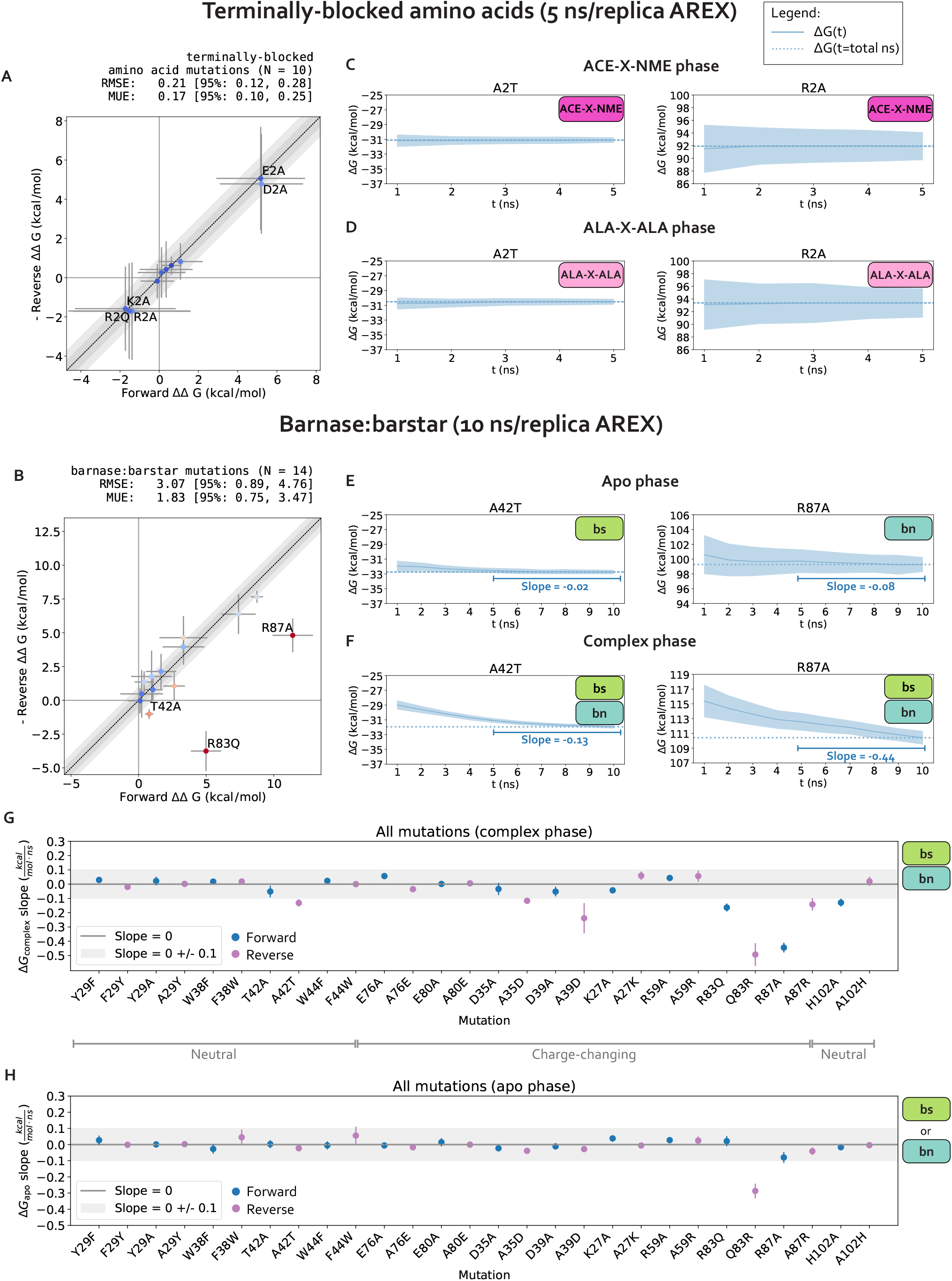
The relative free energy difference (ΔΔG) predictions for small terminally-blocked amino acid mutations are internally consistent and show good convergence, but several of the predictions for interfacial barnase:barstar mutations show poor internal consistency due to slow convergence of complex phase free energy differences (ΔG_complex_s). **(A)** (Negative of the) Reverse versus forward ΔΔGs for each terminally-blocked amino acid mutation computed using alchemical replica exchange (AREX) simulations (number of states = 12 and 24 for neutral and charge mutations, respectively, and simulation time = 5 ns/replica for each phase). Data points are labeled if the mutation involves a charge-change (to emphasize that our counterion introduction scheme works well in the absence of sampling **(A)** The y = x (black dotted) line represents zero discrepancy between forward and (negative of the) reverse ΔΔGs, the dark gray shaded region represents 0.5 kcal/mol discrepancy, and the light gray region represents 1 kcal/mol discrepancy. Data points are colored by how far they are from zero discrepancy (dark blue and red indicate close to and far from zero, respectively). Error bars represent two standard deviations and were computed by bootstrapping the decorrelated reduced potential matrices 200 times. Root mean square error (RMSE) and mean unsigned error (MUE) are shown with 95% confidence intervals obtained from bootstrapping the ΔΔGs 1000 times. **(B)** (Negative of the) Reverse versus forward ΔΔGs for each barnase:barstar mutation computed using alchemical replica exchange (AREX) simulations (number of states = 24 and 36 for neutral and charge mutations, respectively and simulation time = 10 ns/replica for each phase). Data points are labeled if the forward and (negative of the) reverse ΔΔG_binding_s are not within statistical error of each other (i.e., neither the forward nor the negative reverse ΔΔG_binding_ is within 1 kcal/mol of the 95% CI for the other ΔΔG_binding_). For more details on the plot and error bars, refer to the caption for panel A. **(C)** Free energy difference (ΔG) time series for representative mutations A2T (left) and R2A (right) in the ACE-X-NME phase. Alchemical replica exchange simulations were performed with number of states = 12 and 24 for A2T and R2A, respectively and the simulation time was 5 ns/replica. Dashed line indicates the ΔG at t = 5 ns. Shaded region represents ± two standard deviations, which were computed by bootstrapping the decorrelated reduced potential matrices 200 times. **(D)** Same as (B), but for the ALA-X-ALA phase, instead of the ACE-X-NME phase. **(E)** Free energy difference (ΔG) time series for the apo phase of representative mutations with sampling problems: A42T (left) and R87A (right). Alchemical replica exchange simulations were performed with number of states = 24 and 36 for A42T and R87A, respectively and the simulation time was 10 ns/replica. Dashed line indicates the ΔG at t = 10 ns. For details on the error bars, refer to the caption for panel C. **(F)** Same as (E), but for the complex phase instead of the apo phase. **(G)** Slopes of the last 5 ns of the ΔG_complex_ time series for each barnase:barstar mutation are shown as blue (forward mutations) and purple (reverse mutations) circles. ΔG_complex_ time series were generated from complex phase AREX simulations (number of states = 24 and 36 for neutral and charge mutations, respectively, and simulation time = 10 ns/replica). Error bars represent 2 standard deviations and were computed using the SciPy `linregress` function. Slopes within error of the shaded gray region (0 ± 0.1 kcal/mol/ns) are close to zero and are therefore considered “flat.” **(H)** Same as (G), but for apo phase barnase or barstar mutations instead of complex phase barnase:barstar mutations.

Finally, we show that sampling problems often occur in complex phase simulations, especially for charge-changing mutations. We extended the convergence analysis to all 28 barnase:barstar mutations and observed that most of the mutations (27/28) have ΔG_apo_s that converge within 10 ns/replica (Figure 3H). However, 25% (7/28) of mutations have ΔG_complex_s that do not converge, some of which are mutations that have predicted ΔΔG_binding_s close to experiment (R83Q, A39D, A35D) and should therefore be considered problematic mutations (Figure 3G). More importantly, this analysis indicates that convergence may be more difficult to achieve in the complex phase simulations, likely because of difficulties in sampling. Furthermore, out of the seven mutations with poor ΔG_complex_ convergence, six of the mutations are charge-changing, suggesting that sampling may be more challenging for charge-changing mutations (Figure 3G).

In summary, we identified several barnase:barstar mutations with predicted ΔΔG_binding_s that exhibit poor accuracy and described an approach for identifying mutations that potentially have significant sampling challenges. We found that 32% (9/28) of the mutations have discrepant ΔΔG_binding_s or slow ΔG_complex_ convergence with 10 ns/replica AREX simulations. Moreover, while sampling problems are absent for terminally-blocked amino acid mutations, they are likely present in the complex phase for several barnase:barstar mutations, most of which involve charge-changes. In the next section, we attempt to identify the slow degrees of freedom causing sampling challenges.

### 2 Poor complex phase sampling can occur due to mutation-dependent slow protein or water degrees of freedom

We choose two significantly discrepant mutations for deeper analysis of potential sampling challenges, A42T and R87A, each of which also has poor internal consistency (Supplementary Figure 6A, Figure 3B). These mutations encompass distinct types of transformations: A42T is a reverse mutation that involves a neutral, small to medium amino acid change (ALA to THR) and R87A is a forward mutation that involves a charge-changing, large to small transformation (ARG to ALA). Both mutations have slowly converging complex phase free energy differences (ΔG_complex_s) which are likely a result of sampling problems (Figure 3F). In this section, we confirm the presence of sampling problems in A42T and R87A complex phase simulations and identify slow degrees of freedom likely causing poor sampling.

#### 2.1 Sampling challenges can be caused by hindered protein conformational dynamics

We first hypothesize that slow ΔG_complex_ convergence can be attributed to poor sampling of slow protein backbone or side chain motion in the A42T and R87A complex phase simulations. To test this hypothesis, we ran 10 ns/replica complex phase AREX simulations where we imposed restraints on the heavy-atom coordinates to eliminate protein motion as a source of slow degrees of freedom, and compared the ΔG_complex_ time series with and without restraints for each mutation. Because these restraints significantly reduce protein motion, if the slow ΔG_complex_ convergence is caused by slow protein motions that are insufficiently sampled, the restraints should eliminate the sampling problem and the ΔG_complex_ time series should converge immediately. The A42T ΔG_complex_ time series with restraints converges rapidly within 10 ns, lacking the downward trend that is present in the ΔG_complex_ time series without restraints and indicating that the A42T complex phase simulation has a protein sampling problem (Figure 4A). However, for R87A, the ΔG_complex_ time series with restraints is within error of the time series without restraints, exhibiting the same downward trend as the unrestrained time series, which suggests the lack of convergence in the R87A ΔG_complex_ is not solely caused by protein sampling problems (Figure 4D). Although this analysis can help determine the presence (or absence) of protein sampling problems, it does not identify the specific slow degrees of freedom that are likely causing sampling problems.

#### 2.2 Poor sampling can specifically be attributed to individual sidechain torsions, interfacial contacts, or nearby waters

We next determine the specific degrees of freedom which may be responsible for slow ΔG_complex_ convergence by identifying conformational degrees of freedom that are tightly coupled to *∂U* /*∂λ* [54]. We are particularly interested in mutations with slowly-varying *∂U* /*∂λ* (i.e., highly correlated and with large statistical inefficiency, *g*), because slowly-varying *∂U* /*∂λ*s indicate slow ΔG_complex_ convergence. We monitored hundreds of protein and water degrees of freedom near the protein:protein interface over time because we observed that slow convergence is more common for ΔG_complex_ than ΔG_apo_ (Figure 3G-H). Specifically, we monitored the following degrees of freedom: backbone and rotameric torsion transitions near the interface, residue contacts within or between binding partners, and waters near the alchemical residue. We then computed the Pearson correlation coefficient (PCC) between *∂U* /*∂λ* and each degree of freedom, averaging over all replicas. For mutations with slowly-varying *∂U* /*∂λ*, the most highly coupled (largest magnitude PCC) degree of freedom is likely implicated in slow ΔG_complex_ convergence. We emphasize that the slow degrees of freedom discussed hereafter are relatively slow (in the context of the degrees of freedom we analyzed and in the timescales of our simulations), and that they are not necessarily the globally slowest degrees of freedom for each alchemical transformation.

For A42T, which has slowly-varying *∂U* /*∂λ* (*g* = 6.4 ns, Figure 5), the degrees of freedom with the largest magnitude PCCs are the *χ*_1_ angle of T42 (PCC: -0.63, 95% CI: [-0.68, -0.56], Figure 5) and the distance between interface barstar residues T42 and E76 (PCC: 0.61, 95% CI: [0.53, 0.66], Figure 5). The time series of a typical replica show that both degrees of freedom are highly correlated with *∂U* /*∂λ* and slowly sample two metastable states during the 50 ns replica trajectory (Figure 4B-C). The relatively slow sampling of sidechain rotamer and interface contact metastable states (correlation time: 9.2 ns for T42 *χ*_1_ and 9.5 ns for T42-E76, Figure 4B-C) likely explains the slow convergence of the A42T ΔG_complex_ time series in Figure 3F. Water sampling does not seem to play a significant role in causing poor ΔG_complex_ convergence for A42T, as the waters near T42 are only weakly correlated to *∂U* /*∂λ* (PCC: 0.30, 95% CI: [0.21, 0.37], Figure 5).

**Figure 4.**
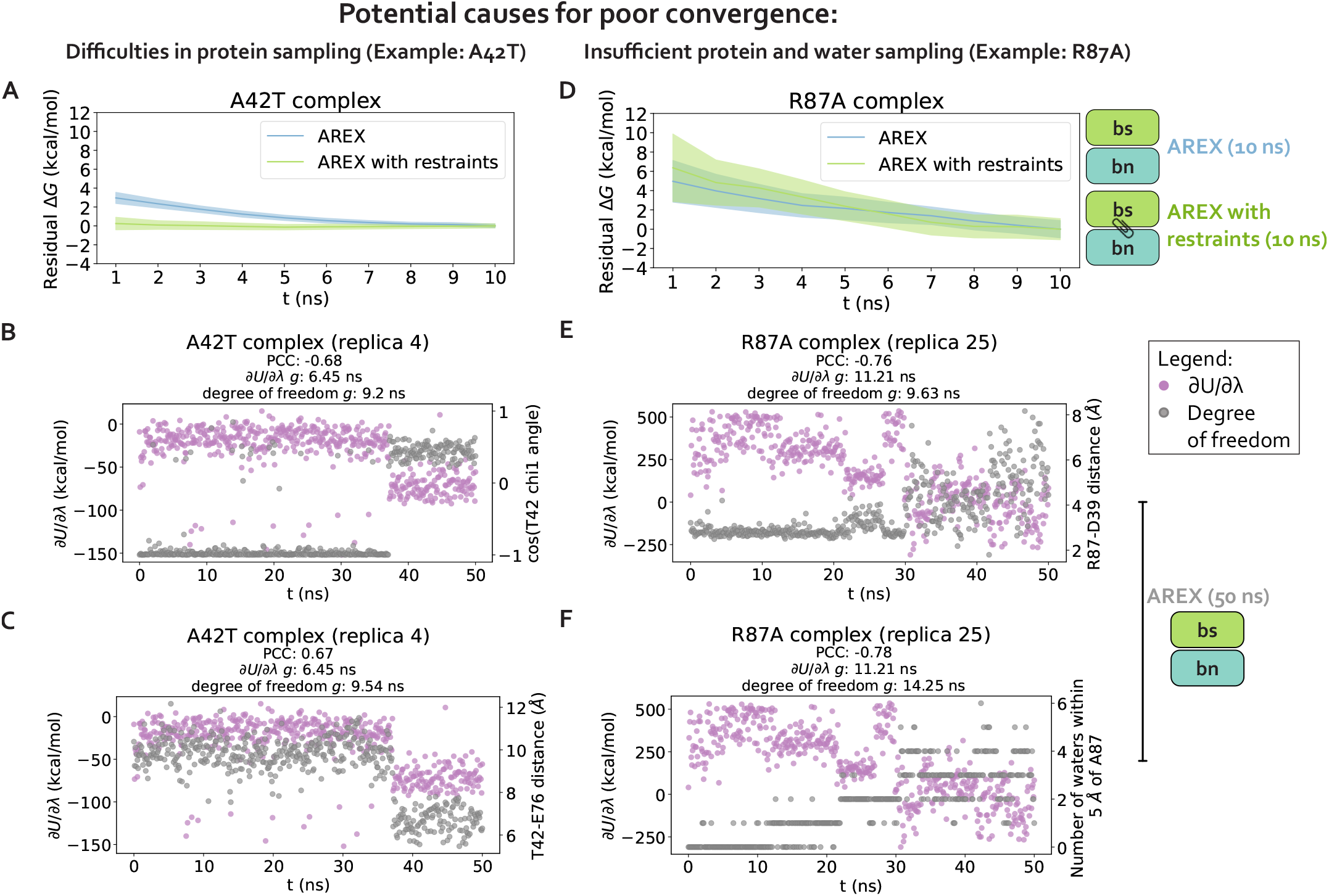
Complex phase convergence problems can arise due to insufficient sampling of protein and water degrees of freedom, e.g., a sidechain rotamer and intra-barstar contact for A42T and an inter-chain contact and neighboring waters for R87A. **(A)** Residual complex phase free energy difference (ΔG) time series for AREX simulations of A42T (number of states = 24 and simulation time = 10 ns/replica), where the residual ΔG is computed as ΔG(*t*) − ΔG(*t* = 10ns). Blue curve represents the time series for the AREX simulation without restraints and green curve represents the time series for the AREX simulation with heavy atoms restraints (force constant = 50 kcal/molÅ^2^). Shaded regions represent ± two standard deviations, which were computed by bootstrapping the decorrelated reduced potential matrices 200 times. **(B)** Time series for *∂U* /*∂λ* (left y-axis, purple) and *χ*_1_ angle for residue T42 (right y-axis, gray) for a representative replica (replica 4) of the A42T complex phase AREX simulation (number of states = 24, simulation time = 50 ns/replica). PCC indicates Pearson correlation coefficient and *g* indicates statistical inefficiency, which is proportional to the correlation time. *g* = 0.1 ns indicates very thorough sampling (because the sampling interval is 0.1 ns) and large values of *g* indicate poor sampling. **(C)** Time series for *∂U* /*∂λ* (left y-axis, purple) and T42-E76 distance (right y-axis, gray) for a representative replica (replica 4) of the A42T complex phase AREX simulation (number of states = 24, simulation time = 50 ns/replica). **(D)** Same as (A), but for R87A instead of A42T (number of states = 36 and simulation time = 10 ns/replica) and using a force constant of 75 kcal/molÅ^2^ instead of 50 kcal/molÅ^2^. **(E)** Time series for *∂U* /*∂λ* (left y-axis, purple) and R87-D39 distance (right y-axis, gray) for a representative replica (replica 25) of the R87A complex phase AREX simulation (number of states = 36, simulation time = 50 ns/replica). **(F)** Time series for *∂U* /*∂λ* (left y-axis, purple) and number of waters within 5 Å of A87 (right y-axis, gray) for a representative replica (replica 25) of the R87A complex phase AREX simulation (number of states = 36, simulation time = 50 ns/replica).

For R87A, which also has slowly-varying *∂U* /*∂λ* (*g* = 32.1 ns, Figure 5), the degree of freedom with the largest magnitude PCC is the number of waters near A87 (PCC -0.74, 95% CI: [-0.78, -0.68], Figure 5). The correlation is also particularly high for the distance between interface residues R87 (barnase) and D39 (barstar) (PCC: -0.73, 95% CI: [-0.76, -0.69], Figure 5). The time series of a representative replica shows that both degrees of freedom are highly correlated with *∂U* /*∂λ* (Figure 4E-F). Both degrees of freedom also have long correlation times (9.6 ns for R87-D39 and 14.3 ns for neighboring waters) and slow equilibration times (evidenced by the upward trend in both degree of freedom time series), which suggests the slow convergence of R87A ΔG_complex_ is likely explained by slowness in R87-D39 and nearby waters (Figure 4E-F).

We next attempted to identify general trends in the slow degrees of freedom across all barnase:barstar mutations and found that there is no common degree of freedom (or category of degrees of freedom) that is implicated in all complex phase sampling problems (Figure 5). We observed that backbone torsions consistently show PCCs less than 0.5 in magnitude (which is smaller than the PCCs of the other degrees of freedom), indicating that backbone torsions are unlikely to be the primary cause of sampling problems. The other four categories—sidechain torsions, intra interface contacts, inter interface contacts, and neighboring waters—each have many high correlation values (magnitude of PCC greater than 0.5), but no single category explains the majority of sampling problems. Therefore, the slowest degrees of freedom are highly variable depending on the mutation (Figure 5).

Finally, we observed that for complex phase simulations, mutations involving charge-changes show slower convergence than charge-preserving transformations. 83% (10/12) of neutral mutations have *∂U* /*∂λ* time series with *g* < 1, whereas 100% (16/16) of charge-changing mutations have *g* > 1 (Figure 5). We emphasize that the slow convergence of charge-changing mutations predominantly occurs in the complex (and not apo) phase (Figure 3G-H), indicating that introduction of a counterion to accommodate charge-changes does not significantly contribute to slow convergence. Instead, the sampling difficulties for charge-changing mutations likely emerge as a result of the strong network of electrostatic interactions at the barnase:barstar interface (Figure 1C).

One limitation of this work is that we studied only one protein:protein complex, and it is possible that other types of sampling problems are present in other protein:protein complexes. From our focused experiments, we cannot extrapolate how common the barnase:barstar sampling issues are for other protein:protein complexes, though it seems likely that the issues observed here are sufficiently fundamental in origin to be present in other complexes. It is worth remarking that the uniquely strong electrostatic nature of the barnase:barstar interface may exacerbate sampling challenges compared to other PPIs with less electrostatically-driven binding. The barnase:barstar interface involves 14 hydrogen bonds, more than the average protein:protein complex [75]. Of the 14 hydrogen bonds, most involve at least one charged residue, which is also atypical for protein:protein complexes [75]. Further work will be necessary to determine the extent to which the sampling problems observed in barnase:barstar are similar to those in other protein:protein systems and identify other mechanisms by which sampling problems could manifest.

Another caveat of this work is that the degrees of freedom explored in this analysis are not exhaustive; other, more complex collective variables (e.g., identified by time-lagged independent component analysis (TICA) [81, 82]) may correlate with *∂U* /*∂λ* even more highly than those explored here. Nevertheless, our scan of simple degrees of freedom reveals specific slow degrees of freedom (sidechain torsions, interfacial contacts, or nearby waters) likely implicated in slow ΔG_complex_ convergence. Moreover, we found that the degrees of freedom causing poor sampling are highly dependent on the mutation. This analysis serves as an example approach for diagnosing sampling problems in other protein:protein complexes. In the next section, we explore approaches for ameliorating the sampling challenges.

**Figure 5.**
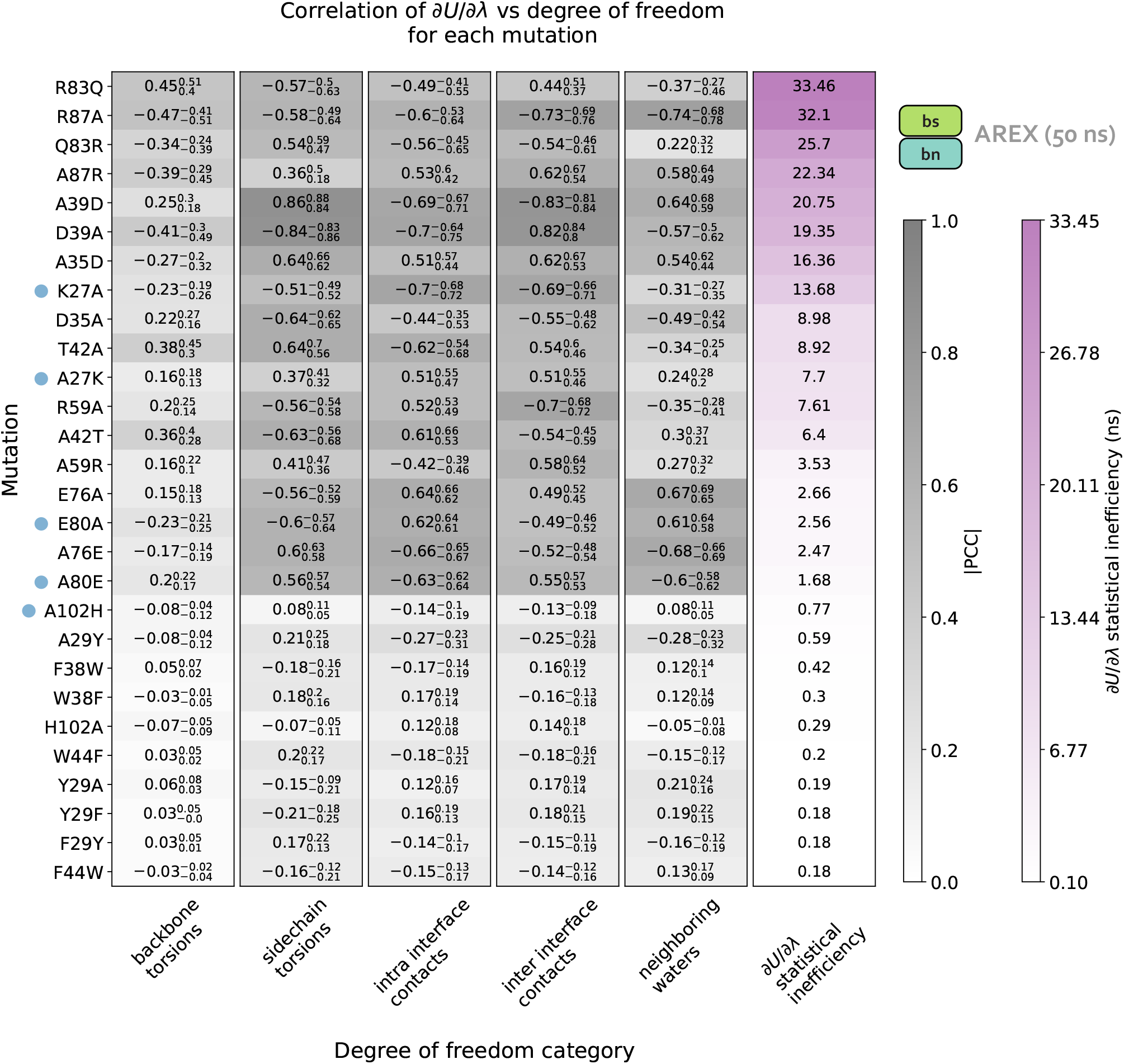
Charge-changing mutations demonstrate worse complex phase sampling than neutral mutations and the slowest degrees of freedom responsible for poor sampling are highly variable depending on the mutation. Data in this plot was generated from 50 ns/replica complex phase AREX simulations. Each row of the heatmap corresponds to a mutation and each of the first five columns corresponds to a degree of freedom category: backbone torsions, sidechain torsions, intra-interface contacts, inter-interface contacts, and neighboring waters. Each category contains a set of degrees of freedom, i.e., the backbone torsions category contains the *ϕ* and *ψ* angles for all interface residues, sidechain torsions contains the *χ*_1_, *χ*_2_, *χ*_3_, and *χ*_4_ angles for all interface residues (if the angle is present for the residue), intra-interface contacts contains pairs of interface residues that are within the same chain, inter-interface contacts contains pairs of interface residues that span different chains, and neighboring waters involves monitoring the number of waters within 5 Å of the mutating residue. Each heatmap value (in the first five columns) is the maximum of (the absolute value of) the Pearson correlation coefficients (PCCs) between *∂U* /*∂λ* and each of the degrees of freedom in the corresponding category for the corresponding mutation. For example, the top left value of the heatmap indicates that for R83Q, the backbone torsion with maximum correlation to *∂U* /*∂λ* has a PCC of 0.45. The background colors for the PCC values are different shades of gray, with darker grays indicating values closer to 1. The subscript and superscript values associated with each PCC represent the 95% confidence interval. Each heatmap value in the last column corresponds to the statistical inefficiency of *∂U* /*∂λ* across all replica trajectories for the corresponding mutation. Statistical inefficiency is proportional to the correlation time, where a value of 0.1 ns indicates very thorough sampling (because the sampling interval is 0.1 ns) and large values indicate poor sampling. Statistical inefficiency values are colored different shades of purple, with darker colors indicating larger values. The rows of the heatmap are ordered from highest to lowest by the statistical inefficiency of *∂U* /*∂λ* across all replicas. Blue dots indicate mutations for which the degree of freedom with the largest magnitude PCC is relatively far from the mutating residue. See **Detailed Methods** for more information about this analysis.

### 3 Given sufficient simulation time, AREX and AREST can provide converged and accurate ΔΔG_binding_ predictions

We explore two potential solutions for overcoming the observed sampling challenges: (1) running much longer simulations with the same sampling strategy (AREX) with the goal of exceeding the relevant slow correlation times to enable convergence, and (2) using an enhanced sampling strategy that aims to reduce the correlation times to shorter timescales. For (2), we consider the addition of solute tempering to alchemical replica exchange (AREST). We explore the extent to which each approach improves convergence for the complex phase simulations of all barnase:barstar mutations, with a special focus on A42T and R87A.

#### 3.1 Significantly longer (50 ns/replica) complex phase AREX simulations yield improved ΔG_complex_ convergence and adequate sampling of slow conformational degrees of freedom

We first demonstrate that running longer (50 ns/replica) complex phase AREX simulations improves barnase:barstar ΔΔG_binding_ predictions. We found that the accuracy of the predictions improved: the RMSE decreased from 2.49 (95% CI: [1.32, 3.74]) kcal/mol with 10 ns/replica AREX to 1.61 (95% CI: [1.12, 2.11]) kcal/mol with 50 ns/replica AREX (Figure 6D). Moreover, with 50 ns/replica AREX simulations, 86% (24/28) of predictions are close to experiment and all mutations have the correct sign (Figure 6D, Supplementary Figure 6C). We also found that the internal consistency improved: the RMSE decreased from 3.07 (95% CI: [0.89, 4.76]) kcal/mol for 10 ns/replica AREX to 0.89 (95% CI: [0.25, 1.43]) kcal/mol for 50 ns/replica AREX (Figure 6C). Finally, we found that with 50 ns/replica AREX simulations, the convergence of ΔG_complex_ improved significantly, such that 100% (28/28) of mutations have converged (Supplementary Figure 11B). We then confirmed that the improved convergence for A42T and R87A is a result of more thorough sampling of the likely slowest degrees of freedom associated with each mutation. Examination of representative time series shows that between 10–50 ns, the slow degrees of freedom are sampled more comprehensively than with only 10 ns (Figure 4B, F).

We next show that the poor accuracy with respect to experiment for mutations with significantly discrepant ΔΔG_binding_s (even after 50 ns) are likely due to errors in force field parameters or extreme sampling problems. Despite the improved predictions obtained from running longer AREX, 14% (4/28) of the mutations still demonstrate significantly poor accuracy: D35A, A35D, Q83R, and A29Y (Figure 6D). Common reasons for discrepant ΔΔG_binding_ predictions include insufficient protein or water sampling, errors in force field parameters, and failure to model multiple protonation states [33]. We found that A29Y has a significantly discrepant ΔΔG_binding_ (and relatively poor internal consistency) because the mutant tyrosine residue does not sample the relevant energetically favorable orientations that enable it to contribute favorably to the barnase:barstar interface (details in Supplementary Information B).

We next investigated the causes of discrepancy for the remaining significantly discrepant mutations (D35A, A35D, and Q83R), all of which pass the internal consistency check with sufficient simulation time (100 ns/replica for Q83R and 50 ns/replica for the other two mutations, see Figure 6C and Supplementary Table 1). We first assessed whether the discrepancies are a result of failing to account for all relevant protonation states and found that protonation states are not the cause of these discrepancies (see Supplementary Information C). Given the absence of protonation state problems, the discrepancies are likely due to inaccurate force field parameters or insufficient sampling (of a slow degree of freedom with a correlation time longer than 50 ns). However, it is worth noting that with sufficient simulation time, the sign is correct for each of these discrepant ΔΔG_binding_s, indicating that the estimates are still useful in characterizing whether a mutation is energetically favorable or unfavorable (Figure 6D).

Despite the exceptions described above, we emphasize that running longer AREX improved our barnase:barstar predictions (RMSE is 1.61, 95% CI: [1.12, 2.11] kcal/mol, Figure 6D), indicating that for several mutations, sampling was insufficient with 10 ns/replica AREX but sufficient with 50 ns/replica. Moreover, 50 ns/replica may not be necessary depending on the desired accuracy, e.g., to achieve an RMSE of less than 2 kcal/mol for barnase:barstar predictions, ∼20 ns/replica AREX simulations should be sufficient (Figure 7C).

**Figure 6.**
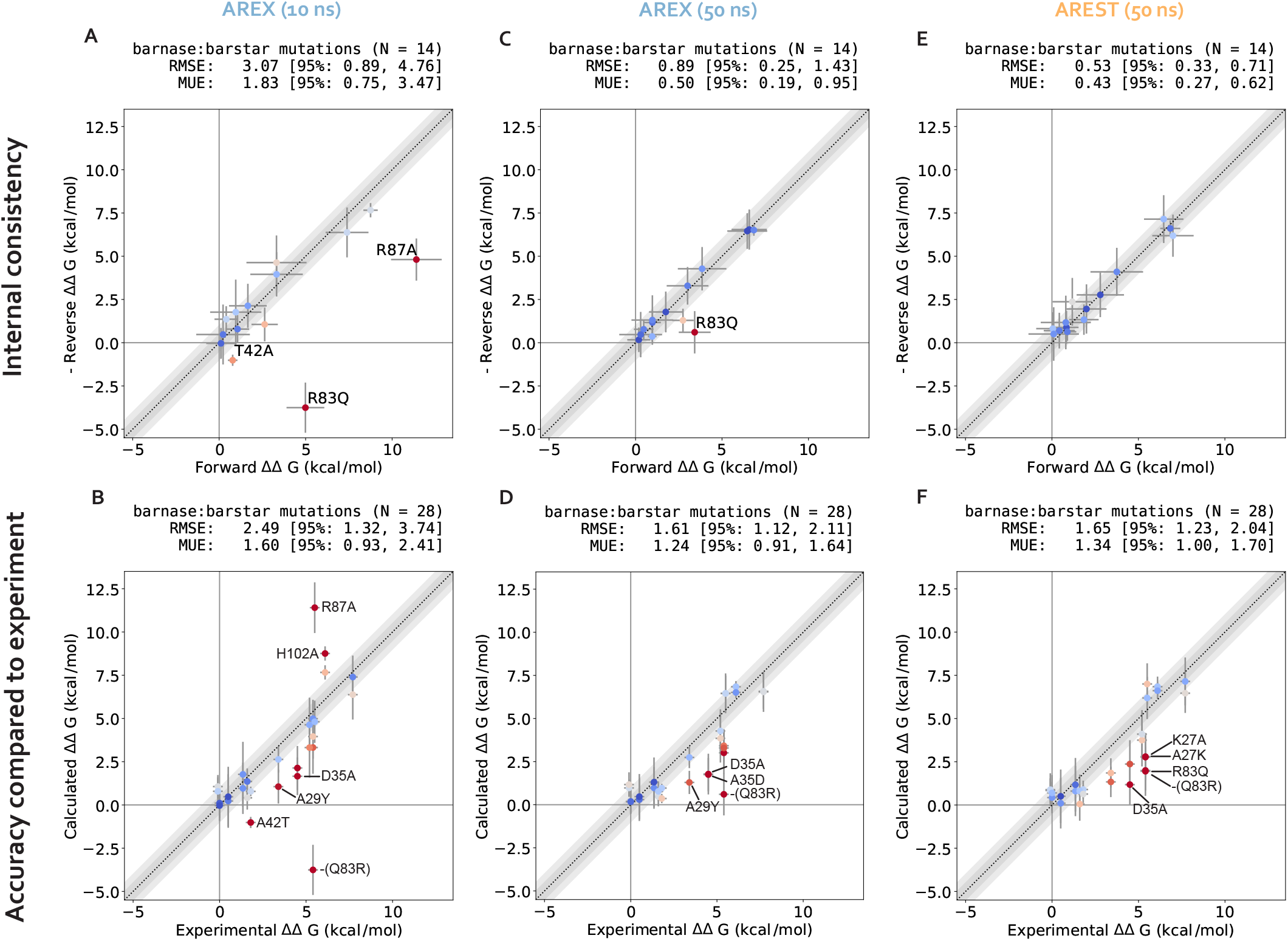
Running long (50 ns/replica) simulations of alchemical replica exchange (AREX) and alchemical replica exchange with solute tempering (AREST) yields improved ΔΔG_binding_ predictions with respect to 10 ns/replica AREX simulations. **(A)** (Negative of the) Reverse versus forward ΔΔG_binding_s for each barnase:barstar mutation computed from AREX simulations (number of states = 24 and 36 for neutral and charge mutations, respectively and simulation time = 10 ns/replica for each phase). The y = x (black dotted) line represents zero discrepancy between forward and (negative of the) reverse ΔΔG_binding_s, the dark gray shaded region represents 0.5 kcal/mol discrepancy, and the light gray region represents 1 kcal/mol discrepancy. Data points are colored by how far they are from zero discrepancy (dark blue and red indicate close to and far from zero, respectively). Data points are labeled if the forward and (negative of the) reverse ΔΔG_binding_s are not within statistical error of each other (i.e., neither the forward nor the negative reverse ΔΔG_binding_ is within 1 kcal/mol of the 95% CI for the other ΔΔG_binding_). Error bars represent two standard deviations and were computed by bootstrapping the decorrelated reduced potential matrices 200 times. Root mean square error (RMSE) and mean unsigned error (MUE) are shown with 95% confidence intervals obtained from bootstrapping the data 1000 times. **(B)** Calculated versus experimental ΔΔG_binding_s for each barnase:barstar mutation computed from AREX simulations (number of states = 24 and 36 for neutral and charge mutations, respectively and simulation time = 10 ns/replica for each phase). The y = x (black dotted) line represents zero discrepancy between calculated and experimental ΔΔG_binding_s, the dark gray shaded region represents 0.5 kcal/mol discrepancy, and the light gray region represents 1 kcal/mol discrepancy. Data points are labeled if the 95% CIs of the calculated and experimental ΔΔG_binding_s are not within 1 kcal/mol of each other. For more details on the plot and error bars, refer to the caption for panel A. **(C)** Same as (A), but using 50 ns/replica AREX simulations for the complex phase and 10 ns/replica AREX simulations for the apo phase instead of 10 ns/replica AREX simulations for both phases. **(D)** Same as (B), but using 50 ns/replica AREX simulations for the complex phase and 10 ns/replica AREX simulations for the apo phase instead of 10 ns/replica AREX simulations for both phases. **(E)** Same as (A), but using 50 ns/replica AREST simulations for the complex phase and 10 ns/replica AREX simulations for the apo phase instead of 10 ns/replica AREX simulations for both phases. **(F)** Same as (B), but using 50 ns/replica AREST simulations for the complex phase and 10 ns/replica AREX simulations for the apo phase instead of 10 ns/replica AREX simulations for both phases.

#### 3.2 AREST convergence is comparable to that of AREX for most mutations

We next demonstrate that running 50 ns/replica AREST simulations also yields improved barnase:barstar ΔΔG_binding_ with respect to 10 ns/replica AREX simulations (for a comparison to 10 ns/replica AREST simulations, see Supplementary Information A.7). We ran 50 ns/replica AREST (with radius = 0.5 nm and *T*_max_ = 600 K, see Supplementary Information D for details on REST parameter selection) for the complex phase of all barnase:barstar mutations and observed sufficient replica mixing for all mutations (Supplementary Figure 12). We observed improvement in the accuracy with respect to experiment; the RMSE decreased from 2.49 (95% CI: [1.32, 3.74]) kcal/mol for 10 ns/replica AREX to 1.65 (95% CI: [1.23, 2.04]) kcal/mol for 50 ns/replica AREST (Figure 6F, Supplementary Figure 6D). We also found that the internal consistency significantly improved with the RMSE decreasing from 3.07 (95% CI: [0.89, 4.76]) kcal/mol for 10 ns/replica AREX to 0.53 (95% CI: [0.33, 0.71]) kcal/mol for 50 ns/replica AREST (Figure 6E). Finally, we also observed that with 50 ns/replica AREST simulations, 100% (28/28) of the complex free energy differences (ΔG_complex_s) converge (Supplementary Figure 11C).

We next show that while the two methods predict similar ΔΔG_binding_s for each mutation (Supplementary Figure 14), AREST converges more efficiently than AREX for two mutations with sampling problems, but not for the rest of the barnase:barstar mutations. We monitored the discrepancy in predicted ΔΔG_binding_ with respect to experiment as a function of time and compared the discrepancy time series for 50 ns/replica AREST versus that from 50 ns/replica AREX simulations. We analyzed the discrepancies in ΔΔG_binding_s for each of the seven mutations identified as potentially containing sampling problems due to slow ΔG_complex_ convergence with 10 ns/replica AREX: A42T, R87A, R83Q, Q83R, H102A, A35D, and A39D (Figure 3G). For A42T, the AREST discrepancy flattens out (to a close-to-zero discrepancy) more quickly than that of AREX, indicating that for A42T, AREST converges with less simulation time than AREX (Figure 7A). Similarly, for R87A, the AREST discrepancy starts to flatten out around 10 ns, while the AREX discrepancy doesn’t start to flatten out until ∼40 ns, demonstrating that for R87A, AREST converges faster than AREX (Figure 7D). We next investigated why AREST yields faster convergence by comparing AREX and AREST sampling of the likely slowest degrees of freedom (T42 *χ*_1_ angle for A42T and number of waters near A87 for R87A) in representative time series. We found that AREST more thoroughly samples these degrees of freedom and the statistical inefficiencies of the AREST time series are smaller than those of AREX, indicating that the faster convergence of AREST is due to reduction of relevant correlation times (Figure 7B, E).

Importantly, we found that for the remaining 71% (5/7) of mutations potentially containing sampling problems, the discrepancy in ΔΔG_binding_ does not converge to zero significantly faster for AREST than AREX (Supplementary Figure 15). Finally, to assess convergence across all mutations, we monitored the root mean square error (RMSE) and mean unsigned error (MUE) over time and observed that for both RMSE and MUE, the AREX and AREST time series are within error of each other (Figure 7C, F). Therefore, although AREST shows faster convergence than AREX for A42T and R87A, AREST convergence is comparable to that of AREX when comparing the two sampling strategies over all barnase:barstar mutations.

## DISCUSSION

### Widespread application of RBFE calculations to protein:protein complexes is primarily limited by the simulation time required to achieve reliable estimates

For some mutations, running RBFE calculations long enough to achieve converged, accurate, and reliable predictions can be computationally expensive, depending on the simulation time required and the computing resources available. For example, to achieve highly accurate RBFE predictions (RMSE ∼1.6 kcal/mol) for barnase:barstar, the most challenging mutations (i.e., charge-changing mutations with sampling challenges) require 50 ns/replica for the complex phase and 10 ns/replica for the apo phase. This amounts to ∼220 graphics processing unit (GPU) hours per mutation on an NVIDIA A100 graphics card—at the cost of roughly $920 per mutation on an equivalent instance on Amazon Web Services (AWS) (Supplementary Information A.12). However, we emphasize that we obtained converged and accurate ΔΔG_binding_ estimates for most of the mutations with 10 ns/replica AREX (Section 1.2), indicating that most mutations would not require such computationally expensive simulations (and instead would cost ∼62 GPU hours and $260 per mutation on AWS). Taken together, our results demonstrate that given current best practices sampling strategies and state-of-the-art computing resources, the primary limiting factor in applying RBFE calculations to protein:protein complexes is the computational cost associated with achieving sufficient sampling for a small subset of mutations.

**Figure 7.**
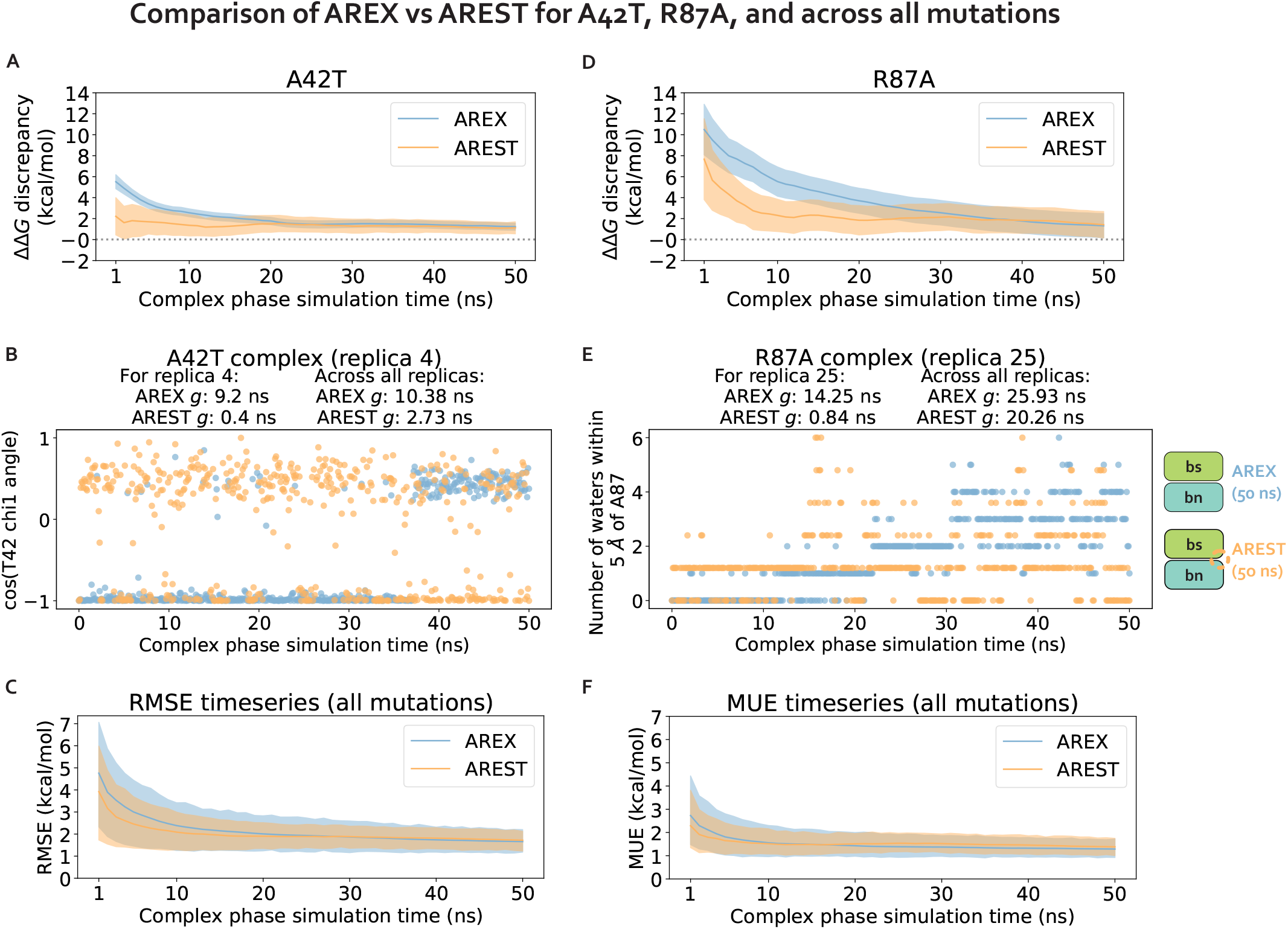
Alchemical replica exchange with solute tempering (AREST) and alchemical replica exchange (AREX) demonstrate comparable convergence for most barnase:barstar mutations. **(A)** ΔΔG_binding_ discrepancy (with respect to experiment) time series for A42T. The discrepancy was computed as ΔG_complex_ − ΔG_apo_ − ΔΔG_experiment_, where ΔG_complex_ corresponds to the (AREX or AREST) complex phase ΔG at a particular time point, ΔG_apo_ corresponds to the apo phase ΔG computed from a 10 ns/replica AREX simulation, and ΔΔG_experiment_ is the experimental value from Schreiber et al [73]. AREX time series shown in blue and AREST time series (with radius = 0.5 nm, *T*_max_ = 600 K) shown in orange. Number of states is 24 for both AREX and AREST. Shaded regions represent ± two standard deviations, computed by bootstrapping the decorrelated reduced potential matrices 200 times. Gray dashed line indicates ΔΔG_binding_ discrepancy = 0. **(B)** Time series of the *χ*_1_ angle for residue T42 for a representative replica (replica 4) of the A42T complex phase AREX simulation (blue) and AREST simulation (orange) (number of states = 24, simulation time = 50 ns/replica). *g* indicates statistical inefficiency, which is proportional to the correlation time. *g* = 0.1 ns indicates very thorough sampling (because the sampling interval is 0.1 ns) and large values of *g* indicate poor sampling. **(C)** Time series of the root mean square error (RMSE) (with respect to experiment) for the ΔΔG_binding_s of all barnase:barstar mutations. The ΔΔG_binding_s used to compute the RMSE at each time point were computed as ΔG_complex_ − ΔG_apo_ for each mutation, where ΔG_complex_ corresponds to the (AREX or AREST) complex phase ΔG at a particular time point and ΔG_apo_ corresponds to the apo phase ΔG computed from a 10 ns/replica AREX simulation. AREX time series shown in blue and AREST time series (with radius = 0.5 nm, *T*_max_ = 600 K) shown in orange. Number of states is 24 for neutral mutations and 36 for charge-changing mutations. Shaded regions represent ± two standard deviations, computed by bootstrapping 1000 times. **(D)** Same as (A), but for R87A instead of A42T. Number of states is 36 for both AREX and AREST. **(E)** Time series of the number of waters within 5 Å of residue A87 for representative replica (replica 25) of the R87A complex phase AREX simulation (blue) and AREST simulation (orange) (number of states = 36, simulation time = 50 ns/replica). **(F)** Same as (C) but for mean unsigned error (MUE) instead of RMSE.

Given that similar types of sampling problems are also challenging for small molecule transformations [54, 55, 83], finding ways to reduce computational cost for alchemical transformations with difficult sampling problems will be highly useful for the development of alchemical free energy calculations in general. One straightforward approach for reducing computational cost involves waiting for improvements in hardware. GPU performance has rapidly improved over the last decade and will continue to improve in the coming years [84]. There are also particularly exciting developments in the realm of cheaper parallelization through the introduction of wider GPUs that enable a single GPU to be partitioned into multiple instances (e.g., NVIDIA’s Multi-Instance GPU feature).

### Improvement of AREX and AREST simulation parameters may reduce the simulation time required for converged ΔΔG estimates for mutations with sampling challenges

Beyond anticipating advancements in hardware, a promising avenue for decreasing computational cost involves further optimizing the AREX and AREST simulation parameters used in this study. For both AREX and AREST, we chose the same number of alchemical intermediate states for all neutral mutations and a different, larger number of states for all charge-changing mutations. Additionally, we defined the alchemical and REST scaling protocols for each state according to simple, piecewise linear functions. Moreover, for AREST, we chose the REST parameters (radius and *T*_max_) by exploring a small set of extreme REST parameters (Supplementary Information D).

Although we confirmed that our AREX and AREST parameter choices do not result in any replica mixing bottlenecks (Supplementary Figures 2, 5, 12), there are likely alternative protocol parameters which could provide more efficient ΔΔG_binding_ convergence. Ideally, each mutation would have optimized protocol parameters that provides a converged and accurate ΔΔG_binding_ estimate in the minimal amount of simulation time. However, because the search space for each of the parameters is large, brute-force optimization is unfeasible and even exploration of extreme values for each parameter for each mutation would be quite computationally expensive. Therefore, future work could involve development of methods for mutation-specific parameter optimization. Furthermore, there are also opportunities for optimizing mutation-independent protocol parameters, such as the integrator timestep [85], alchemical functional form [86–89], and Particle Mesh Ewald error tolerance [67] which may reduce simulation time.

### Adaptation of other enhanced sampling methods for use in alchemical free energy calculations may also decrease the simulation time required to sufficiently sample difficult transformations

There are many existing methods for enhancing sampling in molecular dynamics simulations [90], many of which accelerate sampling of known slow degrees of freedom in a targeted manner [50–53, 91–96]. Some existing enhanced sampling methods also identify the slow degrees of freedom (as an intermediate step) [97, 98], but they do not necessarily identify the slow degrees of freedom that are most highly coupled to the alchemical coordinate (i.e. *∂U* /*∂λ*), which are responsible for slow convergence of RBFEs. Future work could involve incorporating existing enhanced sampling methods into alchemical free energy calculations to further improve sampling and convergence, as has been demonstrated for simple test systems [99]. Furthermore, when adapting methods that identify slow degrees of freedom, it will be important to account for coupling to the alchemical coordinate.

## CONCLUSIONS

In this work, we explored the sampling challenges associated with applying relative binding free energy (RBFE) calculations to estimate the impact of protein mutations in a model protein:protein complex (barnase:barstar). We found that sampling problems are absent when the mutation is not located in the context of a complex network of protein interactions (i.e. in terminally-blocked amino acids), but are present in the complex phase for several barnase:barstar mutations, yielding slow convergence of ΔG_complex_s. Moreover, most of the mutations with complex phase sampling and convergence problems involve charge-changes. Furthermore, we attributed the barnase:barstar complex phase sampling problems to specific slow degrees of freedom (individual sidechain torsions, interfacial contacts, and nearby waters) which are highly dependent on the mutation. Finally, we found that given sufficient simulation time (50 ns/replica), both AREX and AREST can address most of the aforementioned sampling problems, with both methods demonstrating comparable convergence for most mutations.

Ultimately, our analyses and findings provide a model framework for diagnosing and mitigating sampling problems in other protein:protein complexes. By facilitating deep investigation of these sampling challenges in an open-source manner, our study lays the groundwork for the development of better methods for improving sampling in protein:protein RBFE calculations and free energy calculations in general.

## 4 Acknowledgments

We would like to acknowledge Gyorgy Snell (ORCID: 0000-0003-1475-659X) for helpful input during project initiation. We also thank Peter Eastman (0000-0002-9566-9684) and Levi Naden (0000-0002-3692-5027) for helpful feedback on software implementation (Perses [56], https://github.com/choderalab/perses and OpenMMTools, https://github.com/choderalab/openmmtools). We thank OpenEye Scientific Software and Schrodinger for providing us with free academic licenses. We are grateful to Hannah Bruce MacDonald (0000-0002-5562-6866), Hannah Baumann (0000-0002-1736-7744), Bert de Groot (0000-0003-3570-3534), the reviewers, and the editors for their constructive feedback.

## 7 Disclaimers

The content is solely the responsibility of the authors and does not necessarily represent the official views of the National Institutes of Health.

## 8 Author Contributions

Conceptualization: IZ, JDC; Data Curation: IZ; Formal Analysis: IZ; Funding Acquisition: JDC, LER; Investigation: IZ; Methodology: IZ, DAR, JDC, KH; Project Administration: JDC, LER, SS; Resources: JDC, LER; Software: IZ, DAR, IP, MMH; Supervision: JDC, LER, SS; Visualization: IZ; Writing – Original Draft: IZ; Writing – Review & Editing: IZ, JDC, SS, DAR, LER, KH, IP, MMH

## Supplementary Information

Supplementary Figures 1-16 and Supplementary Tables 1-2, along with more details on the methods and follow up investigation on outlier mutations.

## A Detailed Methods

### A.1 Data and code availability

The data and Python code used to produce the results discussed in this paper is distributed open-source under a MIT license and is available at https://github.com/choderalab/perses-barnase-barstar-paper.

Core dependencies include Perses 0.10.1 [56], OpenMMTools 0.21.5 (https://github.com/choderalab/openmmtools), MDTraj 1.9.7 [100], and pymbar 3.1.1 [66]. OpenMM 8.0.0beta (https://anaconda.org/conda-forge/openmm/files?version=8.0.0beta — build 0), a development version of OpenMM 7 [29], was used to generate the input files for alchemical replica exchange (AREX) and alchemical replica exchange with solute tempering (AREST), run equilibration, and run AREX for the terminally-blocked amino acids. OpenMM 7.7.0.dev2 (https://anaconda.org/conda-forge/openmm/files?version=7.7.0dev2), a development version of OpenMM 7 [29] which was built after OpenMM 8.0.0beta and contains a performance enhancement for AREX and AREST, was used for running all other AREX and AREST simulations.

ΔΔG comparison plots were generated with cinnabar 0.3.0 (https://github.com/OpenFreeEnergy/cinnabar). All other plots were generated using Matplotlib 3.5.2 [101] and structural images were generated using PyMOL 2.5.1 [102].

### A.2 Structure preparation

#### Capped peptides

To create structures for the terminally-blocked amino acids, tleap from AmberTools 21.9 [103] was used to generate the ACE-, NME-capped (ACE-X-NME) and zwitterionic ALA-capped (ALA-X-ALA) peptides in idealized alpha helical conformations (see https://github.com/choderalab/perses-barnase-barstar-paper/blob/main/input_files/generate_peptide_pdbs.py).

#### Barnase:barstar

To create a structural model of the wild-type (WT) barnase:barstar complex, chains A and D (which correspond to barnase and barstar, respectively) were extracted from the crystal structure with PDB ID 1BRS [75] because they are the chains with the highest overall quality (see wwPDB X-ray Structure Validation Report). Schrodinger Maestro 2021-2 [104] was used to prepare the structure with the Protein Prep Wizard, i.e., delete the other chains, fill in missing side chains and loops, cap the termini, add hydrogens, and optimize the hydrogen bond network (using pH 8.0, the pH used in Schreiber et al. binding experiments [73]). HIS18 (in barnase) was protonated as HID, HIS102 (in barnase) was protonated as HIE, and H17 (in barstar) was protonated as HID. Default settings were used unless otherwise noted. Because Maestro added NMA caps as inserted residues (i.e., the residue ID was the same as the preceding residue with the addition of an “A”), OpenMM 8.0.0beta [29] was used to rename the NMA residue to NME as well as renumber the NME residue and all subsequent residues to have residue IDs incremented by 1 (see https://github.com/choderalab/perses-barnase-barstar-paper/blob/main/input_files/renumber.py).

Although the experimental relative binding free energy (ΔΔG_binding_) data (Schreiber et al. [73]) was generated using the WT sequences of barnase and barstar, which contain cysteines at barstar residues 40 and 82, the 1BRS structure contains alanines at those positions. Residues 40 and 82 were not mutated to cysteines in our structural model because it has been demonstrated that the structures, activities, and stabilities of mutant (A40 and A82) barstar are similar to those of WT (C40 and C82) barstar [105].

The prepared WT structure was used as the starting structure for forward mutations. For the reverse mutations, the starting structures were mutant barnase and barstar structures, which were generated by mutating the residue of interest in the prepared WT structure using Maestro 2021-3 [104]. The sidechain rotamer that best matched the sidechain orientation of the WT residue was selected.

For the D35A and K27A experiments (accounting for multiple protonation states), models of barstar with ASH35, barnase with LYN27, and terminally blocked amino acids with ASH or LYN were generated by modifying the protonation state of the prepped WT structures using OpenMM 8.0.0beta [29] (see https://github.com/choderalab/perses-barnase-barstar-paper/blob/main/input_files/generate_nonstandard_protonation_states.py).

### A.3 System solvation and parameterization

Solvation and parameterization were performed with OpenMM 8.0.0beta [29]. The systems were solvated using the TIP3P rigid water model [106] in a cubic box with 12 Å and 17 Å solvent padding on all sides for barnase:barstar and terminally-blocked amino acids, respectively. The solvated systems were then minimally neutralized with 50 mM NaCl using the Li/Merz ion parameters of monovalent ions for the TIP3P water model (12-6 normal usage set) [107]. The systems were parameterized with the Amber ff14SB force field [108]. Amber ff14SB allows naked charges on certain hydrogens, i.e., atoms with a non-zero charge, but zero *σ* or *ϵ*. To prevent naked charges from causing simulation failures due to nuclear fusion when enhanced sampling strategies are employed, a small padding was added to each non-water atom with *σ* = 0 nm or *ϵ* = 0 nm. If the atom had *σ* = 0 nm, 0.06 nm padding was added. If the atom had *ϵ* = 0 kJ/mol, 0.0001 kJ/mol padding was added. Finally, if *ϵ* = 0 kJ/mol and *σ* = 1, sigma was set to 0.1 nm. Full details and scripts can be found at: https://github.com/choderalab/perses-barnase-barstar-paper/blob/main/scripts/01_generate_solvated_inputs/generate_solvated_inputs.py.

### A.4 System equilibration

To ensure that our experiments regarding convergence are not the result of structure preparation errors or crystallographic artifacts abruptly followed by production simulation, the barnase:barstar systems were gently equilibrated over 9 stages based on a previously described protocol [109] using OpenMM 8.0.0beta [29]. The number of steps, temperature, ensemble, collision rate, timestep, and force constant for each stage are detailed in the aforementioned reference. The reference protocol was run with a few adjustments:

1. The heavy atoms were restrained in the first four stages, backbone atoms were restrained in the next four stages, and no atoms were restrained for the last stage,
2. A Langevin integrator was used (see below for more details), so the Berendsen thermostat in the reference protocol was not necessary,
3. The last stage of gentle equilibration was extended to 9.25 ns (instead of 5 ns), so that the whole equilibration protocol would involve 10 ns of simulation.

The energy minimization stages were performed using the OpenMM 8.0.0beta LocalEnergyMinimizer with an energy tolerance of 10 kJ/mol. The molecular dynamics stages used the OpenMM 8.0.0beta LangevinMiddleIntegrator [85, 110, 111]. Hydrogen atom masses were set to 3 amu by transferring mass from connected heavy atoms, bonds to hydrogen were constrained, and center of mass motion was not removed. Pressure was controlled by a molecular-scaling Monte Carlo barostat with a pressure of 1 atmosphere, a temperature of 300 K, and an update interval of 50 steps. Non-bonded interactions were treated with the Particle Mesh Ewald method [112] using a real-space cutoff of 1.0 nm and an Ewald error tolerance of 0.00025, with grid spacing selected automatically. Long range anisotropic dispersion corrections were applied to un-scaled (non-REST and non-alchemical) steric interactions [113]. Because their structural models did not originate from crystal structures, the terminally-blocked amino acid systems were not equilibrated with the gentle equilibration protocol; they were minimized and then equilibrated for 10 ns without restraints in the NPT ensemble at 300 K with a collision rate of 2 picoseconds^-1^ and a timestep of 2 femtoseconds. For the barnase:barstar complex systems, a virtual bond was added between the first atoms of each protein chain to ensure that the chains are imaged together. Default parameters were used unless noted otherwise. Further details on the equilibration protocol are available at: https://github.com/choderalab/perses-barnase-barstar-paper/blob/main/scripts/02_run_equilibration/run_equilibration.py.

### A.5 Free energy calculation input file preparation

The hybrid topology, positions, and system for each transformation were generated using Perses 0.10.1 [56] and OpenMM 8.0.0beta [29]. The hybrid topology was generated using a single topology approach. The hybrid positions were assembled by copying the positions of all atoms in the WT (“old”) topology and then copying the positions of the atoms unique to the mutant (“new”) residue (i.e., unique new atoms). The unique new atom positions were generated using the Perses `FFAllAngleGeometryEngine`, which probabilistically proposes positions for one atom at a time based on valence energies alone. Further details on hybrid topology, positions, and system generation (including definitions of the valence, electrostatic, and steric energy functions) are available in the Perses `RESTCapableHybridTopologyFactory` class.

For charge-changing mutations, counterions were added to neutralize the mutant system by selecting water molecules in the WT system that are initially at least 8 Å from the solute and alchemically transforming the WT water molecules into sodium or chloride ions in the mutant system. For example, if the mutation was ALA→ASP, a water molecule in the WT ALA system was transformed into a sodium ion in the mutant ASP system to keep the system at the ASP endstate neutral. If the mutation was GLU→ALA, a water molecule in the WT GLU system was transformed into a chloride ion in the mutant ALA system. Additional details on the counterion implementation can be found in the Perses `_handle_charge_changes()` function found in `perses.app.relative_point_mutation_setup`.

To prevent singularities from arising when turning off the nonbonded interactions involving unique old or unique new atoms, a softcore approach was used that involves “lifting” unique old or unique new interaction distances into the “4th dimension.” A padding distance (*w*(*λ*), see equation 2) was added to the interaction distances involving unique old or unique new atoms so that the atoms could not be on top of each other [68]. *w*_lifting_ (the maximum value for *w*(*λ*)) was selected to be 0.3 nm and when AREX was performed for all terminally-blocked amino acid mutations, replica mixing was sufficient for all mutations, indicating that the thermodynamic length between alchemical states was reasonable even given the softcore lifting term (Supplementary Figure 2). This 4D lifting softcore approach was applied to both the electrostatic and steric interactions, so multi-stage alchemical protocols (e.g., where electrostatics must be turned off before sterics) were not necessary for scaling on or off the electrostatic and steric interactions. Instead, a simple linear protocol was used for interpolating the valence, nonbonded, and lifting terms (Supplementary Figure 1A). This softcore approach is very similar to traditional softcore approaches [86, 114] with the main difference being that for the Lennard Jones potential, our approach uses a lifting distance (*w*(*λ*)) that is independent of *σ* (the distance at which the Lennard Jones potential energy equals zero), whereas the aforementioned traditional approaches define the lifting distance as a function of *σ*. In our approach, the lifting distance was defined to be independent of sigma for simplicity and ease of implementation.

### A.6 Alchemical replica exchange

Alchemical replica exchange (AREX) simulations were performed using Perses 0.10.1 [56] and OpenMMTools 0.21.5 (https://github.com/choderalab/openmmtools). OpenMM 8.0.0beta [29] was used for the terminally-blocked amino acids and OpenMM 7.7.0.dev2 [29] was used otherwise. The alchemical protocol was defined with evenly spaced *λ* values between 0 and 1 (Supplementary Figure 1A). Before AREX was performed, the positions were minimized at each of the alchemical states using the OpenMM `LocalEnergyMinimizer` with an energy tolerance of 10 kJ/mol and a maximum of 100 iterations (except for D39A, A76E, and A39D complex phase AREST simulations, which were minimized without a limit on the number of iterations because instabilities were present with only 100 iterations). Each AREX cycle consisted of running 250 steps (4 femtosecond timestep) with the OpenMM 8.0.0beta `LangevinMiddleIntegrator` [85, 110, 111] at a temperature of 300 K, a collision rate of 1 picosecond^-1^, and a constraint tolerance of 1e-6. All-to-all replica swaps were attempted every cycle [72]. Replica mixing plots were created using OpenMMTools 0.21.5 (https://github.com/choderalab/openmmtools) to extract the mixing statistics from the AREX trajectories. Default settings were used unless otherwise noted. For full details on the AREX implementation: https://github.com/choderalab/perses-barnase-barstar-paper/blob/main/scripts/04_run_repex/run_repex.py.

For terminally-blocked amino acid mutations, the two simulation phases involved different types of caps — the first phase was ACE-X-NME and the other phase was ALA-X-ALA, where X is an amino acid. AREX simulations were run for each phase using 12 replicas for neutral mutations and 24 replicas for charge-changing mutations. While ASH2A and LYN2A are both neutral mutations, 24 replicas were used for each to allow for direct comparison with D2A and K2A. Replicas mixed well for all mutations, indicating good phase space overlap (Supplementary Figure 2). 5000 cycles (i.e., 5 ns) were run for each replica, resulting in 60 ns of sampling per phase per neutral mutation and 120 ns of sampling per phase per charge-changing mutation. Since the AREX simulation time for terminally-blocked amino acid mutations was shorter than that of barnase:barstar mutations (5 ns/replica vs. 10 ns/replica), the ΔΔ*G*s for the terminally-blocked amino acid mutations were computed using fewer samples, which explains the larger error bars for terminally-blocked amino acids (Figure 3A) as compared to barnase:barstar mutations (Figure 3B).

For barnase:barstar mutations, apo and complex phase AREX simulations were performed with 24 replicas for neutral mutations and 36 replicas for charge-changing mutations (including H102A, even though histidine was modeled as HIE). While ASH35A and LYN27A are both neutral mutations, we used 36 replicas for each mutation to allow for direct comparison with D35A and K27A. Replicas mixed well for all mutations, indicating good phase space overlap (Supplementary Figure 5). 10000 cycles were initially run per replica (10 ns/replica), resulting in 240 ns of sampling per phase per neutral mutation and 360 ns of sampling per phase per charge-changing mutation. The complex phase simulations were extended to 50 ns/replica, resulting in 1200 ns per phase per neutral mutation and 1800 ns per phase per charge-changing mutation.

To improve the accuracy of our predicted free energy differences, the sampled alchemical states were bookended with “virtual endstates,” which were not sampled during free energy calculation, but for which reliable estimates of the physical endstates could be robustly produced during analysis. In these bookended endstates, nonbonded interaction energies were defined using the more accurate, but more computationally expensive, Lennard Jones with Particle Mesh Ewald (LJPME) method [115] to better account for the heterogeneous long-range dispersion interactions known to be important when creating or destroying many atoms in alchemical free energy calculations [113]. For full details on the unsampled endstate implementation, see: `perses.dispersed.utils.create_endstates_from_real_systems()`.

To run AREX simulations with heavy-atom coordinate restraints, an OpenMM 7.7.0.dev2 `CustomCVForce` was added to the hybrid system with the energy expression:

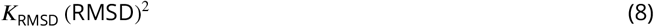

where ***K***_RMSD_ (the harmonic force constant) was chosen to be 50 kcal/molÅ^2^ for A42T and 75 kcal/molÅ^2^ for R87A in order to sufficiently reduce heavy-atom motion without causing instabilities. RMSD was computed using an OpenMM 7.7.0.dev2 `RMSDForce` [29] (added as a collective variable to the `CustomCVForce`). The two forces (`CustomCVForce` and `RMSDForce`) enable restraint of heavy atoms to their initial positions. For full details on the restraint implementation: https://github.com/choderalab/perses-barnase-barstar-paper/blob/main/scripts/04_run_repex/run_repex.py.

### A.7 Alchemical replica exchange with solute tempering (AREST)

The AREST simulations were performed using Perses 0.10.1 [56], OpenMMTools 0.21.5 (https://github.com/choderalab/openmmtools), and OpenMM 7.7.0.dev2 [29] with the same parameters used in alchemical replica exchange above. Replica mixing plots were created using OpenMMTools 0.21.5 (https://github.com/choderalab/openmmtools) to extract the mixing statistics from the AREX and AREST trajectories. Replica mixing was sufficient for all mutations, indicating decent phase space overlap (Supplementary Figure 12). For the REST-specific parameters, ***T***_max_ and REST radius, all pairwise combinations of small, medium, and large values were explored for A42T and R87A. 400 K, 600 K, and 1200 K were selected for ***T***_max_ and 0.3 nm, 0.5 nm, and 0.7 nm were selected for radius, yielding nine combinations of REST parameters. 0.5 nm and 600 K were selected for ***T***_max_ and radius, respectively, for running complex phase AREST simulations for all mutations. The protocol used to scale the effective temperature is shown in Supplementary Figure 1. Full details and script for the AREST simulations can be found at https://github.com/choderalab/perses-barnase-barstar-paper/blob/main/scripts/04_run_repex/run_repex.py.

To supplement Figure 6, the internal consistency, accuracy, and ΔG_complex_ convergence were analyzed for 10 ns/replica AREST (Supplementary Figures 10A-B, 6B, 11A). There were improvements across all metrics with respect to 10 ns/replica AREX, but not with respect to 50 ns/replica AREST.

The extent to which AREX versus AREST (50 ns/replica complex phase simulations) explores lambda space was also assessed. The AREST replica state index *g* tends to be larger than that of AREX for most mutations, indicating that AREST traverses lambda space (i.e., state space) less efficiently than AREX. AREST’s worse visitation of lambda space is likely because the introduction of REST increases the thermodynamic length between lambda windows (Supplementary Figure 13).

### A.8 Free energy difference analysis

Free energy differences (ΔGs) for each phase were estimated using the MBAR implementation in pymbar 3.1.1 [66]. The MBAR estimates were initialized with zeroes for all experiments except R2Q (ACE-X-NME phase) and two of the REST combination experiments (R87A with radius 0.5 nm and ***T***_max_ 600 K and R87A with radius 0.7 nm and ***T***_max_ 1200 K), which were initialized with a BAR estimate to improve solver convergence. The MBAR equations were solved using an adaptive algorithm with a solver tolerance of 1e-12. The algorithm runs both self-consistent and Newton-Raphson methods at each iteration and the method with the smallest gradient is chosen to improve numerical stability. Error bars were computed by bootstrapping the decorrelated reduced potential matrices (number of bootstraps = 200) and evaluating the free energy differences for each bootstrapped matrix with a solver tolerance of 1e-6. To assemble the decorrelated samples to feed into MBAR, the number of equilibration iterations to discard and the subsample rate were determined by applying a simple equilibration detection method [79] (implemented in OpenMMTools 0.21.5, https://github.com/choderalab/openmmtools) to a timeseries of the sum of the reduced potentials over all replicas. Default settings were used unless otherwise noted. For full details on ΔG estimation, see: https://github.com/choderalab/perses-barnase-barstar-paper/blob/main/scripts/05_analyze/analyze_dg.py.

The ΔG time series were generated by estimating the ΔG (using MBAR, as described above) in 1 ns intervals. The MBAR estimates were initialized with zeroes for all experiments except R2Q (ALA-X-ALA phase), which was initialized with a BAR estimate to improve solver convergence. The first 10% of samples were discarded due to equilibration and samples were selected every 5 iterations. Error bars were computed as described above. The slope (and standard deviation) of the last 5 ns was computed using SciPy 1.9.0’s `linregress` function [116]. For the restraint experiments, residual ΔG time series plots were generated in the same manner as described above. The residual ΔG was computed as ΔG(*t*) − ΔG(*t* = 10ns), which was necessary to compare the rate of decay of the ΔGs from the non-restrained and restrained simulations, otherwise the two time series differ by an offset. For the REST parameter comparison experiments, the “true” ΔG was computed by averaging the ΔG over three replicates of 100 ns/replica AREX simulations. For comparison of AREX versus AREST, the ΔΔG discrepancy, RMSE, and MUE time series plots were computed in the same manner as described above, where the ΔΔG discrepancy for each time point was computed as ΔG_complex_ ΔG_apo_ ΔΔG_experiment_ and RMSE and MUE for each time point were computed using the ΔΔG_predicted_s for all 28 mutations.

ΔΔG comparison plots (forward vs negative reverse and calculated vs experiment) were generated using Cinnabar 0.3.0 (https://github.com/OpenFreeEnergy/cinnabar). For more details on generating these plots, see: https://github.com/choderalab/perses-barnase-barstar-paper/blob/main/scripts/05_analyze/0_cinnabar_plots_50ns.ipynb.

### A.9 *ϕ* and *ψ* angle analysis

*ϕ* and *ψ* angle analysis was performed for 5 ns/replica A2T and R2A ACE-X-NME phase AREX simulations (Supplementary Figure 4). The *ϕ* and *ψ* angles were computed for the old positions of each replica trajectory snapshot (saved every 100 ps) for all replicas using MDTraj 1.9.7 [100]. The sine transformation was applied to the angle values in each time series. The statistical inefficiency across all replicas was computed using pymbar 3.1.1 [80]’s `statisticalInefficiencyMultiple`.

### A.10 Y29 residue pair distance analysis

For the Y29 residue pair distance analysis (Supplementary Figure 8), Y29-H102 distances were computed between the closest sidechain heavy atoms and Y29-R83 and Y29-N84 distances were computed between the carbonyl oxygen of R83 or N84 and the sidechain hydroxyl oxygen of Y29. The three residue pair distances were computed for each snapshot (saved every 100 ps) of two different trajectories: 1) the old positions of Y29A AREX (50 ns/replica) at the lambda = 0 endstate and 2) the new positions of A29Y AREX (50 ns/replica) at the lambda = 1 endstate. Distances were computed using MDTraj 1.9.7 [100].

### A.11 *∂U* /*∂λ* correlation analysis

For the *∂U* /*∂λ* correlation analysis (Figure 4B-C, E-F, Figure 5), we monitor the derivative of the potential energy with respect to the alchemical coordinate *λ, ∂U* /*∂λ*, over time. *∂U* /*∂λ* is sensitive to potential energy changes in the alchemical region but insensitive to changes in non-alchemical interactions. An ideal *∂U* /*∂λ* trajectory thoroughly samples a stationary distribution (i.e., it samples all thermally accessible metastable states multiple times), generating a sufficient number of decorrelated samples, which are required in order to produce reliable estimates of free energy differences. On the other hand, if a *∂U* /*∂λ* trajectory gets stuck in one metastable state and fails to visit all metastable states multiple times, there are likely slow degrees of freedom with long correlation times that make it difficult to obtain decorrelated samples. The degree of correlation within a time series can be quantified by computing its statistical inefficiency, g.

To generate the time series for *∂U* /*∂λ* and each degree of freedom, interface residues were defined as all residues within 4 Å of the other chain, with the addition of barstar residue E80 because it is one of the mutating residues in the Schreiber et al ΔΔG_binding_ dataset. Protein and water degrees of freedom were computed for both the old and new positions of each trajectory snapshot (saved every 100 ps) for all replica trajectories in an automated fashion using MDTraj 1.9.7 [100]. The backbone and sidechain dihedral angles (*ϕ, ψ, χ*_1_, *χ*_2_, *χ*_3_, *χ*_4_) were computed for each interface residue. Sine and cosine transformations were applied to each angle time series and the transformation yielding the maximum magnitude in correlation to *∂U* /*∂λ* was selected. For residue contacts, distances were computed between closest heavy atoms for all pairs of interface residues. For neighboring waters, water oxygens within 5 Å of any heavy atom in the mutating residue were counted. *∂U* /*∂λ* was computed (for each trajectory snapshot of all replica trajectories) using numerical differentiation. Finite difference approximation with a symmetric difference quotient was used for intermediate alchemical states and Newton’s difference quotient was used for alchemical endstates, with a step size of 1e-3 for both types of states. For a given mutation, to obtain the Pearson correlation coefficient (PCC) of each degree of freedom with respect to *∂U* /*∂λ* across all replicas, the *∂U* /*∂λ* and degree of freedom time series were separately concatenated across all replicas before computing the PCC. PCCs were computed using SciPy 1.9.0’s `pearsonr` function [116] and the 95% confidence intervals were computed by bootstrapping (number of bootstraps = 200). Each bootstrapped sample was obtained by subsampling the replica indices (with replacement) and then concatenating the time series based on the subsampled replica indices. The statistical inefficiency was computed using pymbar 3.1.1 [80]’s `statisticalInefficiency` and `statisticalInefficiencyMultiple` for individual replicas and across all replicas, respectively.

Relevance of the highest correlation degrees of freedom was assessed by inspecting the proximity of the degree of freedom to the mutation site. A blue dot was included next to each mutation whose highest correlation degree of freedom is relatively far from the mutating residue, indicating that there is not a particularly intuitive explanation for the degree of freedom’s high correlation (Figure 5, Supplementary Figure 7). Many of these mutations (with blue dots) have relatively small *∂U* /*∂λ* statistical inefficiency, which indicates that they likely do not contain significant sampling problems in the first place.

To supplement Figure 5, the same *∂U* /*∂λ* analysis was performed for 10 ns/replica AREX complex phase simulations (Supplementary Figure 7). The overall trends in sampling problems for 10 ns/replica simulations are similar to those of the 50 ns/replica simulations (Figure 5). However, since most of the statistical inefficiencies are underestimated in the 10 ns/replica plot (because many of the simulations have not yet equilibrated), the 50 ns/replica plot provides a more accurate representation of the trends in sampling problems.

### A.12 Amazon Web Services (AWS) cost calculation

The GPU time was estimated using 36 replicas and the AWS costs were estimated based on the on-demand price of an Amazon EC2 P4d instance ($32.77 per hour), which has 8 NVIDIA A100 GPUs.

## B Investigation of the discrepant ΔΔG_binding_ prediction for A29Y

With a 50 ns/replica AREX simulation, A29Y not only has a large discrepancy in ΔΔG_binding_ with respect to experiment (−2.11 kcal/mol), but also with respect to Y29A (1.45 kcal/mol) (Supplementary Figure 6C and Supplementary Table 1). We hypothesize that A29Y has poor accuracy and internal consistency because the mutant tyrosine residue does not sample its most energetically favorable orientation in the barnase:barstar interface. The mutant tyrosine residue in A29Y potentially faces more difficult sampling challenges than the wild-type tyrosine residue in Y29A because the former has to be computationally modeled onto the A29 structure, whereas the positions of the latter are taken from the crystal structure and therefore the wild-type tyrosine residue is guaranteed to be in a low energy conformation. To test our hypothesis, we monitored the distances between Y29 and three nearby residues: H102, whose sidechain stacks with Y29’s aromatic sidechain to form a hydrophobic interaction, R83, and N84, whose backbone carbonyl oxygens form hydrogen bonds with Y29’s sidechain hydroxyl oxygen [75] (Supplementary Figure 8C). We generated time series for each of the three residue pair distances at the mutant endstate (*λ* = 1, where Y29 is fully interacting with its environment) of the A29Y AREX simulation and the wild-type endstate (*λ* = 0, where Y29 is fully interacting with its environment) of the Y29A AREX simulation. We compared the distances in each time series with the crystal structure distance and found that the Y29A wild-type endstate samples the crystal structure distance for all three residue pairs (Supplementary Figure 8A), but the A29Y mutant endstate rarely samples the crystal structure distance for two of the three residue pairs (Supplementary Figure 8B-C). These findings demonstrate that even with 50 ns of simulation time, the mutant tyrosine residue does not sample the relevant orientations that would enable it to contribute favorably to the barnase:barstar interface, which explains why the predicted ΔΔG_binding_ of A29Y has poor internal consistency and accuracy. We expect that with sufficient simulation time (potentially much longer than 50 ns), the mutant tyrosine will sample the relevant orientations, eliminating the discrepancy in ΔΔG_binding_. Future work could involve improving the approach we use for computationally building in mutant residues.

## C Investigation of the discrepant ΔΔG_binding_ predictions for D35A and K27A

We investigated whether the significantly discrepant D35A and K27A predictions (with 50 ns/replica AREX) are a result of failing to account for all relevant protonation states. Since arginine and glutamine do not have alternate protonation states that are easily accessible under physiological conditions, we only examined protonation state effects for D35A and K27A. We first explored the possibility that D35 may exist in both its deprotonated (ASP) and protonated (ASH) forms. We modeled D35 as ASH and ran AREX on ASH → ALA transformations in the complex (10 ns/replica), apo (10 ns/replica), and terminally-blocked (5 ns/replica) phases. We recomputed the D35A ΔΔG_binding_, accounting for possible interconversion between the deprotonated and protonated states (see Section C.1), and found that the ΔΔG_binding_ (1.65 kcal/mol) is within error of the original, deprotonated ΔΔG_binding_ (1.66, 95% CI: [0.57, 2.75] kcal/mol). The similar ΔΔG_binding_s obtained with and without accounting for multiple protonation states indicates that our original ΔΔG_binding_ for D35A is not discrepant because of failing to incorporate all relevant protonation states. Moreover, we observed analogous results for K27A, where the ΔΔG_binding_ (accounting for multiple protonation states, 3.31 kcal/mol) is within error of the original, protonated ΔΔG_binding_ (3.32, 95% CI: [1.80, 4.84] kcal/mol), showing that protonation state effects are not causing the discrepancy in predicted ΔΔG_binding_ of K27A.

### C.1 Computation of ΔΔG_binding_s accounting for multiple protonation states

We are interested in computing the relative binding free energy, 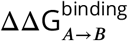, where *A* is the WT amino acid and ***B*** is the mutant amino acid, accounting for all relevant protonation states for both amino acids. We use D35A as an example, where ***A*** is aspartic acid (ASP) and ***B*** is alanine (ALA). ASP may exist in a deprotonated state (***A***) or a protonated state (***AH***), whereas ALA only has one state. To compute 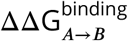, we use the thermodynamic cycles in Supplementary Figure 16.

The relative binding free energy can be defined as the difference in binding free energies between ***B*** and ***A***:

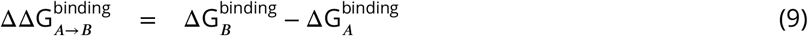

The binding free energy of chemical species *s* (e.g., *A* or *B*), accounting for multiple protonation states, can be computed as:

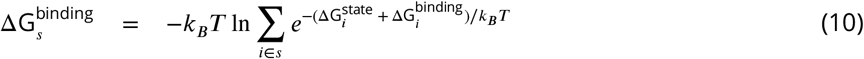

where *k*_*B*_ is the Boltzmann constant, ***T*** is temperature, *i* represents a protonation state of chemical species *s*, 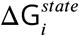 is the protonation state free energy for state *i*, and 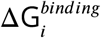 is the binding free energy at protonation state *i*. Given that the protonation state free energy can be computed as:

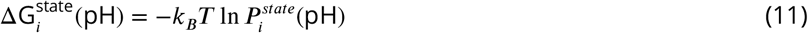

where 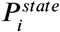 is the probability of chemical species *s* adopting protonation state *i* and pH is the pH of interest (note: we suppress the pH argument throughout the rest of the derivation), and the free energy of deprotonation can be computed as:

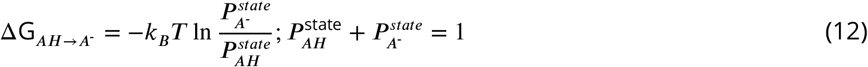

we compute the protonation state free energies as:

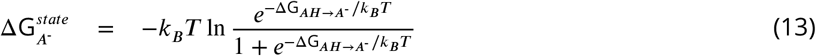

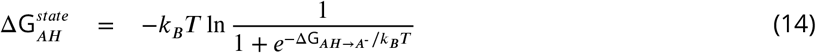

If we set *G*_*B*+*X*_ and *G*_*BX*_ to 0, we can compute the absolute binding free energies of the deprotonated and protonated states of *A* as relative free energy differences (see Supplementary Figure 16):

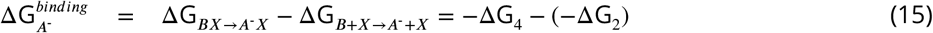

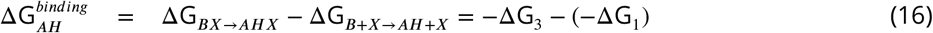

where *X* is the binding partner. We can compute 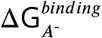 and 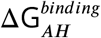 from ΔG_1_, ΔG_2_, ΔG_3_, and ΔG_4_ (Supplementary Table 2). We can also compute 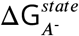 and 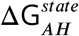 from ΔG_*AH*→*A*_ (i.e., “corrected” 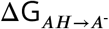 in Supplementary Table 2). We can then feed 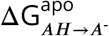 into equation 10 to compute the binding free energy of *A* (i.e., ASP), accounting for both protonation states 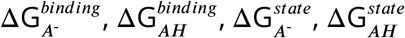. Finally, we can feed 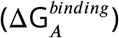 into equation 9 to compute 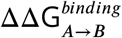. Note that since we set *G*_*B*+*X*_ and *G*_*BX*_ to 0, 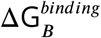 is 0. The above calculation can be repeated for K27A where *A* is LYS and *B* is ALA.

## D REST parameter selection experiments reveal that improvements in convergence are comparable across a broad range of REST parameter combinations

To run REST, the user must select the maximum effective temperature (***T***_max_), which corresponds to the highest effective temperature to which the REST region will be scaled. The user must also choose the REST region, which we define as the mutating residue and all residues within a user-specified radius of it. The higher the ***T***_max_ and the larger the radius, the more significantly the energy barriers will decrease and the more enhanced sampling will be. However, as one increases the ***T***_max_ and the radius, the thermodynamic length also increases, which increases the variance of the free energy difference (ΔG) estimates. Therefore, a key challenge when applying REST is to find the optimal combination of ***T***_max_ and radius that will decrease the correlation time of the slowest degrees of freedom while minimizing the variance of the ΔG estimate.

To explore combinations of ***T***_max_ and radius, we chose small, medium, and large values for each of the parameters. We selected 400 K, 600 K, and 1200 K for ***T***_max_ and 0.3 nm, 0.5 nm, and 0.7 nm for the radius. For each combination of parameters (9 total), we ran AREST for the complex phase of two representative mutations, A42T and R87A, and computed the discrepancy of the AREST ΔG_complex_ (at t = 10 ns) with respect to the “true” ΔG_complex_, which was computed from 100 ns/replica AREX. We used discrepancy as a metric to assess the efficiency of each REST parameter combination in achieving convergence to the true ΔG_complex_.

We compared the discrepancies across the REST parameter combination experiments and also against the reference (***T***_max_ = 300 K and no REST region) experiment. For both A42T and R87A, at the 10 ns time point, most of the REST combination ΔG_complex_s were less discrepant than the reference ΔG_complex_, indicating that AREST improves convergence more efficiently than vanilla AREX for these mutations (Supplementary Figure 9B-C). When comparing the discrepancies in ΔG_complex_s across REST combination experiments, we found that for both A42T and R87A, the discrepancies are within error of each other. The least discrepant parameter combination for both A42T and R87A is ***T***_max_ = 600 K and radius = 0.5 nm, though this combination decreases the discrepancy only slightly better than the other combinations.

We also compared the improvements of AREST over AREX between A42T and R87A. The difference in ΔG_complex_s (at t = 10 ns) for the best REST parameter combination (***T***_max_ = 600 K and radius = 0.5 nm) and the reference AREX simulation is less significant for A42T (∼0.5 kcal/mol) than it is for R87A (∼4.5 kcal/mol). Although these results initially suggest that for A42T, AREST does not significantly improve convergence compared to AREX, if we examine the difference in discrepancies at an earlier time point (2 ns instead of 10 ns), we find that the difference is greater (∼2.5 kcal/mol) than that at 10 ns (Supplementary Figure 9A-B). Therefore, AREST does improve the efficiency of ΔG_complex_ convergence for A42T, but most of the efficiency improvement occurs in the first few nanoseconds of the trajectory and afterwards, the advantages of AREST over AREX for A42T become significantly less prominent. On the other hand, for R87A, the efficiency improvement is present through at least 10 ns, perhaps even longer (Supplementary Figure 9C). Taken together, these results suggest that the same REST parameter combination can affect different mutations in the same system to varying degrees.

## E Supplementary Figures and Tables

**Supplementary Figure 1.**
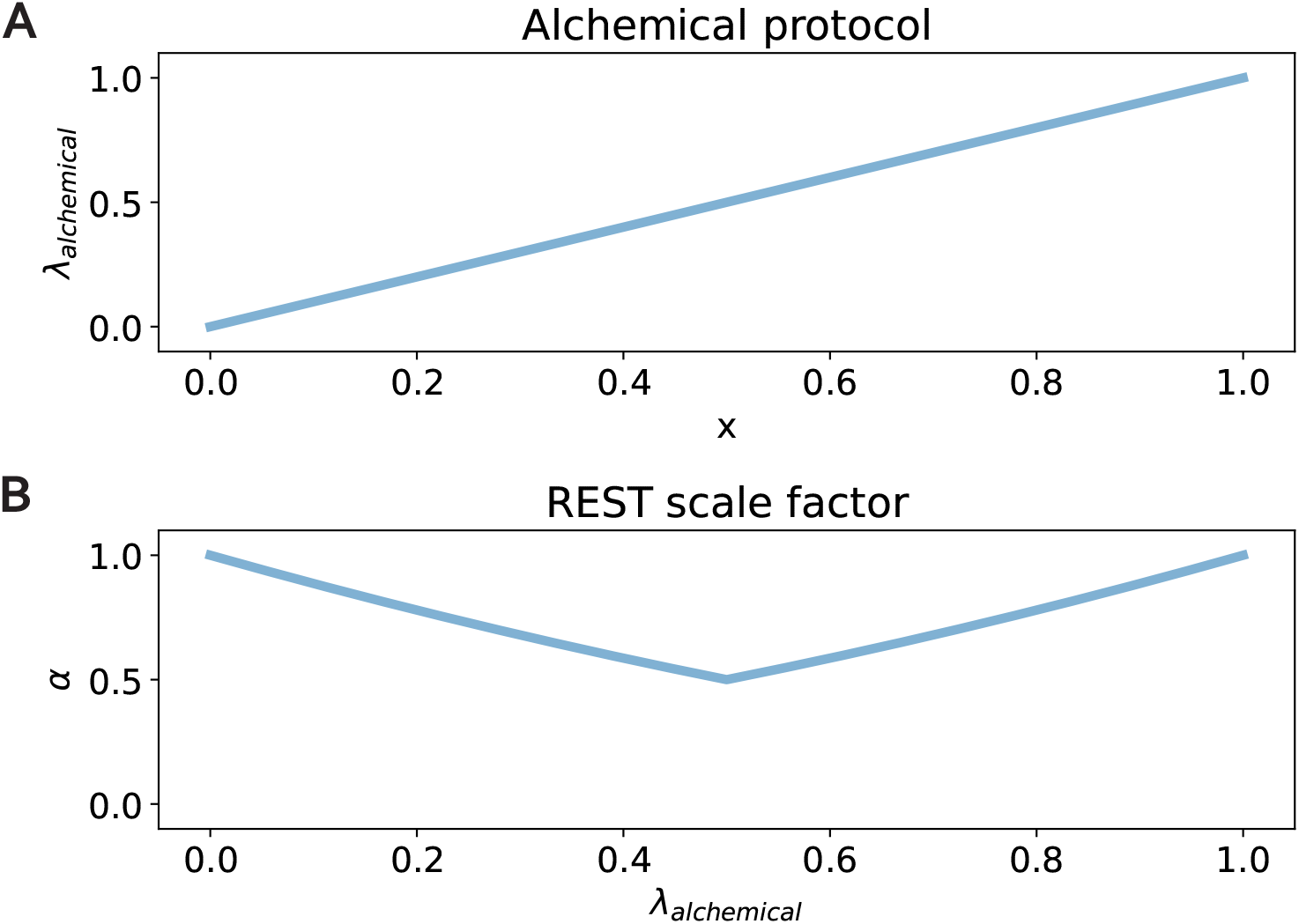
Functions for defining the alchemical protocol and REST scale factor. **(A)** The function used for defining the alchemical protocol, *λ*_alchemical_(*x*) = *x*. **(B)** The function used for defining the REST scale factor, *α*(*λ*_alchemical_, ***T***_max_, ***T***_0_), given ***T***_max_ = 600 K and ***T***_0_ = 300 K. We gradually increase the temperature from ***T***_0_ to ***T***_max_ and back down to ***T***_0_ over the alchemical protocol, reaching ***T***_max_ halfway through the protocol.

**Supplementary Figure 2.**
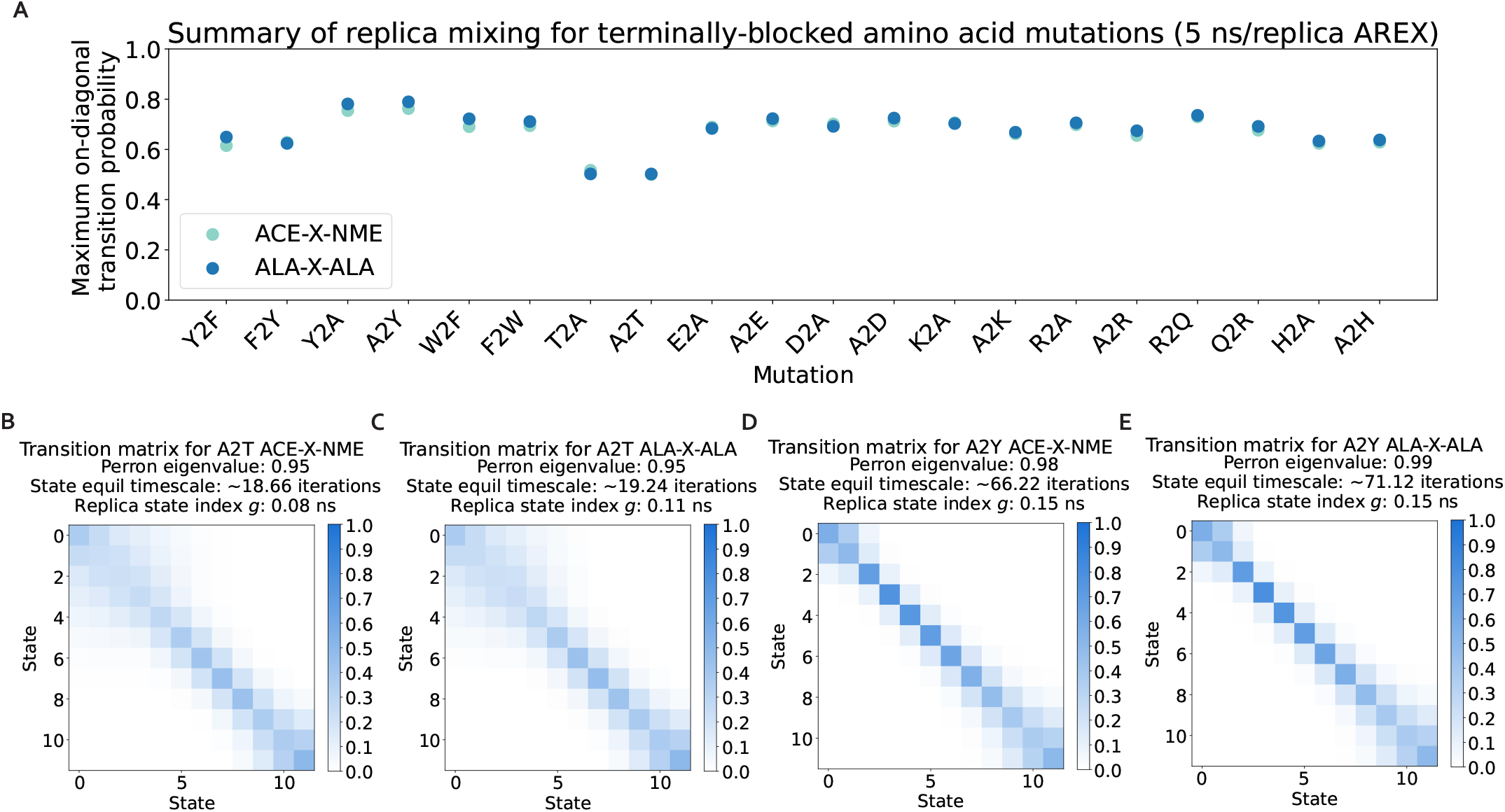
Replicas mix well for all terminally-blocked amino acid alchemical replica exchange (AREX) simulations. **(A)** Maximum on-diagonal transition probability for the transition probability matrices of each of the 20 forward and reverse terminally-blocked amino acid mutations. Transition probability matrices generated from AREX simulations (number of states = 12 and 24 for neutral and charge mutations, respectively and simulation time = 5 ns/replica). The on-diagonal transition probability quantifies the extent to which replicas are exchanging with themselves; values close to 1 indicate there is a mixing bottleneck. Light teal indicates the ACE-X-NME phase and dark blue indicates the ALA-X-ALA phase. **(B)** The transition probability matrix for the 5 ns/replica ACE-X-NME phase AREX simulation of A2T, the mutation with the minimum value in panel A. “Perron eigenvalue” corresponds to the subdominant (second) eigenvalue and measures how well the replicas have mixed, where unity indicates poor mixing due to insufficient phase space overlap between some alchemical states. “State equil timescale” corresponds to the state equilibration timescale, which is proportional to the perron eigenvalue and estimates the number of iterations elapsed before the collection of replicas fully mix once. “Replica state index *g*” corresponds to the replica state index statistical inefficiency and describes how thoroughly the replicas visit all the states (i.e., lambda windows), where a value of 0.001 ns indicates very thorough visitation of states (because the sampling interval is 0.001 ns) and large values indicate poor visitation. **(C)** The transition probability matrix for the ALA-X-ALA phase AREX simulation of A2T, the mutation with the minimum value in panel A. **(D)** The transition probability matrix for the ACE-X-NME phase AREX simulation of A2Y, the mutation with the maximum value in panel A. **(E)** The transition probability matrix for the ALA-X-ALA phase AREX simulation of A2Y, the mutation with the maximum value in panel A.

**Supplementary Figure 3.**
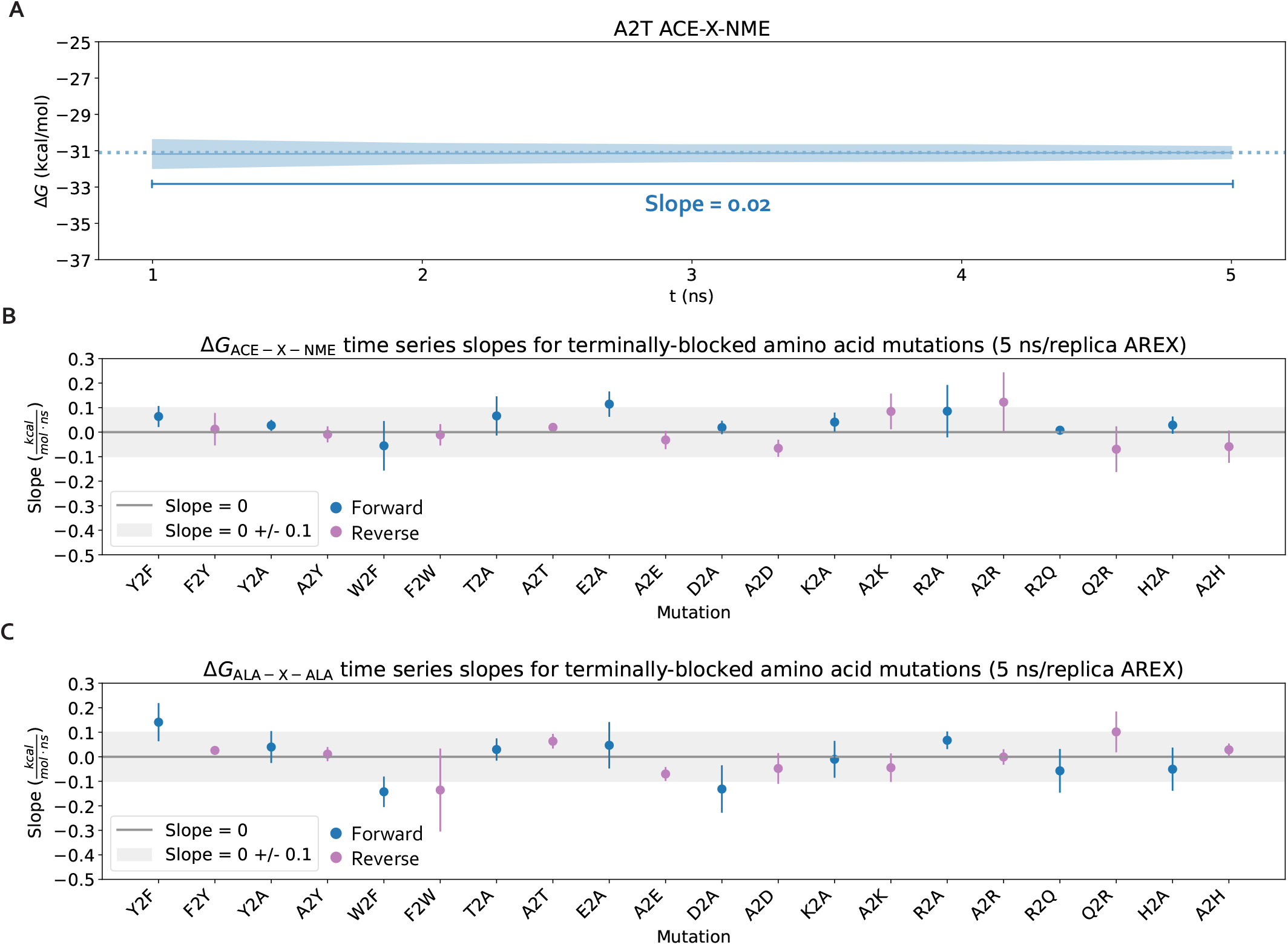
The ΔG time series flatten within 5 ns for all terminally-blocked amino acid mutations. **(A)** ΔG_ACE-X-NME_ time series for A2T, shown to illustrate the data over which the slope is computed. ΔG_ACE-X-NME_ time series was generated from an AREX simulation (number of states = 12 and simulation time = 5 ns/replica). **(B)** Slopes of the ΔG_ACE-X-NME_ time series for each mutation are shown as blue (forward mutations) and purple (reverse mutations) circles. Error bars represent two standard deviations and and were computed using the SciPy `linregress` function. Slopes within error of the shaded gray region (0 ± 0.1 kcal/mol/ns) are close to 0 and are therefore considered “flat.” **(C)** Same as (B), but for ALA-X-ALA phase instead of ACE-X-NME phase.

**Supplementary Figure 4.**
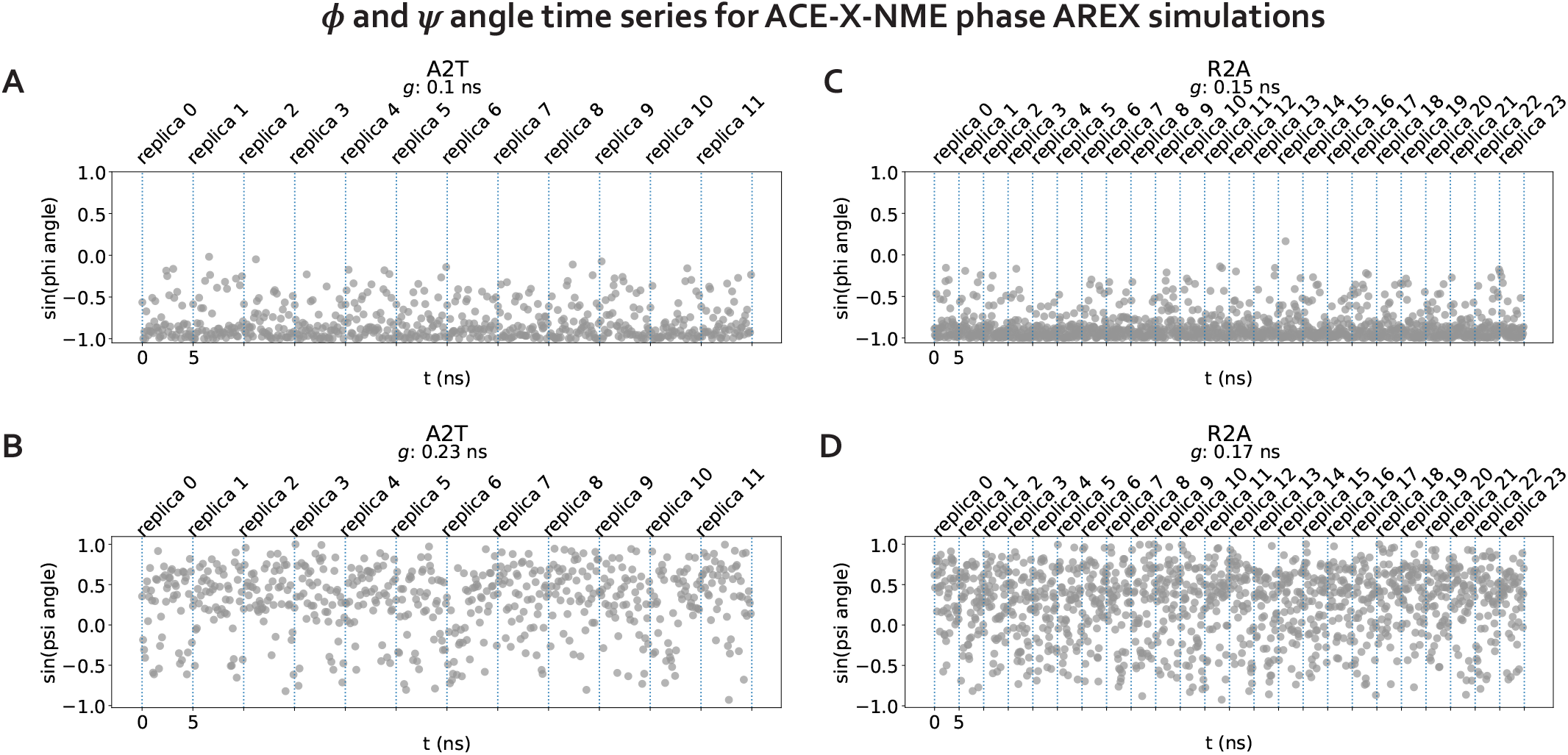
The *ϕ* and *ψ* angles of two representative terminally-blocked amino acid mutations are sampled sufficiently. **(A)** *ϕ* angle time series for each replica of the A2T ACE-X-NME phase AREX simulation (number of states = 12, simulation time = 5 ns/replica). Dotted blue lines separate each replica time series. *g* indicates statistical inefficiency, which was computed from time series with a sampling interval of 0.1 ns. **(B)** Same as (A), but for A2T *ψ* angle instead of A2T *ϕ* angle. **(C)** *ϕ* angle time series for each replica of the R2A ACE-X-NME phase AREX simulation (number of states = 24, simulation time = 5 ns/replica). **(D)** Same as (C), but for the R2A *ψ* angle instead of the R2A *ϕ* angle.

**Supplementary Figure 5.**
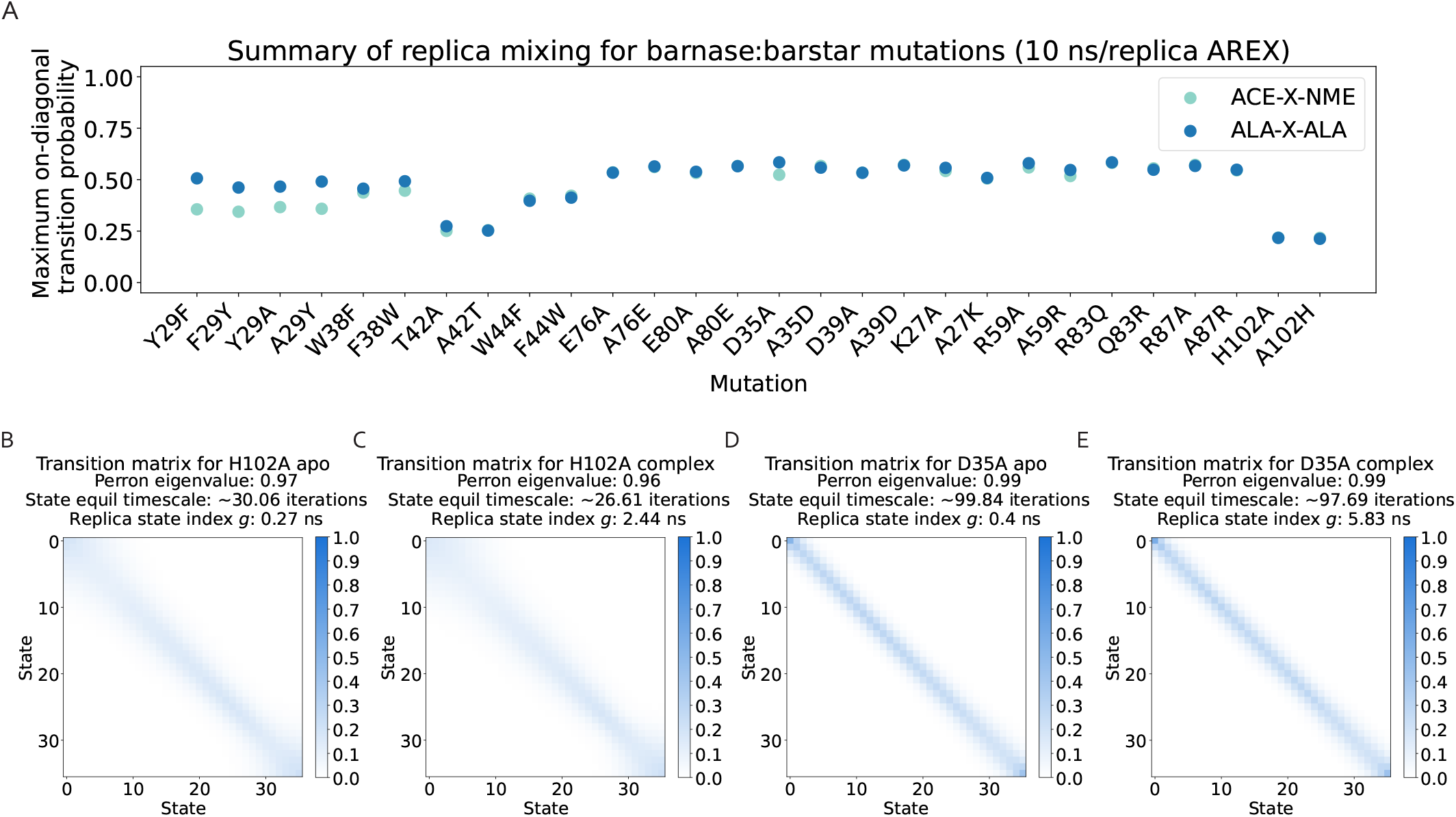
Replicas mix well for all barnase:barstar alchemical replica exchange (AREX) simulations. **(A)** Maximum on-diagonal transition probability for the transition probability matrices of each of the 28 forward and reverse barnase:barstar mutations. Transition probability matrices generated from AREX simulations (number of states = 24 and 36 for neutral and charge mutations, respectively and simulation time = 10 ns/replica). The on-diagonal transition probability quantifies the extent to which replicas are exchanging with themselves; values close to 1 indicate there is a mixing bottleneck. Light teal indicates the apo phase and dark blue indicates the complex phase. **(B)** The transition probability matrix for the 10 ns/replica apo phase AREX simulation of H102A, the mutation with the minimum value in panel A. “Perron eigenvalue” corresponds to the subdominant (second) eigenvalue and measures how well the replicas have mixed, where unity indicates poor mixing due to insufficient phase space overlap between some alchemical states. “State equil timescale” corresponds to the state equilibration timescale, which is proportional to the perron eigenvalue and estimates the number of iterations elapsed before the collection of replicas fully mix once. “Replica state index *g*” corresponds to the replica state index statistical inefficiency and describes how thoroughly the replicas visit all the states (i.e., lambda windows), where a value of 0.001 ns indicates very thorough visitation of states (because the sampling interval is 0.001 ns) and large values indicate poor visitation. **(C)** The transition probability matrix for the complex phase AREX simulation of H102A, the mutation with the minimum value in panel A. **(D)** The transition probability matrix for the apo phase AREX simulation of D35A, the mutation with the maximum value in panel A. **(E)** The transition probability matrix for the complex phase AREX simulation of D35A, the mutation with the maximum value in panel A.

**Supplementary Figure 6.**
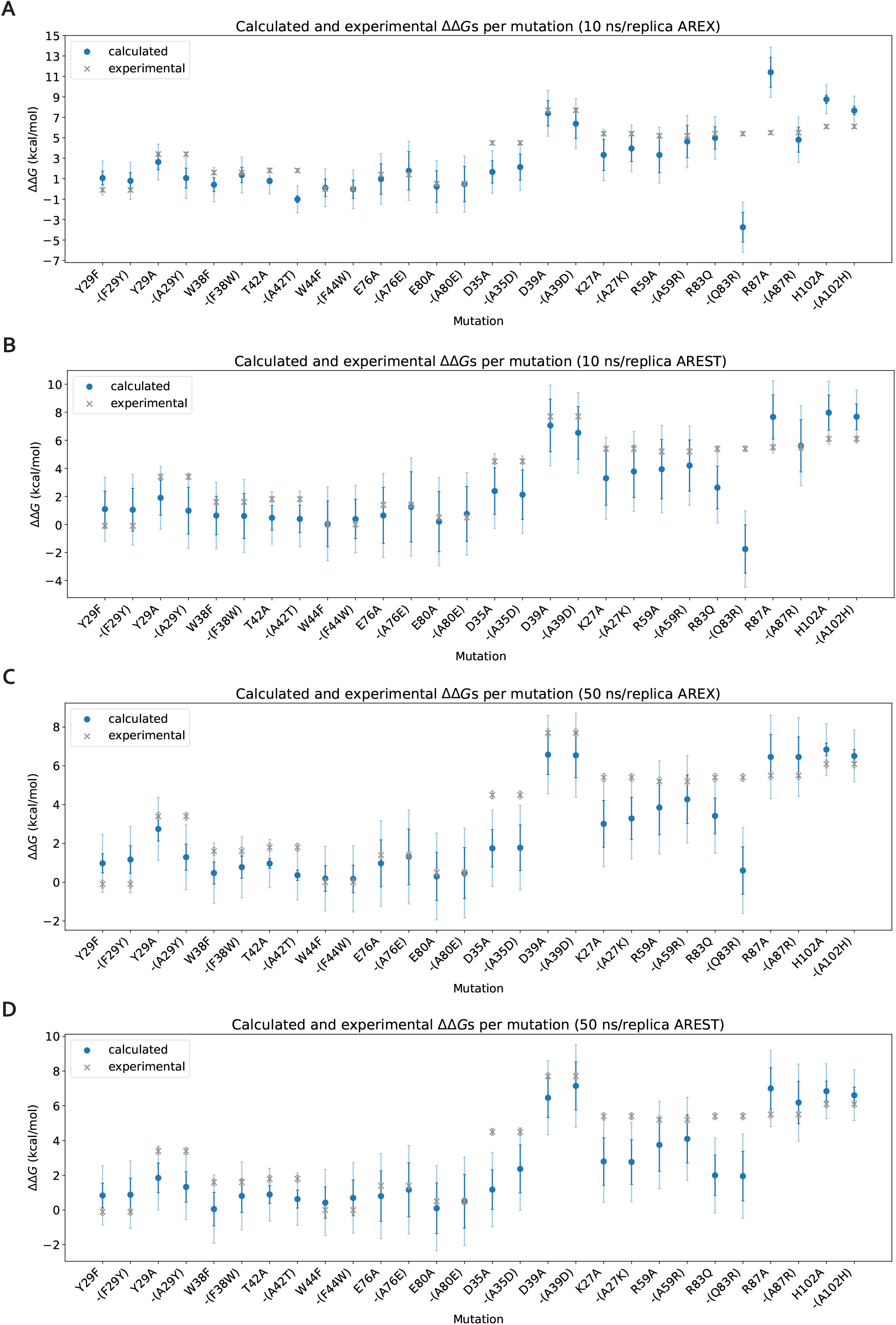
Calculated ΔΔGs for barnase:barstar mutations show decent agreement with experimental ΔΔGs using 10 ns/replica simulations and improved agreement using 50 ns/replica simulations. The data in this figure is the same data as in Figure 6B, D, F and Supplementary Figure 10B and is shown here in an alternate representation for clarity. **(A)** Calculated and experimental ΔΔGs per mutation for 10 ns/replica alchemical replica exchange (AREX). Dark blue error bars represent two standard deviations and were computed by bootstrapping the decorrelated reduced potential matrices 200 times. Light blue error bars represent two standard deviations ± 1 kcal/mol, shown to help determine whether the calculated ΔΔG is within 1 kcal/mol of experimental ΔΔG. Gray error bars indicate two standard deviations and were taken from Schreiber et al. [73] **(B)** Same as (A) but for 10 ns/replica AREST. **(C)** Same as (A) but for 50 ns/replica AREX. **(D)** Same as (A) but for 50 ns/replica AREST.

**Supplementary Figure 7.**
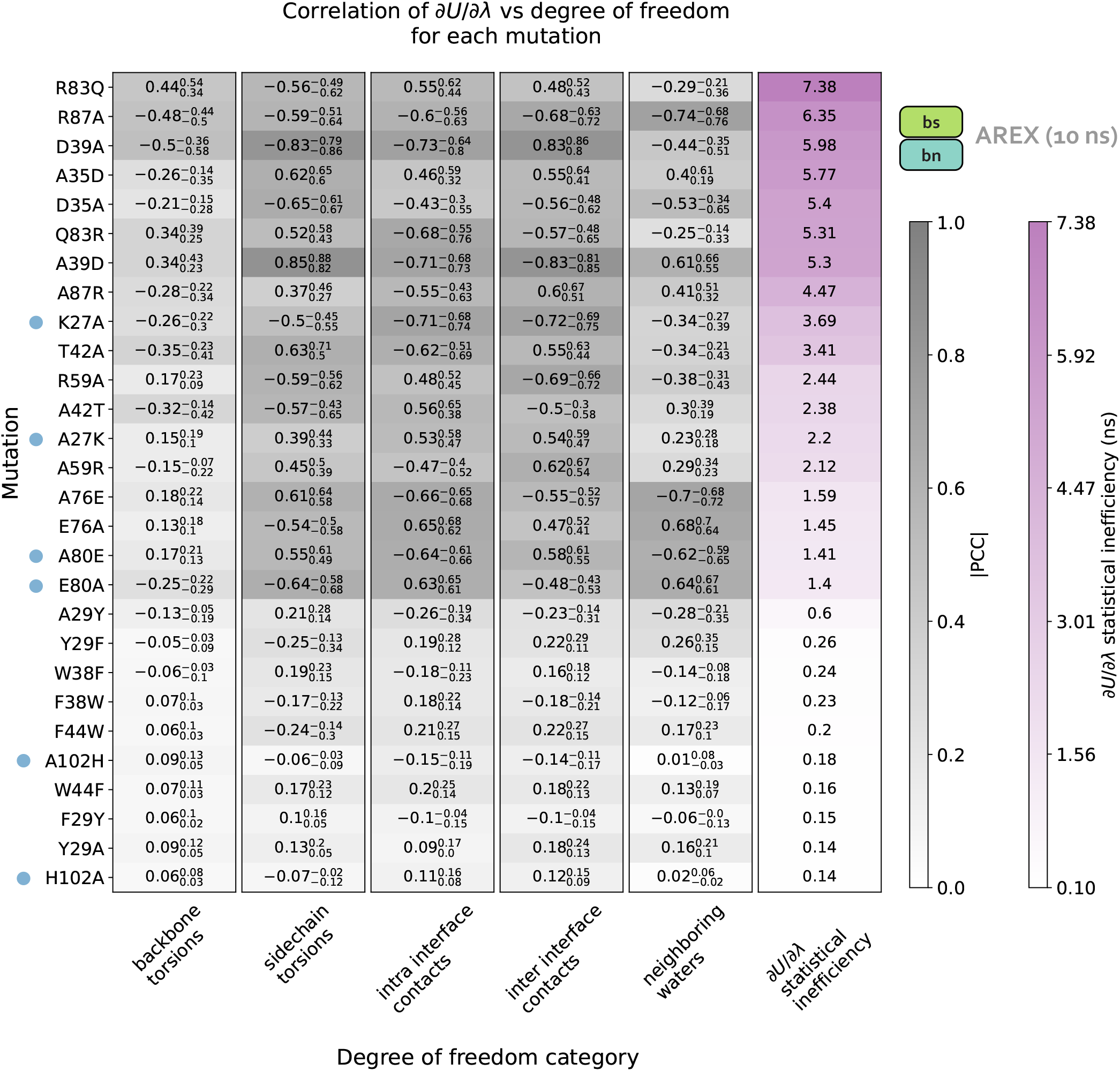
*∂U* /*∂λ* correlation analysis for 10 ns/replica AREX complex phase simulations. For details on how to interpret this plot, see caption for Figure 5.

**Supplementary Figure 8.**
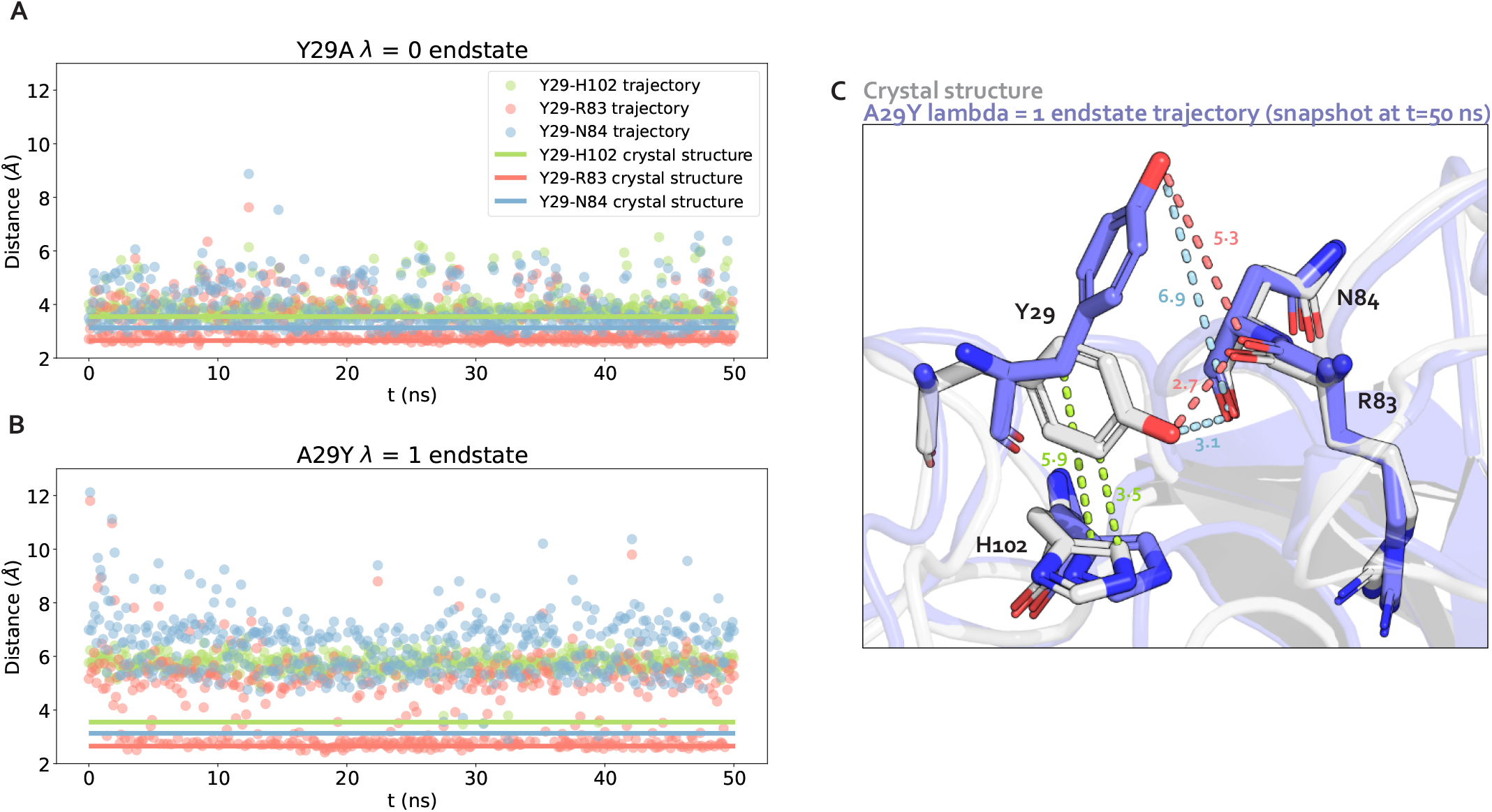
During a 50 ns/replica AREX simulation, the mutant tyrosine residue of A29Y rarely finds its most energetically favorable orientation in the barnase:barstar interface. **(A)** Distance time series for three residue pairs: Y29-H102 (green), Y29-R83 (pink), and Y29-N84 (blue) in the Y29A *λ* = 0 endstate trajectory (simulation time between frames: 100 ps, total simulation time: 50 ns). Horizontal lines represent the crystal structure (PDB ID: 1BRS) distance for each residue pair. **(B)** Same as (A) but for the A29Y *λ* = 1 endstate trajectory instead of the Y29A *λ* = 0 endstate trajectory. **(C)** Structural representation of Y29-H102, Y29-R83, and Y29-N84 residue pairs for the crystal structure (light gray) and the last snapshot (t = 50 ns) of the A29Y *λ* = 1 endstate trajectory (purple). Distances (in Å) between Y29-H102 (green), Y29-R83 (pink), and Y29-N84 (blue) shown as dotted lines. Nitrogen atoms in dark blue and oxygen atoms in red.

**Supplementary Figure 9.**
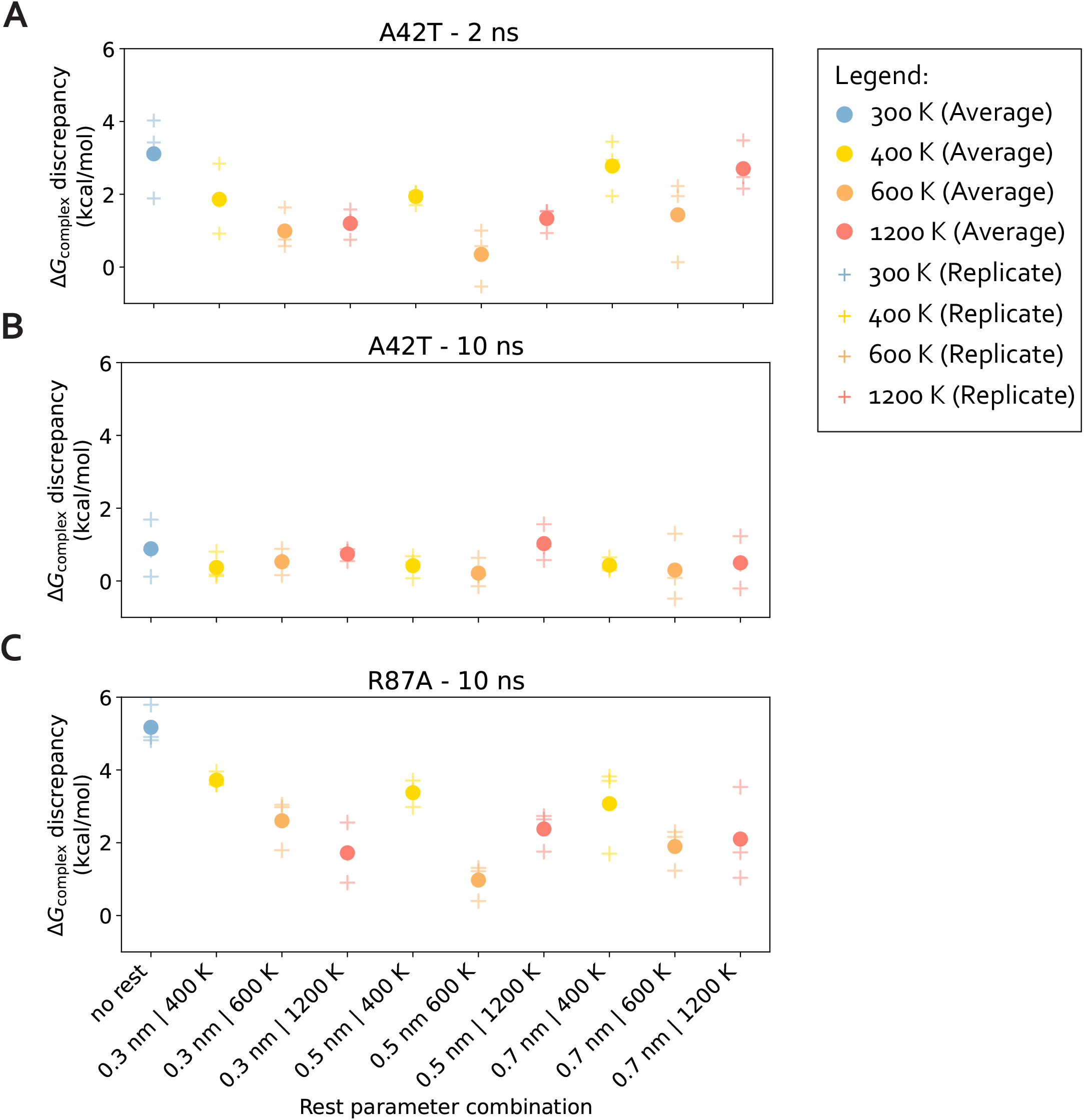
The REST parameter combinations show comparable convergence, but 0.5 nm and 600 K is (marginally) the best for both A42T and R87A. **(A)** Comparison of different combinations of two REST parameters: maximum temperature (***T***_max_) and radius. For each combination, the discrepancy of the complex phase AREST free energy difference at 2 ns with respect to the “true” free energy difference 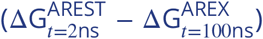 was computed. Blue markers represent the case where no REST was used (i.e., ***T***_max_ = 300 K), yellow markers represent ***T***_max_ = 400 K, orange markers represent ***T***_max_ = 600 K, and red markers represent ***T***_max_ = 1200 K. Circles represent the mean discrepancy across 3 replicates and plus signs represent the discrepancy for each individual replicate. **(B)** Same as (A), but with the discrepancy computed at 10 ns instead of 2 ns. **(C)** Same as (A), but using R87A instead of A42T and with the discrepancy computed at 10 ns instead of 2 ns.

**Supplementary Figure 10.**
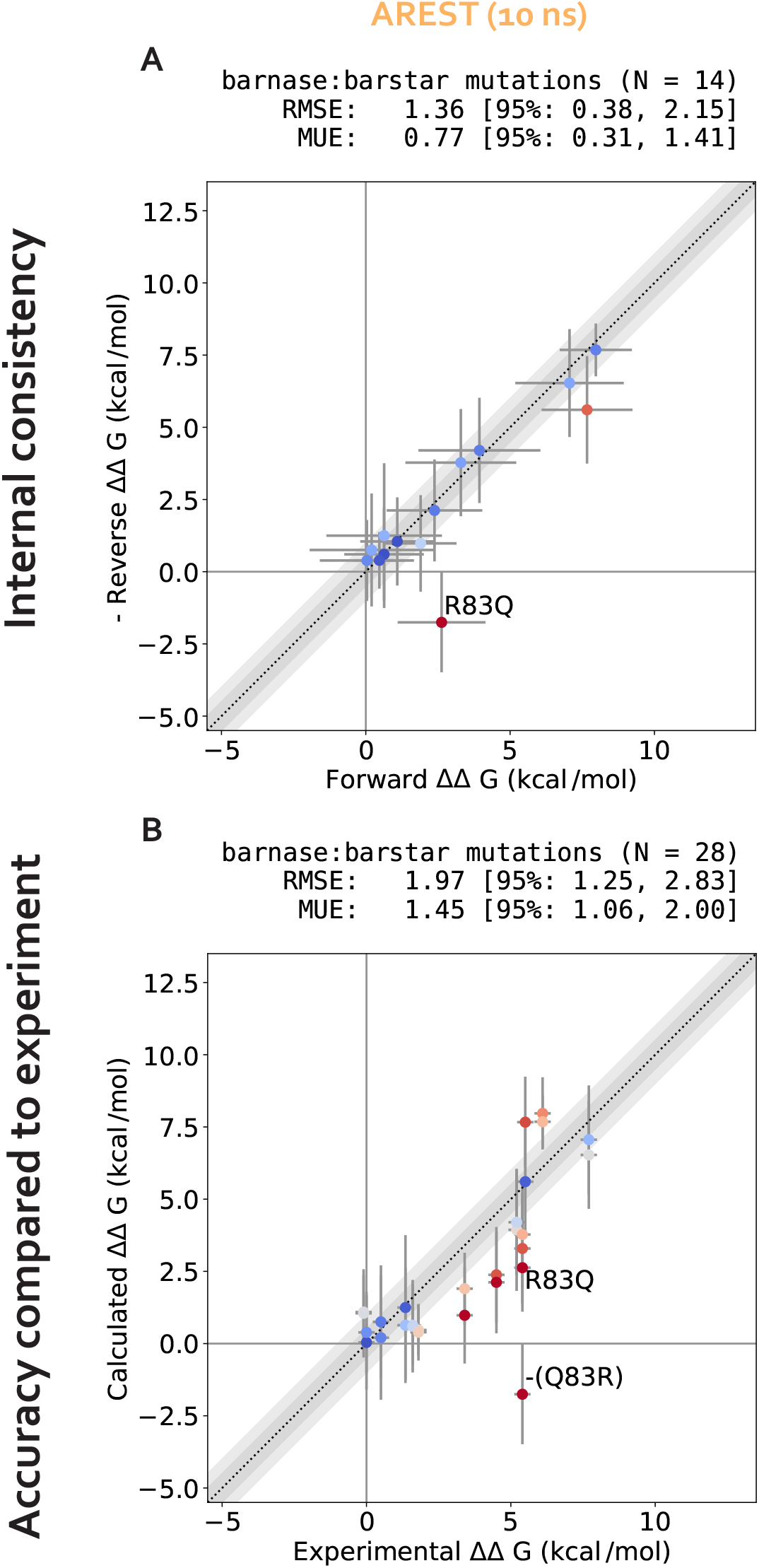
Internal consistency and accuracy for ΔΔG_binding_s from 10 ns/replica simulations of AREST. **(A)** (Negative of the) Reverse versus forward ΔΔG_binding_s for each barnase:barstar mutation computed from AREST simulations (number of states = 24 and 36 for neutral and charge mutations, respectively and simulation time = 10 ns/replica for each phase). **(B)** Calculated versus experimental ΔΔG_binding_s for each barnase:barstar mutation computed from AREST simulations (number of states = 24 and 36 for neutral and charge mutations, respectively and simulation time = 10 ns/replica for each phase). For details on how to interpret these plots, see caption for Figure 6.

**Supplementary Figure 11.**
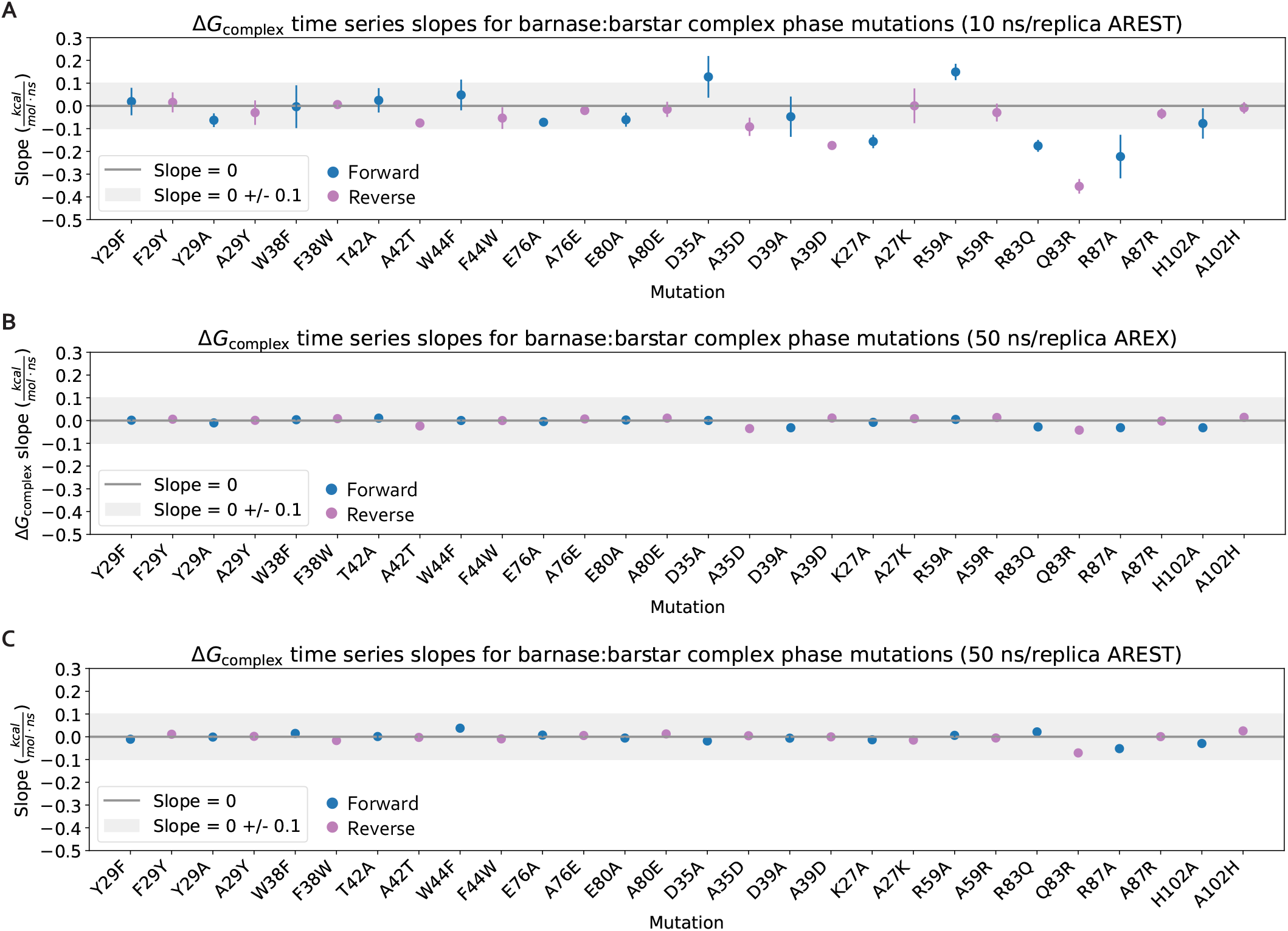
The ΔG_complex_ time series converge with long alchemical replica exchange (AREX) and alchemical replica exchange with solute tempering (AREST) simulations. **(A)** Slopes of the last 5 ns of the ΔG_complex_ time series for each barnase:barstar mutation are shown as blue (forward mutations) and purple (reverse mutations) circles. ΔG_complex_ time series were generated from 10 ns/replica complex phase AREST simulations (number of states = 24 and 36 for neutral and charge mutations, respectively). Error bars represent 2 standard deviations and were computed using the SciPy `linregress` function. Slopes within error of the shaded gray region (0 ± 0.1 kcal/mol/ns) are close to 0 and are therefore considered “flat.” **(B)** Same as (A), but for 50 ns/replica AREX complex phase simulations (number of states = 24 and 36 for neutral and charge mutations, respectively) instead of 10 ns/replica AREST complex phase simulations. **(C)** Same as (A), but for 50 ns/replica AREST complex phase simulations (number of states = 24 and 36 for neutral and charge mutations, respectively) instead of 10 ns/replica AREST complex phase simulations.

**Supplementary Figure 12.**
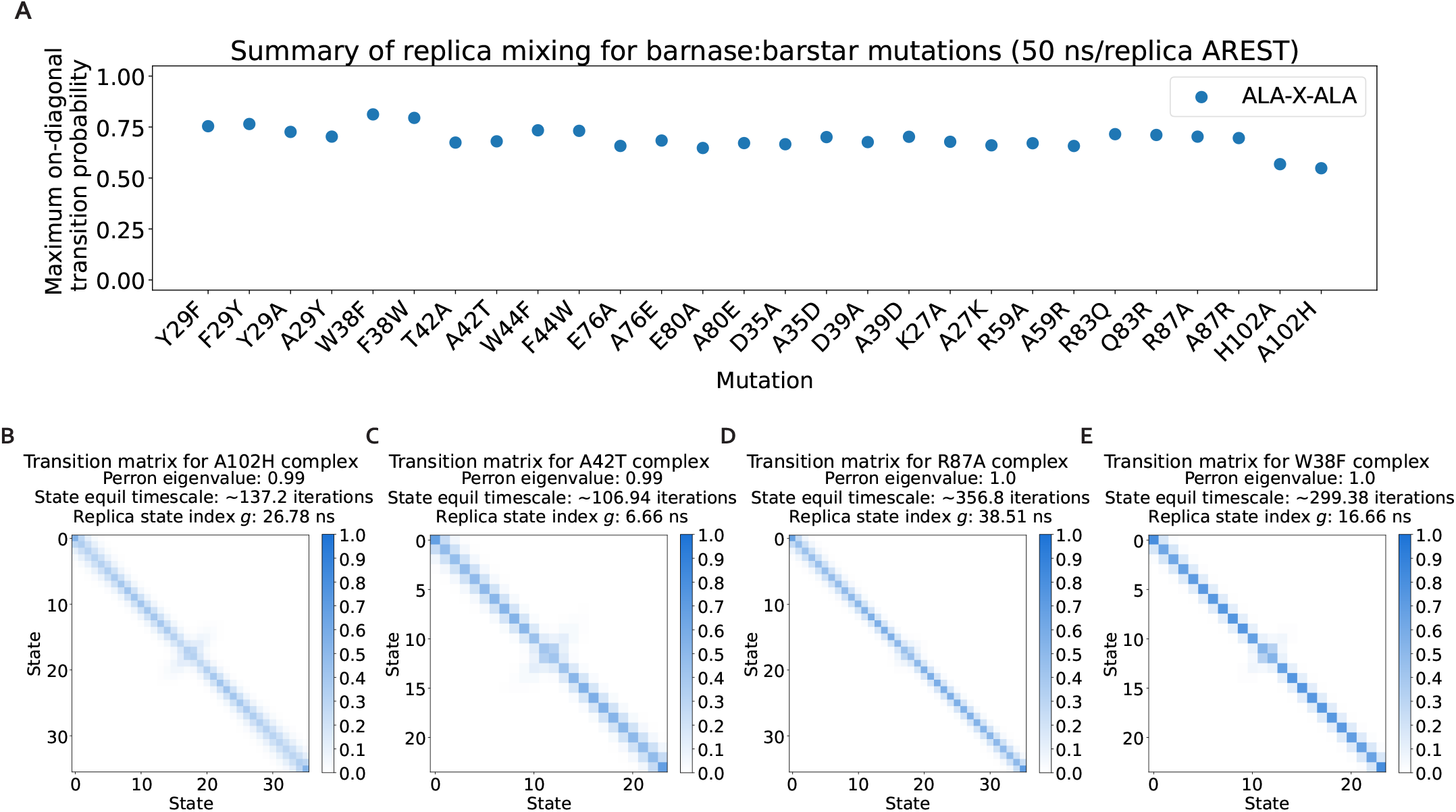
Replica mixing is sufficient for all barnase:barstar alchemical replica exchange with solute tempering (AREST) simulations. **(A)** Maximum on-diagonal transition probability for the transition probability matrices of each of the 28 forward and reverse barnase:barstar mutations. Transition probability matrices generated from complex phase AREST simulations (number of states = 24 and 36 for neutral and charge mutations, respectively and simulation time = 50 ns/replica). The on-diagonal transition probability quantifies the extent to which replicas are exchanging with themselves; values close to 1 indicate there is a mixing bottleneck. **(B)** The transition probability matrix for the 50 ns/replica complex phase AREST simulation of A102H, the mutation with the minimum value in panel A. “Perron eigenvalue” corresponds to the subdominant (second) eigenvalue and measures how well the replicas have mixed, where unity indicates poor mixing due to insufficient phase space overlap between some alchemical states. “State equil timescale” corresponds to the state equilibration timescale, which is proportional to the perron eigenvalue and estimates the number of iterations elapsed before the collection of replicas fully mix once. “Replica state index *g*” corresponds to the replica state index statistical inefficiency and describes how thoroughly the replicas visit all the states (i.e., lambda windows), where a value of 0.001 ns indicates very thorough visitation of states (because the sampling interval is 0.001 ns) and large values indicate poor visitation. **(C)** The transition probability matrix for the complex phase AREST simulation of A42T, a mutation with a maximum on-diagonal transition probability close to the mean in panel A. **(D)** The transition probability matrix for the complex phase AREST simulation of R87A, a mutation with a maximum on-diagonal transition probability close to the mean in panel A. **(E)** The transition probability matrix for the complex phase AREST simulation of W38F, the mutation with the maximum value in panel A.

**Supplementary Figure 13.**
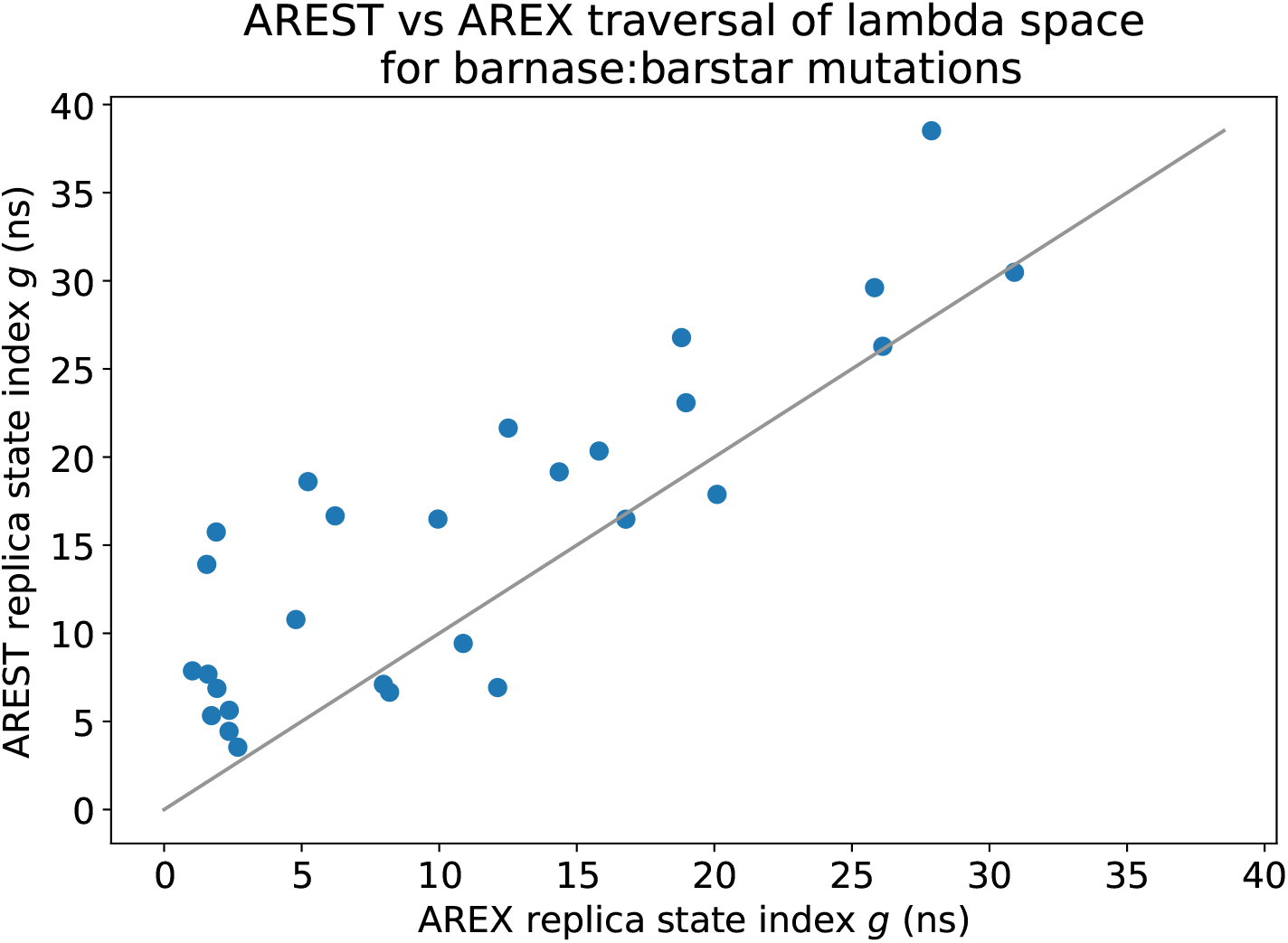
AREST traversal of lambda space is slightly worse than that of AREX. AREST versus AREX replica state index statistical inefficiencies (*g*) for the complex phase simulations of each barnase:barstar mutation (number of states = 24 and 36 for neutral and charge mutations, respectively and simulation time = 50 ns/replica). Replica state index statistical inefficiency describes how thoroughly the replicas visit all the states (i.e., lambda windows), where a value of 0.001 ns indicates very thorough visitation of states (because the sampling interval is 0.001 ns) and large values indicate poor visitation. The y = x (gray) line represents zero discrepancy between AREX and AREST statistical inefficiencies.

**Supplementary Figure 14.**
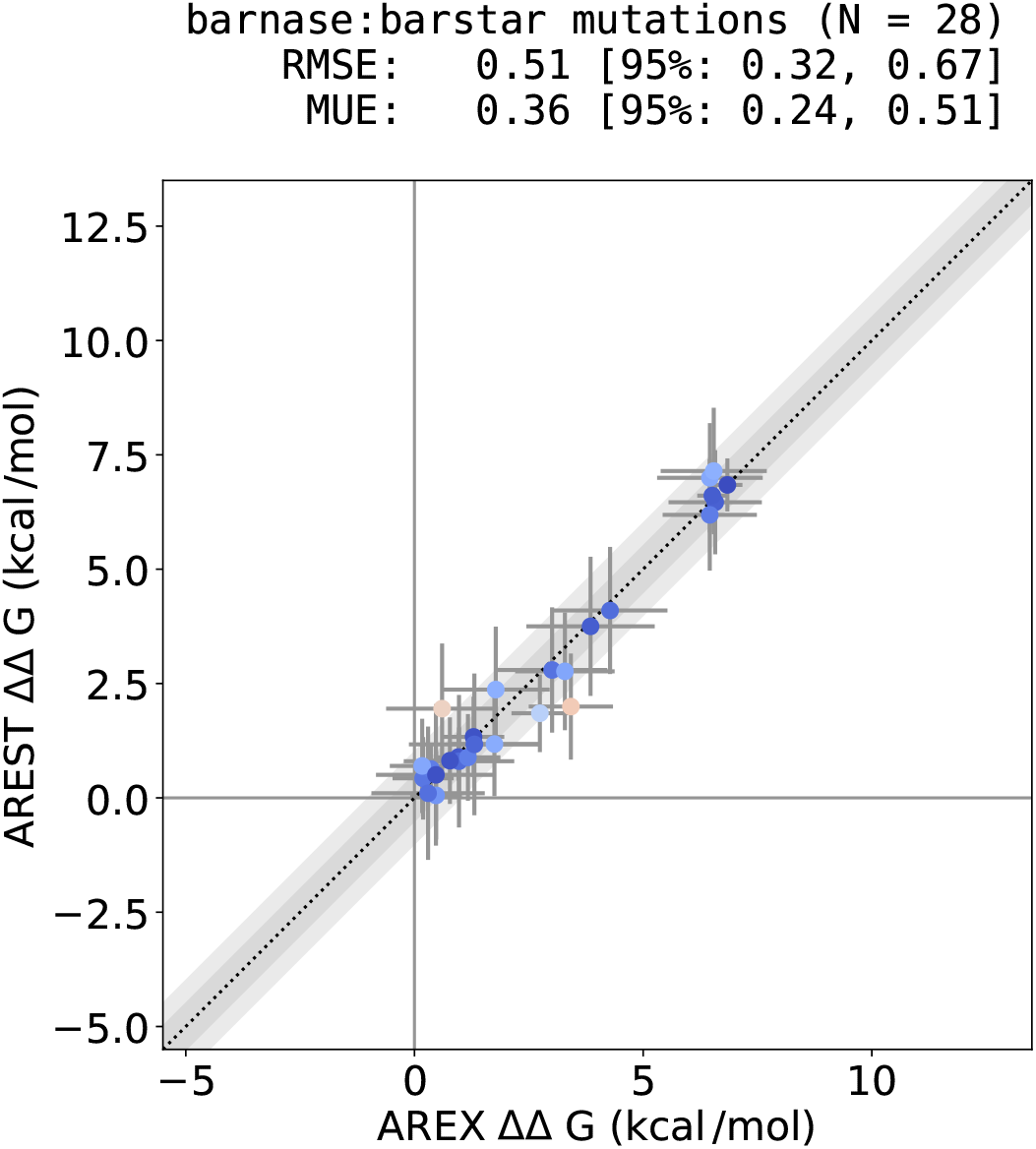
AREX and AREST ΔΔG_binding_s demonstrate good agreement. AREST versus AREX ΔΔG_binding_s for each barnase:barstar mutation (number of states = 24 and 36 for neutral and charge mutations, respectively and simulation time = 10 ns/replica for apo, 50 ns/replica for complex). The y = x (black dotted) line represents zero discrepancy between AREX and AREST ΔΔG_binding_s, the dark gray shaded region represents 0.5 kcal/mol discrepancy, and the light gray region represents 1 kcal/mol discrepancy. Data points are colored by how far they are from zero discrepancy (dark blue and red indicate close to and far from zero, respectively). Error bars represent two standard deviations and were computed by bootstrapping the decorrelated reduced potential matrices 200 times. Root mean square error (RMSE) and mean unsigned error (MUE) are shown with 95% confidence intervals obtained from bootstrapping the data 1000 times.

**Supplementary Figure 15.**
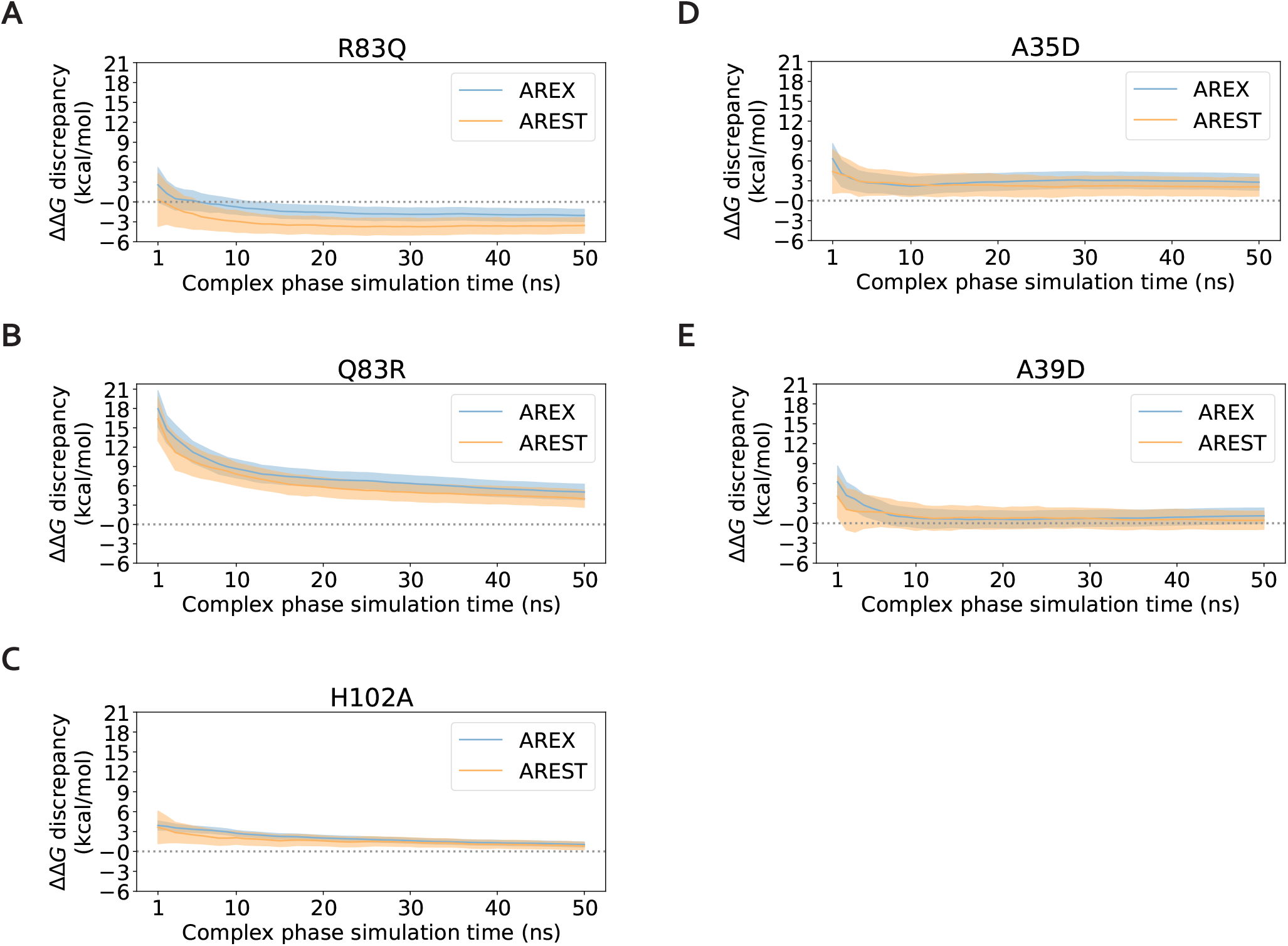
With 50 ns/replica simulation time, AREST does not significantly improve convergence over AREX for most barnase:barstar mutations with slow ΔG_complex_ convergence. **(A)-(E)**: ΔΔG discrepancy (with respect to experiment) time series for significantly discrepant mutations R83Q, Q83R, H102A, A35D, and A39D. The discrepancy was computed as ΔG_complex_ − ΔG_apo_ − ΔΔG_experiment_, where ΔG_complex_ corresponds to the (AREX or AREST) complex phase ΔG at a particular time point, ΔG_apo_ corresponds to the apo phase ΔG computed from a 10 ns/replica AREX simulation, and ΔΔG_experiment_ is the experimental value from Schreiber et al [73]. Alchemical replica exchange (AREX) time series shown in blue and alchemical replica exchange with solute tempering (AREST, with radius = 0.5 nm, ***T***_max_ = 600 K) time series shown in orange. For AREX and AREST simulations, number of states = 24 and 36 for neutral and charge-changing mutations, respectively, and simulation time = 50 ns/replica. Shaded regions represent ± two standard deviations. Gray dashed line indicates ΔΔG discrepancy = 0.

**Supplementary Figure 16.**
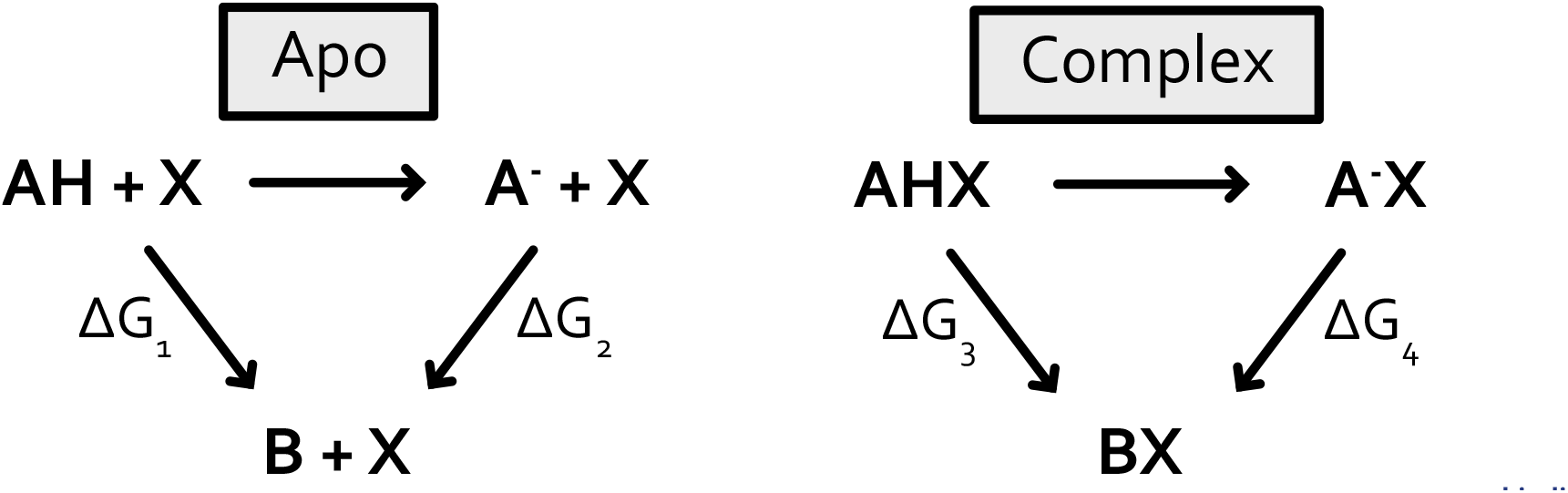
Thermodynamic cycles for computing the relative binding free energy, 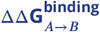, accounting for multiple protonation states. A^-^ represents the deprotonated form of the WT amino acid, AH represents the protonated form of the WT amino acid, B represents the mutant amino acid, X represents the binding partner.

**Supplementary Table 1.** ΔΔG_binding_s for barnase:barstar mutations computed from AREX simulations (with 50 ns/replica and 10 ns/replica simulation time for complex and apo phases, respectively). “Error” corresponds to one standard deviation and was computed by bootstrapping the decorrelated reduced potential matrices 200 times. The last row corresponds to the ΔΔG_binding_ for Q83R where the complex phase simulation time was 100 ns/replica. CSV file available at https://github.com/choderalab/perses-barnase-barstar-paper/blob/main/data/table_50ns_arex.csv.

| Mutation | Predicted $\Delta\Delta G$ (kcal/mol) | Error (kcal/mol) | Experimental $\Delta\Delta G$ (kcal/mol) | Complex phase simulation time (ns/replica) | Apo phase simulation time (ns/replica) | Mutation direction |
| --- | --- | --- | --- | --- | --- | --- |
| Y29F | 0.97 | 0.25 | -0.1 | 50 | 10 | forward |
| Y29A | 2.74 | 0.31 | 3.4 | 50 | 10 | forward |
| W38F | 0.47 | 0.28 | 1.6 | 50 | 10 | forward |
| T42A | 0.96 | 0.12 | 1.8 | 50 | 10 | forward |
| W44F | 0.19 | 0.33 | 0 | 50 | 10 | forward |
| E76A | 0.97 | 0.6 | 1.4 | 50 | 10 | forward |
| E80A | 0.3 | 0.62 | 0.5 | 50 | 10 | forward |
| D35A | 1.75 | 0.48 | 4.5 | 50 | 10 | forward |
| D39A | 6.58 | 0.51 | 7.7 | 50 | 10 | forward |
| K27A | 3.01 | 0.6 | 5.4 | 50 | 10 | forward |
| R59A | 3.85 | 0.7 | 5.2 | 50 | 10 | forward |
| R83Q | 3.42 | 0.46 | 5.4 | 50 | 10 | forward |
| R87A | 6.46 | 0.58 | 5.5 | 50 | 10 | forward |
| H102A | 6.84 | 0.16 | 6.1 | 50 | 10 | forward |
| F29Y | -1.17 | 0.36 | 0.1 | 50 | 10 | reverse |
| A29Y | -1.29 | 0.34 | -3.4 | 50 | 10 | reverse |
| F38W | -0.78 | 0.28 | -1.6 | 50 | 10 | reverse |
| A42T | -0.36 | 0.13 | -1.8 | 50 | 10 | reverse |
| F44W | -0.17 | 0.35 | 0 | 50 | 10 | reverse |
| A76E | -1.31 | 0.72 | -1.4 | 50 | 10 | reverse |
| A80E | -0.47 | 0.66 | -0.5 | 50 | 10 | reverse |
| A35D | -1.78 | 0.59 | -4.5 | 50 | 10 | reverse |
| A39D | -6.54 | 0.58 | -7.7 | 50 | 10 | reverse |
| A27K | -3.29 | 0.54 | -5.4 | 50 | 10 | reverse |
| A59R | -4.28 | 0.63 | -5.2 | 50 | 10 | reverse |
| Q83R | -0.6 | 0.61 | -5.4 | 50 | 10 | reverse |
| A87R | -6.46 | 0.51 | -5.5 | 50 | 10 | reverse |
| A102H | -6.51 | 0.16 | -6.1 | 50 | 10 | reverse |
| Q83R | -2.76 | 0.59 | -5.4 | 100 | 10 | reverse |

**Supplementary Table 2.**
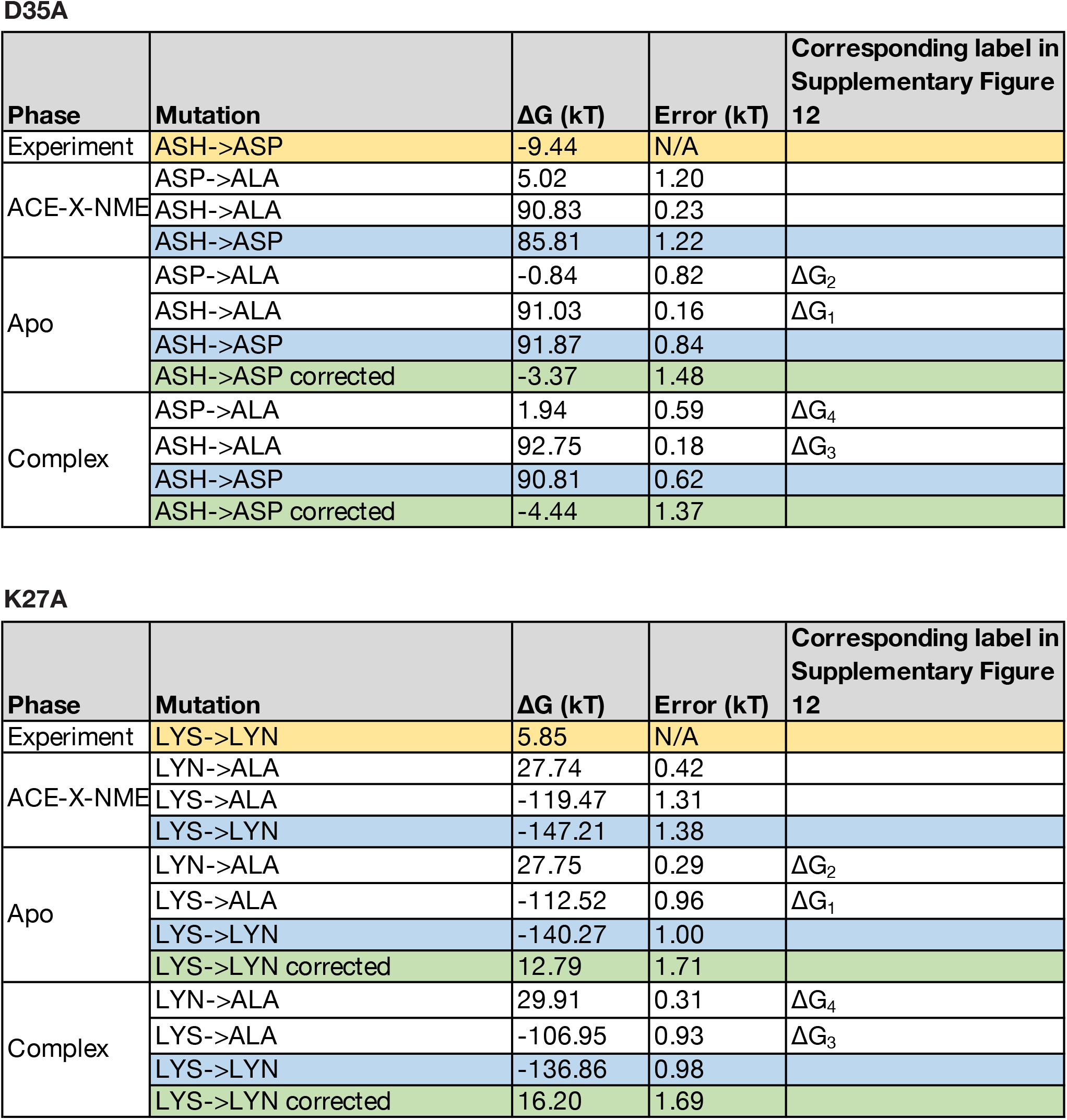
ΔGs for computation of ΔΔG_binding_s (accounting for multiple protonation states) for D35A and K27A. The “experiment” ΔGs are shown in yellow and were computed as ΔG = *k*_*B*_ ***T*** (pK_a_ − pH) ln 10 using a pH of 8.0 [73]. The ΔGs for mutations to alanine are shown in white and were computed from 5 ns/replica ACE-X-NME, 10 ns/replica apo, or 10 ns/replica complex phase simulations. The ΔGs for deprotonation (*AH* → *A*^-^) are shown in blue and were computed by subtracting pairs of ΔGs for mutations to alanine (white rows). The “corrected” ΔG_*AH*→*A*_s are shown in green and were computed according to equation 1 of Mongan et al [117] (using the ΔG_*AH*→*A*_s in the yellow and blue rows in the table). “Error” corresponds to one standard deviation and was computed by bootstrapping the decorrelated reduced potential matrices 200 times (white rows). The bootstrapped uncertainties were propagated for the blue and green rows. The XLSX file is available at https://github.com/choderalab/perses-barnase-barstar-paper/blob/main/data/D35A_K27A.xlsx.

